# Rapid active zone remodeling consolidates presynaptic potentiation

**DOI:** 10.1101/493452

**Authors:** Mathias A. Böhme, Anthony W. McCarthy, Andreas T. Grasskamp, Christine B. Beuschel, Pragya Goel, Meida Jusyte, Desiree Laber, Sheng Huang, Ulises Rey, Astrid G. Petzold, Martin Lehmann, Fabian Göttfert, Pejmun Haghighi, Stefan W. Hell, David Owald, Dion Dickman, Stephan J. Sigrist, Alexander M. Walter

## Abstract

Synaptic transmission is mediated by neurotransmitter release at presynaptic active zones (AZs) followed by postsynaptic neurotransmitter detection. Plastic changes in transmission maintain functionality during perturbations and enable memory formation. Postsynaptic plasticity targets neurotransmitter receptors, but presynaptic plasticity mechanisms directly regulating the neurotransmitter release apparatus remain largely enigmatic. Here we describe that AZs consist of nano-modular release site units and identify a molecular sequence adding more modules within minutes of plasticity induction. This requires cognate transport machinery and a discrete subset of AZ scaffold proteins. Structural remodeling is not required for the immediate potentiation of neurotransmitter release, but rather necessary to sustain this potentiation over longer timescales. Finally, mutations in Unc13 that disrupt homeostatic plasticity at the neuromuscular junction also impair shot-term memory when central neurons are targeted, suggesting that both forms of plasticity operate via Unc13. Together, while immediate synaptic potentiation capitalizes on available material, it triggers the coincident incorporation of modular release sites to consolidate stable synapse function.

## Introduction

Neurotransmitter-laden synaptic vesicles (SVs) release their content at presynaptic active zones (AZs) in response to Ca^2+^ influx through voltage gated channels that respond to action-potential (AP) depolarization. Neurotransmitter binding to postsynaptic receptors subsequently leads to their activation for synaptic transmission. Modulation of transmission strength is called synaptic plasticity. Long-term forms of synaptic plasticity are major cellular substrates for learning, memory, and behavioral adaptation^1, 2^. Mechanisms of long-term synaptic plasticity modify the structure and function of the presynaptic terminal and/or the postsynaptic apparatus. AZs are covered by complex scaffolds composed of a conserved set of extended structural proteins. ELKS/Bruchpilot (BRP), RIM, and RIM-binding protein (RBP) functionally organize the coupling between Ca^2+^-channels and release machinery by immobilizing the critical (M)Unc13 release factors in clusters close to presynaptic Ca^2+^-channels and thus generate SV release sites, at both mammalian and *Drosophila* synapses^3-12^. Whether and how discrete AZ release sites and the associated release machinery are reorganized during plastic changes remains unknown.

One crucial form of presynaptic plasticity is the homeostatic control of neurotransmitter release. This process, referred to as presynaptic homeostatic potentiation (PHP), is observed in organisms ranging from invertebrates to humans, but is perhaps best illustrated at the larval neuromuscular junction (NMJ) of *Drosophila melanogaster*^13, 14^. Here, PHP requires the core AZ-scaffolding proteins RIM, RBP and Fife^15-17^ and physiologically coincides with the upregulation of SV release sites^17, 18^. Yet it is unknown how these AZ-scaffolds mediate release site addition, which downstream molecules are needed for PHP, whether AZ-scaffold independent reactions occur and whether these mechanisms extend to other forms of plasticity, e.g. during learning in the central nervous system.

Here, we combine genetic and electrophysiological analysis to reveal a molecular sequence that triggers structural remodeling of AZ scaffolding proteins (BRP/RBP) that ultimately lead to (M)Unc13 addition within minutes. Using super-resolution light microscopy we identify a modular AZ nano-architecture built by these proteins (which corresponds to SV release sites) that rapidly extends by incorporating additional modules for plasticity. This “rapid remodeling” critically depends on the core AZ scaffolding proteins RBP/BRP, but neither on the early AZ assembly factors Liprin-α/Syd-1, nor on RIM or Fife. Additionally, AZ-remodeling was abolished in transport mutants previously shown to promote BRP/RBP transport. Strikingly, rapid addition of AZ nano-clusters was not required for the immediate expression of PHP on a minutes’ timescale, but was essential to sustain potentiation thereafter. We identify Unc13A as a direct molecular target for PHP in experiments in which Unc13A was delocalized from the AZ scaffolds. This mutant displayed sizable synaptic transmission but fully lacked PHP and AZ-remodeling. The same interference in mushroom body Kenyon cells of the *Drosophila* brain eliminated short-term memory, indicating that Unc13A is also a plasticity target in the central nervous system. In summary, we show that synapses capitalize on the available AZ material for immediate potentiation, but coincidently undergo release site addition via modular building blocks to consolidate stable synaptic potentiation. Thus, our work lays a foundation from which to mechanistically understand a likely conserved presynaptic plasticity process important for dynamically adjusting and stabilizing neurotransmission across multiple timescales.

## Results

### Rapid and chronic homeostatic plasticity regulate AZ protein levels

As a robust paradigm for assessing presynaptic plasticity over different time scales, we focused on PHP, which is well characterized at *Drosophila* NMJs^13^. To induce plasticity on a timescale of minutes, postsynaptic ionotropic glutamate receptors were partially blocked using the non-competitive open-channel blocker Philanthotoxin-433 (PhTx)^19^ (Fig. 1a-d). This reduces postsynaptic sensitivity to neurotransmitter release from single SVs (reflected in a reduction of the amplitude of spontaneously occurring “minis”, single SV fusion events, Fig. 1b). Initially, this also leads to a proportional decrease in AP-evoked transmission, but in less than 10 minutes, PHP increases the number of SVs released per AP (quantal content) to compensate for the postsynaptic interference, resulting in AP-evoked transmission comparable to baseline levels (Fig. 1b)^19^. To identify molecular adaptations during plasticity, we investigated whether the levels of any of the evolutionarily conserved AZ proteins were altered. We thus immunostained against BRP, RBP, Unc13A (we focused on Unc13A, the Unc13 isoform dominating evoked SV release at *Drosophila* NMJ synapses^7^; flybase: unc-13-RA), Syx-1A, Unc18 and Syd-1 (as motoneuronally expressed Syd-1-GFP) (Fig. 1c; Supplementary Fig. 1a). In agreement with previous observations^18, 20^, we found that 10 minutes of PhTx treatment increased AZ BRP-levels by about 50% (Fig. 1c,d). In addition, we found that RBP, Unc13A and Syx-1A increased by about 30%, 60%, and 65%, respectively (Fig. 1c,d). The AZ levels of RBP/BRP, Unc13A/BRP and Syx-1A/BRP scaled proportionally over all AZ sizes (Supplementary Fig. 1b; Ctrl, black lines). This proportionality was preserved upon PhTx treatment (Supplementary Fig. 1b; PhTx, blue lines). Notably, the AZ-levels and distribution of the essential Sec1/(M)Unc18 family protein Unc18 – which was recently found to function in PHP^21^ – were unaffected (Fig. 1c,d;), demonstrating specific up-regulation of a subset of AZ proteins. Another AZ protein, the assembly factor Syd-1 even displayed a slight reduction upon PhTx (Supplementary Fig. 1a), further underscoring a high degree of specificity.

**Figure 1:**
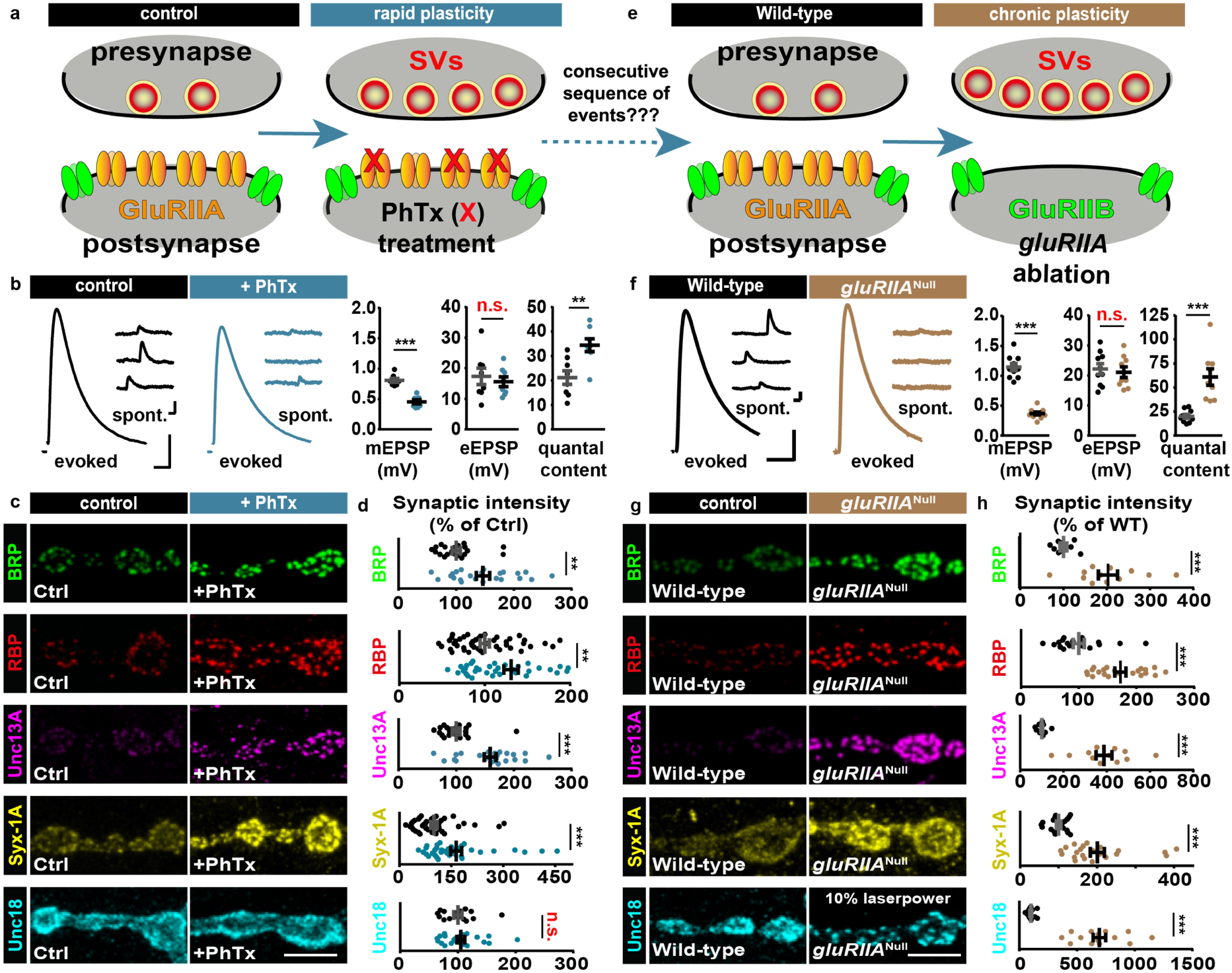
Rapid homeostatic plasticity regulates AZ protein levels. **(a)** Sketch of investigated conditions for rapid plasticity: control synapses (left) are compared with rapid plasticity (10 minutes of PhTx (red “X”); right). Rapid plasticity increases the number of SVs released (red). **(b,f)** Representative traces of eEPSP (evoked), mEPSP (spont.) and their quantification in Ctrl (black) and PhTx (blue) treated (b) or wild-type (black) and *gluRIIA*^*Null*^ (brown) (f) cells. **(c,g)** NMJs labelled with indicated antibodies in Ctrl (black) and PhTx (blue) treated (c) or in wild-type (black) and *gluRIIA*^*Null*^ (brown) (g) animals. **(d, h)** Quantification of BRP, RBP, Unc13A, Syx-1A and Unc18 AZ-levels in % of Ctrl in Ctrl (black) and PhTx (blue) treated (d) or in % of Wild-type (WT) in Wild-type (black) and *gluRIIA*^*Null*^ (brown) (h) animals. **(e)** Sketch of investigated conditions for chronic plasticity: Wild-type synapses (left) are compared with *gluRIIA*^*Null*^ mutants. Chronic plasticity greatly increases the number of SVs released. See also Supplementary Fig. 1. Exact normalized and raw values, detailed statistics including sample sizes and P values are listed in Supplementary Table 1. Scale bars: (b,f) eEPSP: 25 ms, 5 mV; mEPSP: 50 ms, 1 mV; (c,g) 5 µm. Statistics: Student’s unpaired T-test was used for comparisons in (b) and (f) mEPSP amplitude, quantal content and Mann-Whitney U test for all other comparisons. **P ≤ 0.01; ***P ≤ 0.001; n.s., not significant, P > 0.05. All panels show mean ± s.e.m.

To verify that these AZ-adaptations were specific to functional glutamate receptor interference, and to address their relevance over a longer time window, we investigated larvae bearing mutations in a glutamate receptor subunit (Fig. 1e-h). Deletion of the high-conductance receptor subunit IIA (GluRIIA) results in a similarly reduced postsynaptic sensitivity to single SV fusion events (Fig. 1f) as PhTx treatment (Fig. 1b). Because under these circumstances PHP also increases quantal content to achieve similar AP-evoked transmission (Fig. 1f), *gluRIIA* mutants have extensively been used to investigate long-term PHP (over the 3-4 days of larval development)^22, 23^. Immunostainings against BRP, RBP, Unc13A, and Syx-1A confirmed their (in this case larger) elevation on this longer timescale (compare Fig. 1g,h with 1c,d) (100%, 70%, 400% and 200%, respectively, compared to 50%, 30%, 60% and 65% upon PhTx treatment). Unlike the stoichiometric increase observed for BRP/Unc13A within minutes, this long-term PHP revealed enhanced Unc13A AZ-incorporation (Supplementary Fig. 1c). Another distinction was a remarkable reorganization and 8-fold increase of Unc18 (Fig. 1g,h; Supplementary Fig. 1d,e). Our data thus imply that considerable AZ restructuring occurs within minutes of PHP induction, which is further enhanced across longer-lasting timescales.

### Rapid addition of Unc13A/BRP/RBP release site modules during homeostatic plasticity

We next sought to investigate how the altered levels of AZ proteins during PHP affected their nanoscopic topology by super-resolution STED microscopy (x-y resolution ∼ 30-40 nm). As noted before, planar AZs revealed clearly distinguishable individual Unc13A/BRP/RBP spots arranged in a ring-like geometry^3, 7-9^ (Fig. 2a). It was recently shown by single molecule imaging, these individual clusters likely contain several (probably few tens of) molecules in the case of BRP ^24^. We detected the number of clusters per AZ in single AZ-images with a simple peak detection algorithm which was largely heterogeneous for all three proteins (Fig. 2c-e (black bars)). However, in all cases the cluster number per AZ increased upon PhTx-treatment (Fig. 2c-e (blue bars)), and slightly increased further in *gluRIIA*^Null^ mutants (Supplementary Fig. 2a; brown bars). With increasing AZ-cluster number the AZ diameter (measured from the AZ center to the center of the clusters) also increased in both conditions (Supplementary Fig. 2b,c), consistent with previous STED analysis performed on BRP-rings ^18^. Notably, the observed remodeling only affected cluster numbers, and did not alter their intensities (Supplementary Fig. 2b,c). Thus, the first conclusion of our analysis is that PHP increases the number Unc13A/BRP/RBP nano-clusters per AZ within minutes.

**Figure 2:**
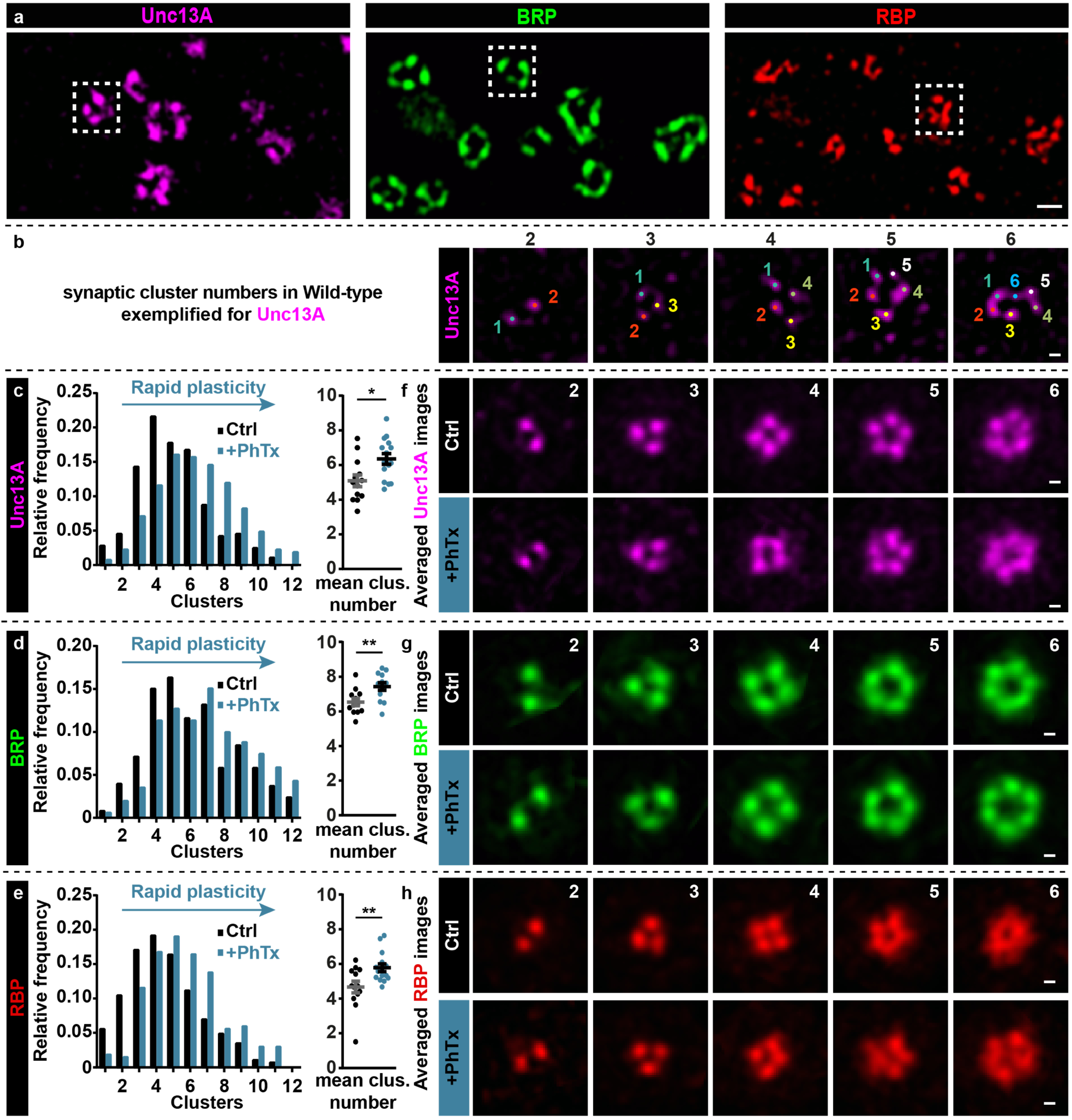
Rapid plasticity alters Unc13A/BRP/RBP AZ protein cluster number. **(a)** STED microscopy images containing several AZs with variable numbers of protein clusters of Unc13A (magenta); BRP (green) and RBP (red). Dashed white boxes mark one AZ example used for the cluster number counting. **(b)** Example AZs with 2-6 Unc13A clusters, marked by colored dots and used for cluster number counting. **(c-e,** left) Frequency distribution of Unc13A (c), BRP (d) and RBP (e) modules per AZ either without (Ctrl, -PhTx; black) or with PhTx (+PhTx; blue) treatment. **(f-h)** Average of rotated STED images stained against Unc13A (f), BRP (g) and RBP (h) with 2-6 modules either without (Ctrl) or with PhTx (+PhTx) treatment. See also Supplementary Figs. 2-4. Exact values, detailed statistics including sample sizes and P values are listed in Supplementary Table 1. Scale bars: (a) 200 nm; (b,f-h) 50 nm. Statistics: Mann-Whitney U test. n.s., not significant, P > 0.05. Panels (c-e) show NMJ-wise means of AZ-mean ± s.e.m.

We also wanted to investigate whether the overall AZ-topology changed upon cluster incorporation. Notably, averaging of STED-images was recently used to generate a three-dimensional model of an “average” synapse, displaying the mean protein localization at high resolution ^25^. Thus, to compare the overall single AZ-topology, AZ images were centered, sorted by the number of clusters, aligned by rotation, and averaged. Two different alignment methods were used. In the first procedure, images were simply rotated such that the cluster with the highest intensity was positioned to the top (Supplementary Fig. 3a and Methods for details). Even though this procedure only targeted a single pixel per image (the position of the brightest cluster), the remaining (lower intensity) clusters were often found in similar relative positions, such that averaging revealed a simple polygonal geometrical series (Supplementary Fig. 3a), demonstrating some regularity. In a more refined analysis, we simultaneously considered the position of all clusters per AZ (Supplementary Fig. 3c and Methods for details) which also revealed a simple geometrical pattern for Unc13A, BRP and RBP (Fig. 2f-h and Supplementary Fig. 3c). This stereotypical arrangement was best seen for AZs containing 2-6 clusters but less clear for AZs containing more than that (which could mean that these are less regular; Supplementary Fig. 2d,e). This arrangement was unaltered upon PhTx-treatment or *gluRIIA* ablation (Figure 2f-h and Supplementary Fig. 4a-c). Notably, the results of these averaging approaches obviously produced specific patterns, as neither random categorization of the single AZ images nor applying this methodology to Syx-1A and Unc18 (which are diffusely distributed at the AZ) resulted in regular but instead in highly irregular/random fluorescence patterns (Supplementary Figs. 3b and 2f,g). This also demonstrates that structural features are only conserved across AZs containing the same number of clusters. It should be noted that only the average images (Fig. 2f-h; Supplementary Fig. 4) depend on this procedure, while detecting the effects of PhTx or *gluRIIA* ablation on cluster numbers (Fig. 2c-h) was fully independent of this.

Thus, these findings imply that complexes of the core AZ-scaffold form discrete nano-modular structures, which correspond to SV release sites, and that rapid presynaptic plasticity triggers their fast AZ-incorporation which is even further enhanced over longer-timescales.

### Mutations that impair BRP/RBP transport disrupt rapid AZ-remodeling

The remarkable remodeling of AZ material within the short timeframe of PhTx-treatment (minutes) raised the question of how this is mechanistically achieved. We first considered whether local presynaptic protein translation could be required ^26^. However, treatment of larvae with 50 µg/ml of the translation blocker cycloheximide (prior and during PhTx-treatment) did not disrupt structural remodeling of BRP or vGlut (vesicular glutamate transporter) at AZs (Supplementary Fig. 5a), consistent with re-modeling being translation-independent. Moreover, the functional increase in quantal content remained expressed in the presence of the blocker (Supplementary Fig. 5b), consistent with previous reports^19, 20^.

Because active kinesin-dependent protein transport is required for long-term homeostatic plasticity in *gluRIIA*^Null^ mutants^27^, we asked whether BRP/RBP transport mechanisms might be employed for AZ remodeling. For this we investigated proteins involved in BRP/RBP transport by their mutation which causes abnormal BRP/RBP accumulation in the moto-neuronal axons, such as Atg1 (Unc-51)^28^, serine–arginine (SR) protein kinase at location 79D (Srpk79D;^29, 30^), and App-like interacting protein (Aplip-1, Jip1 or JNK interacting protein in mammals), a selective RBP transport-adaptor^31^ (Fig. 3a,b). While we observed clear PhTx-induced BRP-/Unc13A-upscaling in Wild-type controls as well as in *atg1* mutants (Fig. 3a,b), remodeling was fully absent upon null-mutation of *srpk79D, aplip-1*^32^ or in animals bearing an Aplip-1 point mutation that selectively prevents kinesin light chain interaction (*aplip-1*^*ek4*^)^33^ (Fig. 3a,b). Additionally, STED microscopy revealed that *aplip-1*^ek4^ and *srpk79D*^ATC^ mutants appeared to contain fewer BRP/Unc13A clusters per AZ on average than Wild-type (compare Fig. 2c,d with Fig. 3c) and fully lost the capacity to increase cluster numbers upon PhTx-treatment (even a decrease was observed, Fig. 3c). We also discovered that upon motoneuronal Aplip-1 or Srpk79D knock-down, Unc13A-GFP co-accumulated with aberrant axonal BRP aggregates (Supplementary Fig. 5c,d). Interestingly, a partial co-accumulation of BRP/Unc13A-GFP (but to a lower extent in comparison to the knock-downs) was also present in the control situation, in line with an at least partial co-transport that we occasionally observed in live-imaging experiments (Supplementary Fig. 5c-g; Movie 1).

**Figure 3:**
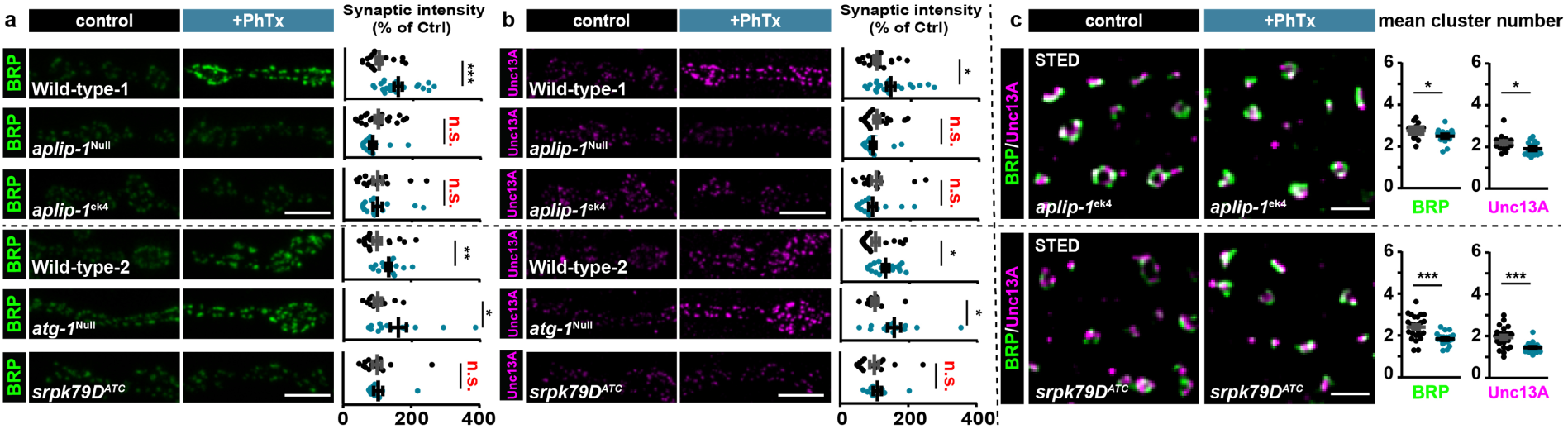
Rapid AZ-remodeling during homeostatic plasticity requires Aplip-1 and Srpk79D. **(a,b)** Muscle 4 NMJs of segment A2-A5 from 3rd instar larvae and quantification of synaptic levels of Wild-type – 1, *aplip-1*^Null^, *aplip-1*^*ek*4^, Wild-type – 2, *atg1* and *srpk79D*^*ATC*^ labelled with the indicated antibodies without (Ctrl; black) and with 10 minutes of PhTx (+PhTx; blue) treatment. Two independent experiments were performed, Wild-type – 1 was used as control for *aplip-1*^Null^ and *aplip-1ek*^4^ while Wild-type – 2 was used for *atg1*^Null^ and *srpk79D*^*ATC*^. **(c)** Average BRP and Unc13A cluster number per AZ either without (Ctrl, -PhTx; black) or with PhTx (+PhTx; blue) treatment in *aplip-1*^*ek*4^ and *srpk79D*^ATC^. See also Supplementary Figs. 5, 6 and Movie 1. Exact normalized and raw values, detailed statistics including sample sizes and P values are listed in Supplementary Table 1. Scale bars: (a,b) 5 µm; (c) 500nm. Statistics: Mann-Whitney U test. * P ≤ 0.05; **P ≤ 0.01; ***P ≤ 0.001; n.s., not significant, P > 0.05. All panels show mean ± s.e.m.

We further wanted to elaborate on the involvement of active protein transport during PhTx-induced AZ-remodeling by interfering with the cytoskeletal “tracks” used for short-range transport with Latrunculin B (an actin polymerization blocker). While the AZ levels of BRP/Unc13A were already slightly enhanced by Latrunculin B treatment (Supplementary Fig. 6a,b), PhTx-treatment failed to induce the typical increase of the AZ levels of these proteins (in fact a reduction was observed, Supplementary Fig. 6a,b). Thus, the actin cytoskeleton and active BRP/RBP transport are required for rapid AZ remodeling.

### Rapid homeostatic remodeling of AZ structure depends on BRP and RBP

We next investigated which of the evolutionarily conserved core AZ scaffolding proteins are required for rapid AZ-remodeling. Loss of RBP fully blocked the rapid, PhTx-induced increase of BRP/Unc13A (Fig. 4a,b). Also BRP was essential, because the typical increase in Syx-1A and Unc13A observed upon PhTx treatment was abolished (Compare Fig. 4a,b with Fig. 1c,d). Notably, BRP-amounts appear to be rate-limiting because PhTx-induced AZ-remodeling was blocked in larvae heterozygous for a *brp* null allele (brp^Null^/+)(Fig. 4a,b). In contrast, null mutation of RIM, which abolished PHP^17^, did not interfere with AZ-remodeling (Fig. 4a,b). Furthermore, the simultaneous deletion of RIM and Fife (a possible RIM homologue which is required for PHP^15, 34^) did not interfere with AZ-remodeling. Thus, RIM and Fife appear to act downstream of BRP/RBP and are non-essential for AZ-remodeling (Fig. 4a,b).

**Figure 4:**
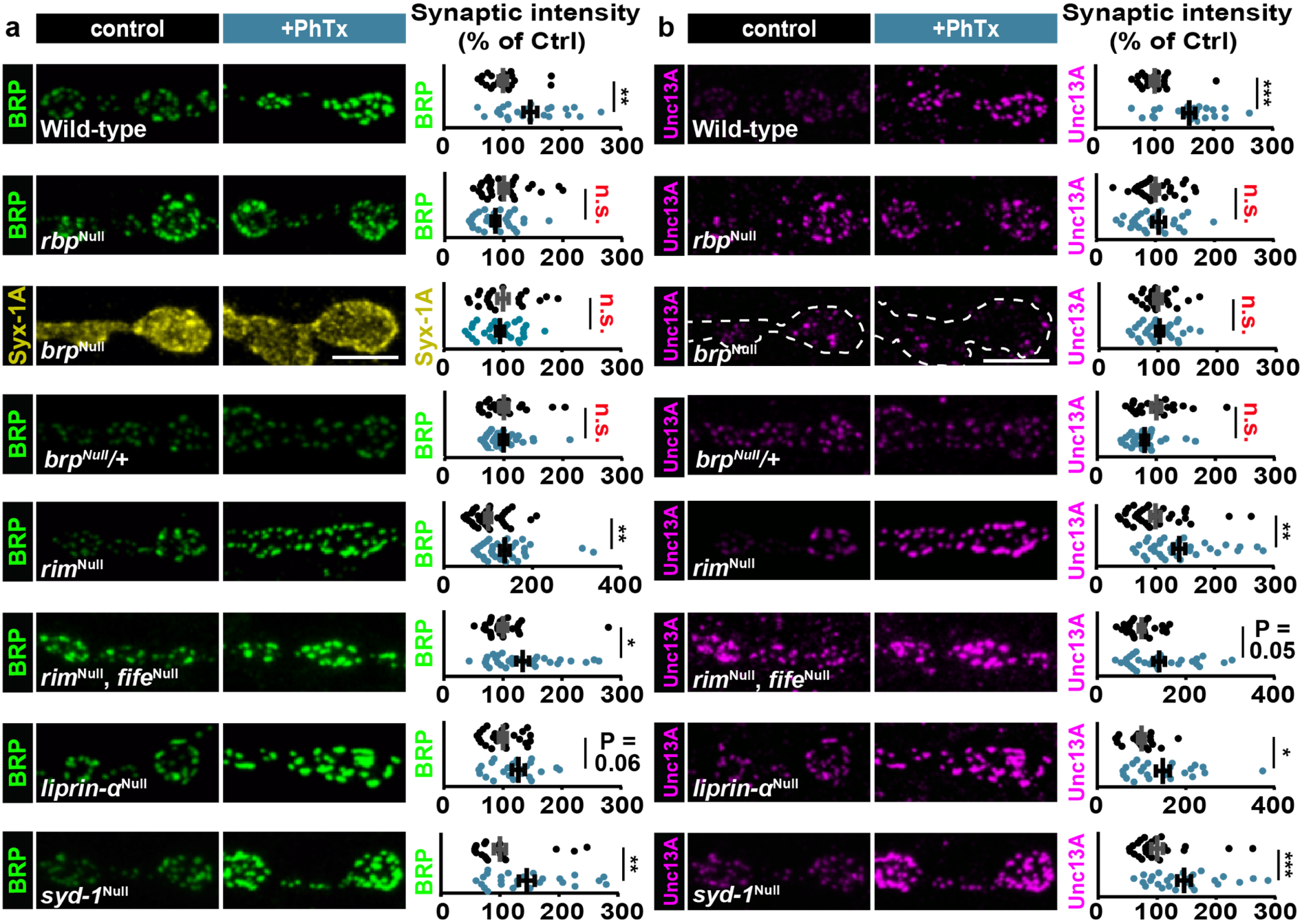
Rapid AZ-remodeling during homeostatic plasticity requires BRP and RBP. **(a,b)** Confocal images and quantification of synaptic intensities of muscle 4 NMJs of abdominal segment 2-4 from 3rd instar larvae at Wild-type,, *rbp*^Null^, *brp*^Null^, *brp*^Null^/+, *rim*^Null^, *rim*^Null^,*fife*^Null^, *liprin-α*^Null^ and *syd1*^Null^ NMJs labelled with the indicated antibodies without (control; black) and with 10 minutes PhTx (+PhTx; blue) treatment. For Wild-type, images and data were adapted and replotted from Fig. 1. Please note, although Wild-type controls were performed in parallel to every mutant genotype to check for functional AZ-remodeling upon PhTx-treatment in each set of experiments, we do not show all WT-control here due to space limitations. Therefore, AZ-protein levels should not be compared between genotypes. Exact normalized and raw values (also of additional Wild-type controls for genotypes where PhTx-treatment failed to induce AZ-remodeling), detailed statistics including sample sizes and P values are listed in Supplementary Table 1. Scale bars: 5 µm. Statistics: Mann-Whitney U test. * P ≤ 0.05; **P ≤ 0.01; ***P ≤ 0.001; n.s., not significant, P > 0.05. All panels show mean ± s.e.m.

AZ assembly is initiated by the conserved scaffolding proteins Liprin-α and Syd-1, which both regulate AZ size^35-41^. We reasoned that AZ growth –as observed here during plasticity– may capitalize on the same molecular machinery as *de novo* AZ formation, and therefore tested whether BRP/Unc13A-scaling depended on those proteins. However, *liprin-α*^Null^ and *syd-1*^Null^ mutants revealed normal PhTx-induced BRP/Unc13A-scaling (Fig. 4a,b), indicating that these factors are dispensable. Thus, systematic investigation of evolutionarily conserved AZ scaffolding proteins reveals a selective dependence on the core AZ-scaffolds BRP and RBP for structural remodeling during plasticity.

### AZ-remodeling is required for the chronic, but not rapid, functional expression of homeostatic plasticity

Several studies have shown that presynaptic release positively correlates with AZ-size^3, 42-45^. Therefore, we expected that the increase of AZ-BRP/Unc13A observed upon PhTx treatment would functionally increase presynaptic release (Fig. 1a-d). However, it is not entirely obvious whether the AZ-remodeling (which continues beyond the minutes’ timescale during long-term PHP (Fig. 1e,g,h) would be essential for rapid PHP. For instance, loss of RBP was shown to occlude both AZ remodeling (Fig. 4) and the functional increase in quantal content ^16^, suggesting a pivotal role in both adaptations. Yet PHP and AZ-remodeling do not go hand-in-hand in the case of RIM (and Fife) mutants, whose AZs remodel, but which cannot express PHP (increased quantal content^15, 17^). These observations prompted us to systematically investigate the relevance of AZ-remodeling for the rapid induction and sustained expression of PHP.

We first investigated the dependence of rapid PHP on the rapid remodeling of Unc13A/Syx-1A in *brp*^Null^ larvae. Strikingly, AZ-remodeling was blocked (Fig. 4a,b), but functional PHP expression (an increased quantal content) persisted at levels comparable to the Wild-type/Control situation (Fig. 5a,b). In addition, mutants of Srpk79D (whose AZs also do not undergo PhTx-induced AZ-remodeling; Fig. 3a,b and replotted in Fig. 5d) were likewise able to increase their quantal content (Fig. 5c), showing that AZ remodeling can be uncoupled from rapid PHP expression. Thus, even though AZ-remodeling occurs on a similar time-scale, it is not required to rapidly enhance the quantal content in these cases.

**Figure 5:**
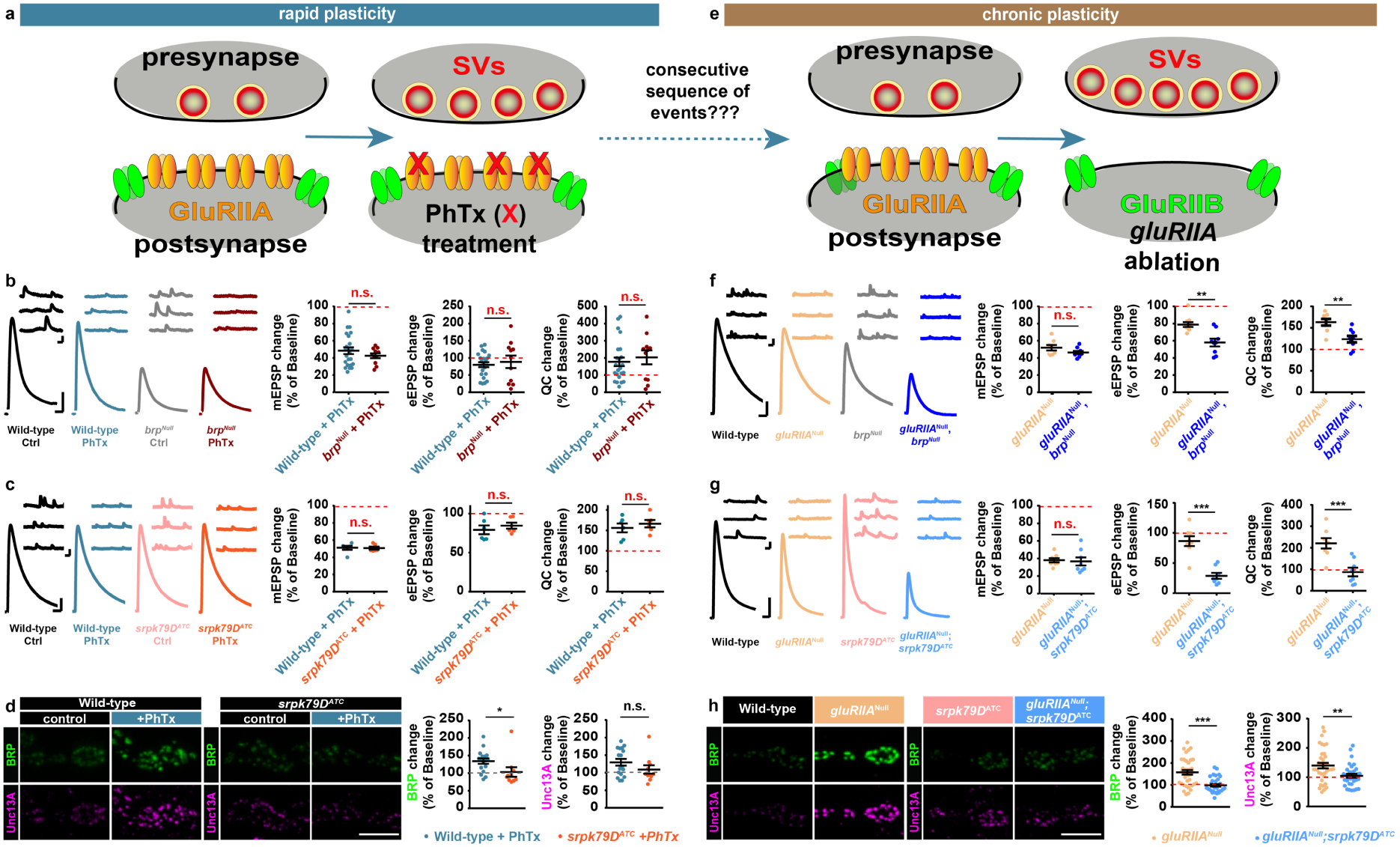
Structural AZ-remodeling sustains NT-release potentiation over longer timescales. **(a)** Sketch of investigated conditions for rapid plasticity: control synapses (left) are compared with rapid plasticity (10 minutes of PhTx (red “X”); right). Rapid plasticity increases the number of SVs released (red). **(b,c)** (left) Representative traces of eEPSP (evoked) and mEPSP (spont.) of the indicated genotypes with and without PhTx-treatment. (Right) Quantifications of percentage change of mEPSP amplitude, eEPSP amplitude and quantal content (QC) upon PhTx-treatment. Values are divided by the corresponding measurement in the absence of PhTx for each genotype (dashed red line corresponds to 100%/no change). **(d)** (left) Confocal images of muscle 4 NMJs of abdominal segment 2-5 from 3rd instar larvae at Wild-type and *srpk79D*^ATC^ NMJs labelled with the indicated antibodies without (control; black) and with 10 minutes PhTx (+PhTx; blue) treatment. (Right) Quantification of percentage change of synaptic BRP and Unc13A levels in Wild-type (blue) and *srpk79D*^ATC^ (orange) upon PhTx-treatment compared to baseline of control treatment for each genotype (dashed red line). Data are modified from Fig. 2. **(e)** Sketch of investigated conditions for chronic plasticity: Wild-type synapses (left) are compared with *gluRIIA*^*Null*^ mutants. Chronic plasticity greatly increases the number of SVs released. **(f,g)** Same as in (b,c) but compared to baseline of each control genotype. **(h)** Same as in (d) but compared to baseline fluorescence values of Wild-type for *gluRIIA*^Null^ and *srpk79D*^ATC^ for *gluRIIA*^Null^;*srpk79D*^ATC^. See also Supplementary Fig. 7. Exact normalized and raw values, detailed statistics including sample sizes and P values are listed in Supplementary Table 1. See also Supplementary Figure 10 and 11 for non-normalized values. Scale bars: eEPSP: 25 ms, 5 mV; mEPSP: 50 ms, 1 mV; (d,h) 5 µm. Statistics: Student’s unpaired T-test was used for comparisons in (b) mEPSP change, (f), (g) and Mann-Whitney U test for all other comparisons. *P ≤ 0.05; **P ≤ 0.01; ***P ≤ 0.001; n.s., not significant, P > 0.05. All panels show mean ± s.e.m.

We next investigated whether elevation of the AZ protein levels was required to consolidate the increased quantal content over chronic time-scales in *gluRIIA*^Null^ mutants (Fig. 5e-h). Indeed, PHP was severely impaired in *brp*^Null^,*gluRIIA*^Null^ double mutants (Fig. 5f). In an otherwise Wild-type background, the increase in quantal content (upon *gluRIIA*-null mutation) was much larger than when BRP was additionally deleted (Fig. 5f). We could ensure that the impairment was not due to the overall reduced release in *brp*^*Null*^ mutants, as a loss to compensate for the *gluRIIA* ablation was also seen in *srpk79D*^*ATC*^ mutants (which had comparable synaptic transmission to Wild-type cells (Fig. 5g), no AZ-remodeling upon PhTx-treatment (Fig 3a,b), intact PHP upon PhTx-treatment (Fig. 5c) and severely impaired PHP expression in *gluRIIA*^Null^ (Fig. 5g)). Importantly, AZ-remodeling was also fully blocked in *gluRIIA*^Null^;*srpk79D*^ATC^ double mutants (Fig. 5h). An intermediate behavior was seen in the case of the *aplip-1*^*ek*4^ mutant, (Supplementary Fig. 7), possibly because other transport adapters might compensate in this situation. Together, this suggests that PHP rapidly increases neurotransmitter release through modulation of the available AZ components, but in addition immediately induces AZ-remodeling to ensure its consolidation.

### Presynaptic potentiation requires Unc13A

We next sought to identify the molecular substrate of PHP. Previous experiments established a requirement of the ?1 voltage gated Ca^2+^-channel subunit Cacophony (Cac)^19, 46^. In line with this, the levels of Cac as well as the Ca^2+^-influx increase upon PhTx treatment (Fig. 6a,b)^46-48^. We furthermore investigated Unc13A. A slight PhTx-induced BRP-/RBP-scaling persisted upon Unc13A loss, but was weaker than in the Wild-type situation (Fig. 6a,b), possibly due to slightly elevated BRP/RBP-AZ-levels already in the non-PhTx-treated *unc13A*^Null^ situation^7^. Notably, Cac-levels were still increased, even to a slightly larger extent than in the Wild-type situation (Fig. 6a,b). However, functional PHP, the increase in quantal content, was completely lost (Fig. 6c,d). This indicates that Unc13A –like RIM and RBP^16, 17^– plays an essential role in the plastic enhancement of NT release during PHP.

**Figure 6:**
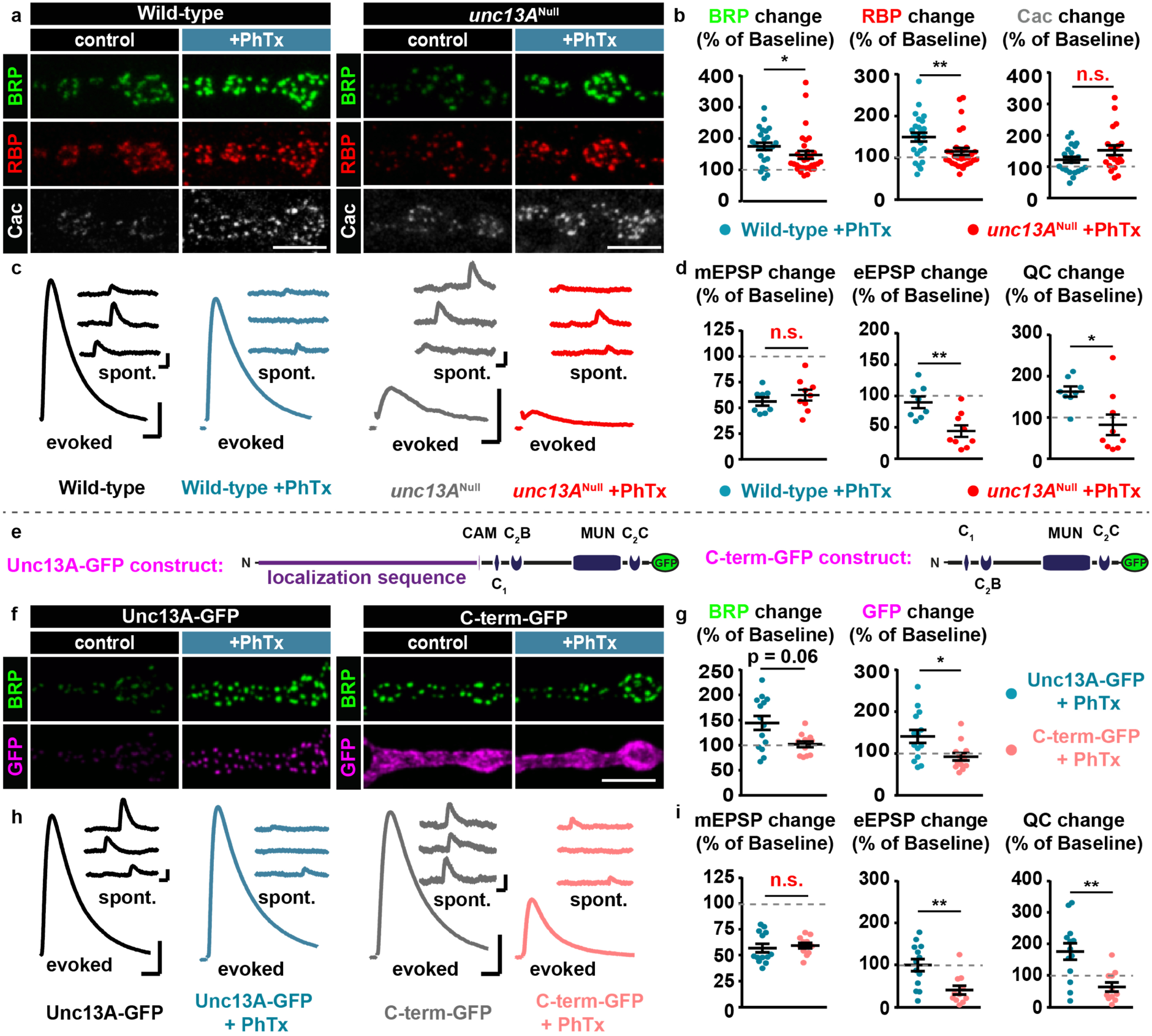
Unc13A and its N-terminus are critical for rapid PHP and AZ-remodeling. **(a)** Confocal images of muscle 4 NMJs of abdominal segment 2-5 from 3rd instar larvae at Wild-type (left) and *unc13A*^Null^ (right) NMJs labelled with the indicated antibodies without (control; black) and with 10 minutes PhTx (+PhTx; blue) treatment. **(b)** Quantification of percentage change of synaptic BRP, RBP and Cac AZ-levels in Wild-type (blue) and *unc13A*^Null^ (red) upon PhTx-treatment compared to the same measurement in the absence of PhTx for each genotype (dashed grey line indicates 100%/no change). **(c)** Representative traces of eEPSP (evoked) and mEPSP (spont.) in Wild-type and *unc13A*^Null^ animals without (Ctrl; black or grey) and with 10 minutes PhTx (+PhTx; blue or light red) treatment. **(d)** Quantifications of percentage change of mEPSP amplitude, eEPSP amplitude and quantal content (QC) in PhTx-treated Wild-type (blue) and *unc13A*^Null^ (light red) cells compared to the same measurement obtained without PhTx for each genotype. Traces for (c) were replotted from Fig. 1. **(e)** Left: Full length Unc13A construct used in rescue experiments of *unc13*^Null^ animals. Functional domains for AZ localization, Calmodulin-(CAM), lipid-binding (C1, C2B, C2C) and the MUN domain relevant for SV release are shown. Right: Schematic of Unc13A construct lacking the N-terminal localization sequence (C-term-GFP rescue). **(f-i)** Same as in (a-d) for cells re-expressing Unc13A-GFP (blue) or C-term-GFP (light red) in the *unc13*^Null^ background. See also Supplementary Figure 8. See also Supplementary Figure 10 and 11 for non-normalized values. Exact normalized and raw values, detailed statistics including sample sizes and P values are listed in Supplementary Table 1. Scale bars: (a,f) 5 µm; (c,h) eEPSP: 25 ms, 5 mV; mEPSP: 50 ms, 1 mV. Statistics: Student’s unpaired T-test was used for comparisons in (d), (i) mEPSP change and Mann-Whitney U test for all other comparisons. *P ≤ 0.05; **P ≤ 0.01; ***P ≤ 0.001; n.s., not significant, P > 0.05. All panels show mean ± s.e.m..

### The Unc13A N-terminus is critical for rapid PHP, AZ-remodeling, and learning

The observation that Unc13A is essential for PHP is fully consistent with the previous findings that RIM and RBP are required (^16, 17^, because these proteins likely function in Unc13A AZ recruitment and activation (see discussion). As in other species (*M*. *musculus*/*C*. *elegans*), this interaction depends on the (M)Unc13 N-terminus^7, 49-53^. To investigate the functional relevance of the Unc13A N-terminus for rapid PHP and AZ-remodeling, we used an Unc13A mutant lacking the N-terminal AZ-localization sequence (named C-term-GFP; Fig. 6e), which uncouples Unc13A from the central BRP/RBP scaffold^3^ (and therefore supposedly also uncouples SV fusion from a possible regulatory function of RIM –see discussion). Importantly, the magnitude of AP-evoked synaptic transmission in these mutants was largely restored compared to the detrimental effect of *unc13*^Null^ mutation (compare Fig. 6h (C-term-GFP; grey traces) with Fig. 6h (full-length Unc13A-GFP; black traces) and ^3^). However, in contrast to control larvae (*Unc13*^Null^ with full-length Unc13A-GFP rescue; Fig. 6f-i), C-term-GFP mutants (*Unc13*^Null^ with C-term-GFP rescue; Fig. 6f-i) completely lacked AZ-remodeling (Fig. 6f,g) (note that BRP levels were already enhanced in the non-PhTx-treated group, Supplementary Fig. 8a,b), Furthermore, unlike in control larvae, no rescue of evoked transmission and no increase in quantal content was seen upon PhTx treatment, indicating that C-term mutants were deficient of functional PHP (Fig. 6h-i). This demonstrates a dependence of PHP on the Unc13A N-terminus.

Although synapses vary tremendously in their excitability, input/output relationship, and transmitter type, the presynaptic release machinery is remarkably conserved in most systems and across species^54^. Thus, we wondered whether the principles of rapid presynaptic adaptation of the peripheral nervous system might be utilized in other synapse types and in other forms of plasticity, like the ones involved in learning and memory formation. Because short-term memory functions on timescales comparable to the PhTx-induced rapid homeostatic plasticity^55^, we investigated whether the Unc13 C-term mutant also exhibited learning deficits.

We specifically expressed C-term-GFP alone, or while simultaneously knocking down endogenous full-length Unc13A (Unc13A-RNAi), in mushroom body Kenyon cells (KCs). KC output synapses undergo learning-induced plasticity and are required for the formation of short-term memories^56^. Knockdown of endogenous Unc13A and expression of C-term-GFP were confirmed using antibodies against GFP (labelling C-term-GFP but not endogenous protein) and the Unc13A N-terminus (labelling the endogenous protein but not C-term-GFP where the N-terminal epitope is deleted) (Fig. 7a). Additionally, we confirmed the strong efficacy of the Unc13A-RNAi via Western blot (knock-down in the entire brain using the pan-neuronal elav-Gal4 driver; Supplementary Fig. 9a).

**Figure 7:**
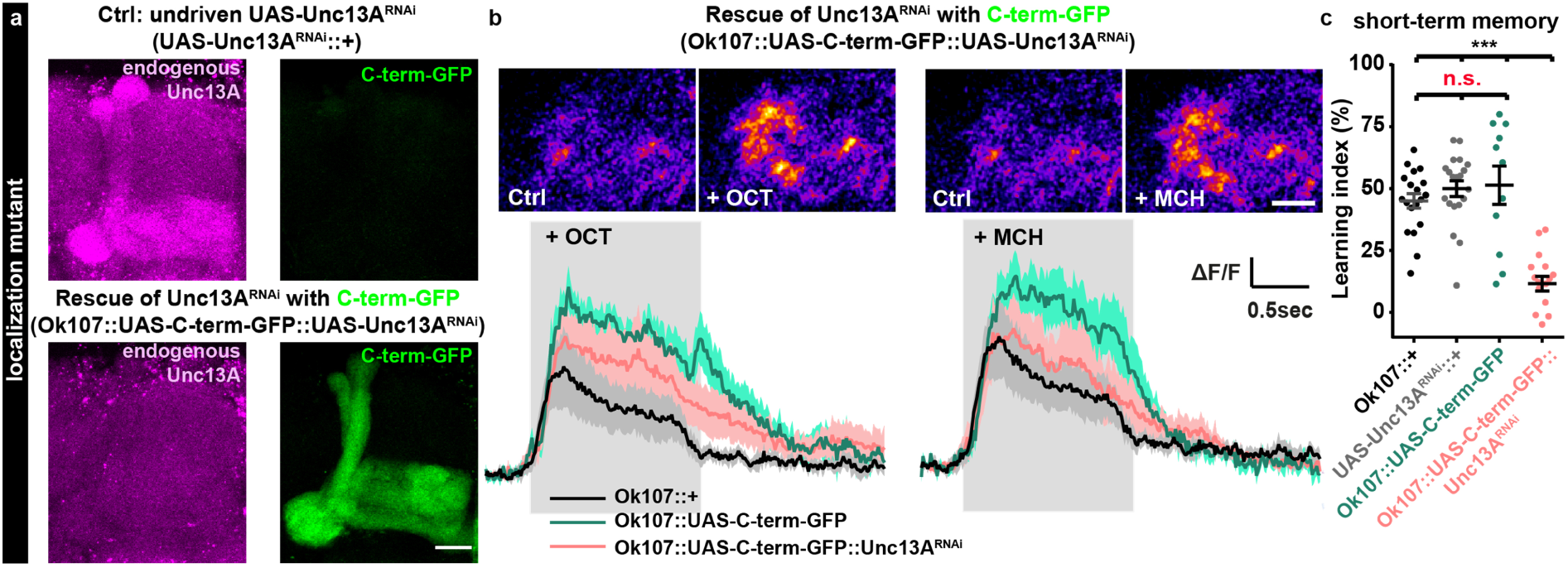
Mutants incapable of PHP at the NMJ impair short-term memory when expressed in the olfactory learning center of the fly brain. **(a)** Confocal images of adult *Drosophila* mushroom-body regions of control (top; undriven UAS-Unc13A^RNAi^ (UAS-Unc13A^RNAi^::+)) and rescue of driven UAS-Unc13A^RNAi^ with C-term-GFP (bottom; Ok107::UAS-C-term-GFP::UAS-Unc13A^RNAi^) brains labelled with the indicated antibodies. **(b)** Averaged odor responses measured at the level of presynaptic boutons of M4/6 (MBON-β’2mp/MBON-γ5β’2a/MBON-β2β’2a) mushroom body output neurons (compare^57^ or^96^). Above: sample images of two-photon recordings from M4/6 cells (20 frames averaged respectively) before and after odor onset. Grey shading indicates the time at which the odor was applied. Left panels OCT and right MCH response. Below: averaged odor responses. 5 responses per odor were averaged per animal. Solid lines show mean responses (n = 5 to 6 animals per genotype). Shaded areas represent the SEM. **(c)** Short-term memory scores after mushroom body-specific C-term-GFP rescue after Unc13A downregulation via locally driven RNAi expression (Ok107::UAS-C-term-GFP::Unc13A^RNAi^) compared to controls expressing the driver, but not the RNAi (Ok107::+), the RNAi without driver (UAS-Unc13A^RNAi^::+) or mushroom body specific overexpression of the C-term-GFP construct (Ok107::UAS-C-term-GFP). See also Supplementary Fig. 9. Exact values, detailed statistics including sample sizes and P values are listed in Table S1. Scale bars: (a) 20 µm; (b) 10 µm. Statistics: nonparametric one-way analysis of variance (ANOVA) test, followed by a Tukey’s multiple comparison test. ***P ≤ 0.001; n.s., not significant, P > 0.05. All panels show mean ± s.e.m.. For representative images experiments were repeated twice with at least 6-7 brains per genotype.

Because the C-term-GFP construct largely rescued the detrimental effect of *unc13* null mutation at the NMJ^3^, we predicted that general transmission from KCs should be functional for both the C-term-GFP and the C-term-GFP/Unc13A-RNAi conditions. To verify this, we performed *in vivo* two photon Ca^2+^-imaging (GCaMP6f) experiments in adult flies to assess odor-evoked responses at the M4/6 (MBON-β’2mp/MBON-γ5β’2a/MBON-β2β’2a) mushroom body output neurons, a postsynaptic circuit element directly downstream of KCs^57, 58^. Indeed, robust Ca^2+^ transients in response to odor stimulation were observed in all genetic constellations tested (Fig. 7b). We thus conclude that KC output synapses expressing C-term-GFP or C-term-GFP/Unc13A-RNAi are functional under naive conditions. Together with the finding that naive odor avoidance was not statistically different between all these groups (Supplementary Fig. 9b), this allowed us to test whether either condition would interfere with learning and memory.

We assessed short-term memory in adult *Drosophila* using classical aversive olfactory conditioning. Flies were trained by pairing an odor with an electric shock and learning was scored by subsequently assaying avoidance of that odor^59^. All control groups showed similarly robust memory performance, while the relevant C-term-GFP/Unc13A-RNAi mutant (where endogenous Unc13A is knocked down and replaced by the C-term mutant incapable of PHP (Fig. 6e-i)) showed severe short-term memory impairments (Fig. 7c). Whether these impairments are indeed a consequence of the loss of a similar plasticity mechanism as the one observed at the NMJ or whether they are also related to differences in the synaptic transmission profile (e.g. short-term plasticity which is also affected in this mutant^3^) remains to be established. Nevertheless, our data clearly indicate that Unc13A is a target for similar forms of plasticity (Unc13A-RNAi knockdown also impaired short-term memory, Supplementary Fig. 9b,c). Thus, our data imply that structural, functional, and behavioral adaptations are linked, that different forms of presynaptic plasticity may converge on Unc13A, and that multiple forms may operate via conserved mechanisms.

## Discussion

Synapses are able to modify their transmission strength by undergoing plastic changes. This synaptic plasticity is crucial for neuronal circuit adaptation including learning and memory processes^60, 61^. Molecular mechanisms for postsynaptic plasticity have been defined in considerable detail^2^. However, presynaptic mechanisms also modulate transmission strength in many synapse types and species^13, 55, 62^. Homeostatic plasticity is a well-studied form of presynaptic plasticity at the *Drosophila* NMJ where an enhancement of AP-evoked neurotransmitter release counterbalances decreased postsynaptic receptor sensitivity. A number of relevant signaling molecules and pathways, including BMP signaling, CaMKII signaling, TOR signaling, proteasomal degradation and trans-synaptic signaling are required for this^13, 14, 63-65^. These factors appear to converge on two principal avenues to enhance presynaptic transmitter release, via increased Ca^2+^ channel amounts and AP-induced Ca^2+^ influx^46-48^ and secondly via an increase in the number of releasable SVs and their associated release sites^17, 18, 48^. However, some conditions were observed where Ca^2+^ influx or Ca^2+^-channel levels were increased but the quantal content was not (Fig. 6a-d and ^17^), suggesting that release site addition or activation is a required contributor. On the minutes’ time-scale, structural AZ-remodeling was observed, yet whether and how this contributes to the enhancement of NT-release remained unclear^18^.

In the present study, we uncover a presynaptic sequence of molecular events that mediate AZ-remodeling (Fig. 8). We identify the presynaptic cytomatrix as a highly dynamic structure that can add discrete nano-modules of core proteins within minutes. In the initial phase of this structural remodeling, RBP and BRP are needed as their loss occludes an increase in Syx-1A- and Unc13A levels (Fig. 4). Somewhat unexpectedly, although AZ-remodeling occurs on the same minutes’ time-scale, it is dispensable for the rapid potentiation of NT-release during PhTx-induced PHP (e.g. *brp*^Null^, *aplip-1*^ek^^4^ or *srpk79D*^ATC^)(Fig. 6). However, the remodeling is essential for the long-term consolidation of the potentiation (the capacity to restore the AP-evoked response in *gluRIIA*^Null^ mutants was severely impaired when combined with *brp*^Null^ or *srpk79D*^ATC^)(Fig. 6).

**Figure 8:**
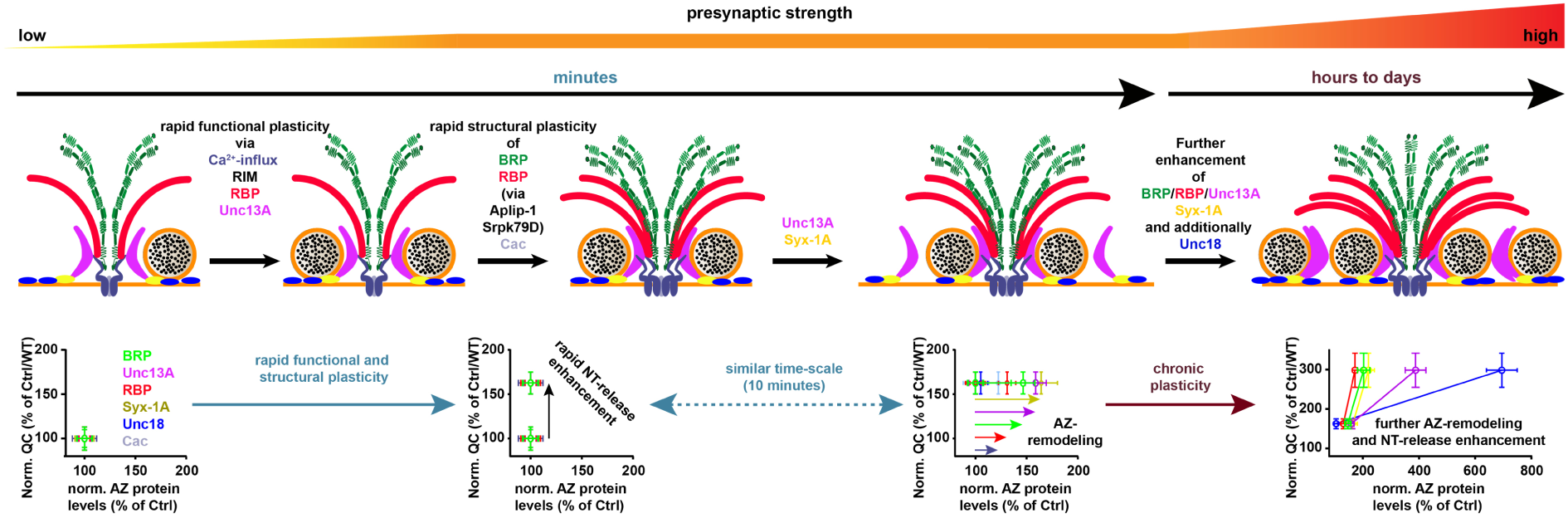
Sequence of events enabling rapid and sustained homeostatic plasticity. Top row: Illustration of AZ modes addressed. Bottom row: Plot of normalized quantal content (QC) vs. AZ protein levels of experiments performed in Fig. 1 normalized to either Ctrl (-PhTx) for rapid plasticity or Wild-type for chronic plasticity. In the basal activity mode (left), BRP (green), RBP (red), Syx-1A (yellow), Unc18 (blue) and Unc13A (magenta) provide two SV release sites at the Ca^2+^-channel (Cac; light blue). However just one release site is active (occupied by SV). (Second left) During the rapid functional plasticity phase the quantal content (and thus neurotransmitter (NT) release) is rapidly enhanced within minutes via mechanisms involving altered Ca^2+^-influx as well as RIM, RBP and Unc13A. On a comparable time-scale (minutes), BRP and RBP are incorporated in a pre-existing AZ in an Aplip-1/Srpk79D dependent manner and additionally Cac-levels also increase (third cartoon). The BRP/RBP incorporation enhances AZ levels of Unc13A/Syx-1A providing an additional release sites (fourth left). This rapid structural AZ-remodeling is not required for the rapid functional plasticity but directly acts on the consolidation of the release enhancement. (Right) On longer time-scales, chronic plasticity then further enhances the AZ-levels of BRP, RBP, Syx-1A in a conserved stoichiometry while Unc13A and Unc18 increase out of scale increasing the number of release sites and thus transmitter release/quantal content activity even further.

Thus, the rapid release enhancement appears to capitalize on AZ-material already present, for example by increasing Ca^2+^ influx^46^ and by the activation of already present but dormant release sites consistent with PHP depending on RIM, RBP and Unc13A ({Muller, 2015 #15;Muller, 2012 #16}; this study). Interestingly, the rapid potentiation coincides with the accumulation of BRP, Cac, RBP, Unc13A, Syx-1A and (later) Unc18 in the AZ, which is required to consolidate and possibly extend the release enhancement. Thus, the synapse utilizes two coincident programs which together ensure immediate rescue and also supply the synapse with backup-material in the form of BRP/RBP/Unc13A nano-modules in case the disturbance persists.

Notably, recent work using STED microscopy to characterize hippocampal synapses also identified AZ nano-modules by clusters of Bassoon, vGlut and Synaptophysin, and observed a scaling of vGlut and Synaptophysin upon chemically induced LTP^66^. This nicely aligns with the structural AZ-remodeling described here, suggesting an evolutionarily conserved process, tuning synaptic transmission by adding nano-modular structures to both sides of the synapse. Additionally, activity dependent alterations in Syx-1A nano-clusters were also recently described^67^, further pointing to the AZ being a highly dynamic structure which adapts to different environmental demands.

How does the AZ scaffold remodel within minutes? While local protein translation is required for some forms of plasticity^68^, acute translation block did not interfere with PhTx induced AZ-remodeling (Supplementary Fig. 5a). However, we found evidence that effective AZ protein transport and a functional cytoskeleton is a precondition here: The BRP/RBP transport adaptor/regulator proteins Aplip-1 and Srpk79D were required for rapid enhancement of BRP/RBP AZ levels (and loss of Aplip-1-mediated BRP/RBP transport impaired short-term memory (Supplementary Fig. 9c)). Moreover, acute actin-depolymerization prevented the PhTx-induced BRP/Unc13A addition into AZs, further supporting a crucial role for their transport and in line with a recent study where *Drosophila* Mical, a highly conserved, multi-domain cytoplasmic protein that mediates actin depolymerization, was shown to be necessary for PHP^63^.

Considering the short timeframe of this adaptation, long-range transport appears unlikely. Instead, we favor the idea that Aplip-1 and Srpk79D function to engage an AZ-proximal reserve pool of components for rapid integration. This pool could originate from a local reservoir in the distal axon or terminal, or even between AZs, from which plasticity may trigger integration into established AZs^18^. Transport processes may fill or empty the reservoir. Between AZs, the reservoir could be composed of diffusely distributed proteins falling below the detection limit^69^, or could reflect a local rearrangement of material (note the reduction of AZs containing few AZ-protein modules after PhTx treatment (Fig. 2c-e) or upon *gluRIIA* ablation (Supplementary Fig. 2a)). Reducing the amount of BRP (by removing one gene copy) blocked the rapid structural adaptation (Fig. 4a,b), possibly because all available material was required to build AZs of proper functionality leaving no material for the reservoir. Regardless of the specific molecular mechanism, guided active transport along the cellular cytoskeleton or rearrangements of the cytoskeleton itself appear to serve a general function in synaptic plasticity in multiple species^63, 70-72^.

RIM and RBP are established targets for multiple forms of presynaptic plasticity in several synapse types and species^5, 16, 17, 62, 73^. Here we additionally identified a critical role for Unc13A (Fig. 6). In fact, with our data we can infer the inter-relation between these factors in presynaptic plasticity. RIM proteins are known to activate (M)Unc13s in several species^49, 50, 53, 74-76^. While *C*. *elegans* and mouse (M)Unc13 proteins interact via an N-terminal C2A domain with RIM, no such domain is known for *Drosophila*, but the principal functional interaction may well be conserved (via another region of the N-term). This was directly tested by an Unc13A mutant, whose N-term was deleted^3^. This mutant largely rescued the severe loss of synaptic transmission in the *unc13*^Null^ condition with comparable AP-evoked transmission as the full-length Unc13A rescue (Compare Fig. 6h black with grey traces). However, this mutant completely lacked the capacity to undergo PHP, consistent with a required RIM/RBP/Unc13A interplay for plasticity. Furthermore, expressing the same mutant in the *Drosophila* mushroom body memory center severely impaired short-term memory formation (Fig. 7), pointing to a relevance of the RIM/RBP/Unc13 plasticity module in the *Drosophila* central nervous system.

Thus, the morphological and molecular similarities between long-term sensitization in *Aplysia*, presynaptic LTP in the mammalian brain and homeostatic plasticity or learning in *Drosophila* indicate that the sequence of molecular events we describe here might be highly conserved.

## Supporting information

## Online Methods

### Experimental Model and Subject Details

#### Fly husbandry, stocks and handling

Fly strains were reared under standard laboratory conditions^77^ and raised at 25°C on semidefined medium (Bloomington recipe). For RNAi experiments flies and larvae were kept at 29°C. For experiments both male and female 3^rd^ instar larvae or flies were used. The following genotypes were used: Wild type: +/+ (*w*^*1118*^). *gluRIIA*^Null^: *df(2L)cl*^*h4*^/*df(2L)gluRIIA&IIB^*SP22*^* (A22);*GluRIIB-GFP/+* or *AD9/df(2L)cl^*h4*^ or gluRIIA*^*SP16*^/ *gluRIIA*^*SP16*^. Supplementary Fig. 1a: Ok6::Syd-1-GFP: *Ok6-Gal4/+;UAS-Syd-1-GFP/+*. Transport mutants: Fig. 3,5; Supplementary Fig. 7: *aplip-1*^Null^: *aplip-1*^*ex213*^/Df(3L)BSC799; *aplip-1*^*ek4*^: *aplip-1*^ek4^/Df(3L)BSC799; *atg-1: atg1*^*ey07351*^/*Df(3L)BSC10*; *srpk79D*^*ATC*^: *srpk79D*^*atc*^*/*srpk79D^*atc*^. Supplementary Fig. 5c: Ctrl: *Ok6-Gal4/+;UAS-Unc13A-GFP/+*; *aplip-1-RNAi*: *Ok6-Gal4/+;UAS-Unc13A-GFP/UAS-Aplip-1-RNAi;* Supplementary Fig. 5d; Ctrl: *Ok6-Gal4/+;UAS-Unc13A-GFP/+*; srpk79D-RNAi: *Ok6-Gal4/+;UAS-Unc13A-GFP/UAS-srpk79D-RNAi*. Intravital imaging: Movie 1 and Supplementary Fig. 5e-g: Ok6/+;UAS-Unc13A-GFP/UAS-BRP-D3-Straw. Fig. 4,5: *rbp*^Null^: *rbp*^*Stop1*^/*rbp*^*S*2^.*^01^*; *brp*^Null^: *brp*^*Δ6*^.*^1^*/*brp*^*69*^; *brp*^Null^/+: *brp*^*69*^/*+; rim*^Null^: *rim*^*ex*1^.*^103^*/*Df(3R)ED5785*; *rim*^Null^,*fife*^Null^: *rim*^*ex*1^.*^103^*,*fife*^*ex1027*^/*rim*^*ex1*^.*^103^*,*fife*^*ex1027*^; *liprin-α*^Null^: *liprin-α*^*F3ex15*^/*liprin-α*^*R60*^; *syd-1*^Null^: *syd-1*^*1.2*^/*syd-1*^*3*.*4*^. Fig. 5: *gluRIIA*^Null^,*brp*^Null^: *gluRIIA*^SP16^,*brp*^6^.^1^/*gluRIIA*^SP16^,*brp*^69^; *gluRIIA*^Null^;*srpk79D*^ATC^: AD9/*df(2L)cl*^*h4*^; *srpk79D*^ATC^ /*srpk79D*^ATC^. Supplementary Fig. 7: *gluRIIA*^Null^;*aplip-1*^ek4^: AD9/*df(2L)cl*^*h4*^;*aplip-1*^ek4^/*Df(3L)BSC799*. Fig. 6; Supplementary Fig. 8: *unc13A*^Null^: *EMS7*.*5/P84200; For UAS-Cac-GFP in unc13A*^Null^: *Ctrl: Ok6-Gal4, UAS-Cac-GFP/+*; *unc13A*^Null^: *Ok6-Gal4, UAS-Cac-GFP/+;;EMS7*.*5/P84200;* Unc13A-GFP: *elav-GAL4/+;;UAS-Unc13A-GFP/+;P84200/P84200;* C-term-GFP: *elav-GAL4/+;;UAS-C-term-GFP/+;P84200/P84200*. Learning and memory: Fig. 7; Supplementary figure 9: Ok107::+: *Ok107-Gal4/+*; Ok107::Aplip-1^RNAi^: *UAS-Aplip-1-RNAi/+;Ok107-Gal4/+*; MB247::+: *MB247-Gal4/+*; MB247::Aplip-1-RNAi: *MB247-Gal4/UAS-Aplip-1-RNAi*; UAS-Unc13A^RNAi^::+: *UAS-Unc13A-RNAi/+*; Ok107::UAS-C-term-GFP: *UAS-C-term-GFP/+; Ok107-Gal4/+*: OK107::UAS-C-term-GFP::Unc13A^RNAi^: *UAS-C-term-GFP/UAS-Unc13A-RNAi; Ok107-Gal4/+;* Ok107::UAS-Unc13A^RNAi^: *UAS-Unc13A-RNAi/+;Ok107-Gal4/+*. Fig. 7b: Ok107::+: *VT1211-LexA::LexAop-GCaMP6f/+;;Ok107-Gal4/+*; Ok107::UAS-C-term-GFP: *VT1211-LexA::LexAop-GCaMP6f/+;UAS-C-term-GFP/+;Ok107-Gal4/+*; OK107::UAS-C-term-GFP::Unc13A^RNAi^: *VT1211-LexA::LexAop-GCaMP6f/+; UAS-C-term-GFP/UAS-Unc13A-RNAi; Ok107-Gal4/+*. Western blot, Supplementary Fig. 9a: elav::+: *elav-Gal4/+*; UAS-Unc13A-RNAi::+: *UAS-Unc13A-RNAi/+*; elav::UAS-Unc13A^RNAi^: *elav-Gal4/+;;UAS-Unc13A-RNAi/+*. Stocks were obtained from: A22 ^23^; AD9, df(2L)cl^h4^, *gluRIIA*^SP16^ ^22^; GluRIIB-GFP ^78^; *Ok6-GAL4* ^*79*^; *rbp*^*Stop1*^, *rbp*^*S2.01*^ ^9^; *rim*^*ex1.103*^ ^17^; *fife*^ex1027 34^; *liprin-α*^*F3ex15*^, *liprin-α*^*R60*^ ^39^; *UAS-Syd-1-GFP*; *syd-1*^*1*^.*^2^*, *syd-1*^*^3^*.*^4^* 41^; *brp*^*Δ*6^.*^1^* ^38^; *brp*^*69*^ ^8^; EMS7.5, *UAS-Unc13A-GFP* ^7^; UAS-BRP-D3-Straw ^78^; UAS-Cac-GFP ^80^; *elav-Gal4* ^*81*^; *UAS-Unc13A-RNAi, UAS-C-term-GFP* ^3^; *aplip-1*^*ex213*^ ^*32*^; *aplip-1*^*ek4*^ ^33^; *srpk79D*^*atc*^ ^29^; *Ok107-Gal4* ^82^; *MB247-Gal4* ^*83*^. *P84200* was provided by the Drosophila Genetic Resource Center (DGRC). The *aplip-1*^*ek4*^; Df(3L)BSC799; *atg1*^ey07351^; Df(3L)BSC10; Df(3R)ED5785 lines were provided by the Bloomington *Drosophila* Stock Center. *UAS-Aplip-1-RNAi* and *UAS-srpk79D-RNAi* from VDRC.

### Method Details

#### Immunostaining

Larvae were dissected and stained as described previously^41^. The following primary antibodies were used: guinea-pig Unc13A (1:500;^7^); mouse Syx1A 8C3 (1:40; Developmental Studies Hybridoma Bank, University of Iowa, Iowa City, IA, USA; AB Registry ID: AB_528484); mouse Unc18/Rop 4F8 (1:500; Developmental Studies Hybridoma Bank, University of Iowa, Iowa City, IA, USA; AB Registry ID: AB_1157869; mouse GFP 3E6 (1:500, Thermo Fisher Scientific Inc., MA, USA, A-11120; AB Registry ID: AB_221568), mouse Nc82 = anti-BRP^C-term^ (1:100, Developmental Studies Hybridoma Bank, University of Iowa, Iowa City, IA, USA; AB Registry ID: AB_2314865); rabbit BRP^Last200^ (1:1000;^84^); rabbit RBP^C-term^ (1:500;^9^); guinea-pig vGlut; (1:2000;^85^). Except for staining against RBP; Syx1A and Unc18, where larvae were fixed for 10 min with 4 % paraformaldehyde (PFA) in 0.1 mM phosphate buffered saline (PBS), all fixations were performed for 5 min with ice-cold methanol. The glutamate receptor blocker PhTx-433 (Sigmal-Aldrich, MO, USA) was prepared as a 4 mM stock solution either in DMSO (final DMSO concentration, 0.5%) or dH_2_O. Rapid pharmacological homeostatic challenge was assessed by incubating semi-intact preparations in 20 µM PhTx diluted in HL3 (see below) containing 0 or 1.5 mM CaCl_2_ for 10 min at room temperature^19^. Controls were treated in the same way but were incubated either in pure HL3 or HL3 containing 0.5% DMSO (in dependence of how PhTx stock solution was prepared) for 10 min. After incubation, the dissection was completed and the preparation was rinsed three times with fixative solution. During the dissection, extreme care was taken to avoid excessive stretching of body wall muscles, as this may significantly impair induction of homeostasis^19^.

For translation block experiments semi-intact preparations were pre-incubated in HL3 containing 50µg/ml Cycloheximide (Chx; Sigmal-Aldrich, MO, USA) dissolved in DMSO for 10 minutes, a concentration previously shown to block protein synthesis in Drosophila ^86^. Rapid pharmacological homeostatic challenge was then assessed by incubating semi-intact preparations in 20 µM PhTx (or a similar volume of H_2_O in control experiments) diluted in HL3 (see below) containing 50µg/ml Chx for 10 min at room temperature. For actin-depolymerization experiments semi-intact preparations were pre-incubated in HL3 containing 15 µM Latrunculin B (Abcam, UK) in DMSO for 10 minutes. For control experiments larvae were incubated with a solution containing a similar volume of DMSO. Rapid pharmacological homeostatic challenge was then assessed by incubating semi-intact preparations in 20 µM PhTx (or a similar volume of H_2_O in control experiments) diluted in HL3 (see below) containing 15 µM Latrunculin B in DMSO (or DMSO alone in control experiments) for 10 min at room temperature. Afterwards, prepping and staining procedures were performed as described above/below. Control animals were always reared in parallel and treated identically in all experiments.

Secondary antibodies for standard immunostainings were used in the following concentrations: goat anti-HRP-Cy5 (1:250, Jackson ImmunoResearch, PA, USA); goat anti-HRP-647 (1:500, Jackson ImmunoResearch 123-605-021, PA, USA); goat anti-rabbit-Cy3 (1:500, Jackson ImmunoResearch 111-165-144, PA, USA); goat anti-mouse-Cy3 (1:500, Jackson ImmunoResearch 115-165-146); goat anti-mouse or anti guinea pig Alexa-Fluor-488 (1:500, Life Technologies A11001/A11073, CA, USA). Larvae were mounted in vectashield (Vector labs, CA, USA). Secondary antibodies for STED were used in the following concentrations: goat anti-mouse or rabbit Alexa594 (1:500, Thermo Fisher Scientific Inc. A11032/A11037, MA, USA); goat anti-mouse Atto590 (1:100); goat anti-rabbit Atto590 (1:100); goat anti-guinea pig star635 (1:100); goat anti-rabbit star635 (1:100); goat anti-mouse or rabbit Atto647N (1:250; Active Motif; 15038/15048). Atto590 (ATTO-TEC AD 590-31) and star635 (Abberior 1-0101002-1) coupled to respective IgGs (Dianova). For STED imaging larvae were mounted in Mowiol (Max-Planck Institute for Biophysical Chemistry, Group of Stefan Hell) or ProLong Gold (Life-Technologies, CA, USA) on high-precision glass coverslips.

Western blot analysis was done as previously described^87^. Briefly, adult flies were dissected in cold Ringer’s solution and homogenized in Lysis buffer (1x PBS, 0.5%Triton, 2%SDS, 1x Protease inhibitor, 1x Sample buffer) followed by full-speed centrifugation at 18°C. One brain’s supernatant for each group was subjected to SDS-PAGE and immunoblotted according to standard procedures. The following antibodies were used: guinea-pig Unc13A (1:2000;^7^) and mouse Tubulin (1:100000; Sigmal-Aldrich, MO, USA; Cat# T9026, AB Registry ID: AB_477593). Antibodies obtained from the Developmental Studies Hybridoma Bank were created by the NICHD of the NIH and maintained at The University of Iowa, Department of Biology, Iowa City, IA 52242.

#### Image Acquisition, Processing, and Analysis

Confocal microscopy was performed with a Leica SP8 microscope (Leica Microsystems, Germany). Images of fixed and live samples were acquired at room temperature. Confocal imaging of NMJs was done using a z-step of 0.25 µm. The following objective was used: 63×1.4 NA oil immersion for NMJ confocal imaging. All confocal images were acquired using the LAS X software (Leica Microsystems, Germany). Images from fixed samples were taken from muscle 4 of 3^rd^ instar larval 1b NMJs (segments A2-A5) or nerve bundles (segments A1–A3). Images for figures were processed with ImageJ software to enhance brightness using the brightness/contrast function. If necessary, images were smoothened (0.5 pixel Sigma radius) using the Gaussian blur function. Confocal stacks were processed with ImageJ software (http://rsbweb.nih.gov/ij/). Quantifications of AZs (scored via BRP) were performed following an adjusted manual ^88^, briefly as follows. The signal of a HRP-Cy5 antibody was used as template for a mask, restricting the quantified area to the shape of the NMJ. The original confocal stacks were converted to maximal projections, and after background subtraction, a mask of the synaptic area was created by applying a certain threshold to remove the irrelevant lower intensity pixels. The segmentation of single spots was done semi-automatically via the command “Find Maxima” embedded in the ImageJ software and by hand with the pencil tool and a line thickness of 1 pixel. To remove high frequency noise a Gaussian blur filter (0.5 pixel Sigma radius) was applied. The processed picture was then transformed into a binary mask using the same lower threshold value as in the first step. This binary mask was then projected onto the original unmodified image using the “min” operation from the ImageJ image calculator. For spots/µm^2^ the number of spots was divided by the size of the mask of the synaptic area.

For colocalization analysis (Pearson correlation coefficient) the ImageJ plugin “JACOP” (http://rsb.info.nih.gov/ij/plugins/track/jacop2.html) was used. To determine the synaptic protein levels, a custom-written ImageJ script was used that detects the locations with highest local maxima in pixel values to generate regions of interests (ROIs) and sets a point selection at each. The intensities were then measured and all selections were deleted, leaving intensity values and (x,y) locations in the results list. The results list was then used to create a circle of size = 5 pixels (pixel size 100 nm) centered around each (x,y) location and the integrated density within these ROIs was measured and taken for further calculations. The same ROIs were then used in the channel containing signals of the co-stained protein. For scatter plots, co-stainings of BRP either with RBP, Unc13A or Syx-1A with or without PhTx-treatment or Wild-type and *gluRIIA*^Null^ were used. The AZ numbers were counted (number of BRP spots) and the local synaptic levels of both co-stained proteins were measured. AZs were then sorted into five bins (AZ number divided by five) depending on their synaptic BRP levels and then the respective second channel intensities distributed to the appropriate bin. Binned BRP levels were then plotted against binned levels of the second channel.

#### STED Microscopy

Two-color STED images were recorded on custom-built STED-microscopes^89, 90^, which either combine two pairs of excitation laser beams of 595 nm and 635 nm or 595 nm and 640 nm wavelength with one STED fiber laser beam at 775 nm. All STED images were acquired using Imspector Software (Max Planck Innovation GmbH, Germany). STED images were processed using a linear deconvolution function integrated into Imspector Software (Max Planck Innovation GmbH, Germany). Regularization parameters ranged from 1e^−09^ to 1e^−10^. The point spread function (PSF) for deconvolution was generated by using a 2D Lorentz function with its half-width and half-length fitted to the half-width and half-length of each individual image. For Fig. 3C, dual-color STED imaging with time-gated detection was performed using a commercial Leica SP8 TCS STED microscope (Leica Microsystems, Germany) equipped with a 100x NA 1.4 objective (HC PL Apo CS2; Leica Microsystems, Germany). Briefly, the system includes an inverted DMi8 CS microscope equipped with a 100x pulsed white light laser (WLL; ∼80-ps pulse width, 80-MHz repetition rate; NKT Photonics, Denmark) with a STED lasers for depletion (pulsed) at 775 nm. Detection of Alexa594 after excitation at 594 nm and emission detection of 604-650 nm and Atto647 after excitation at 640 nm at emission of 656 – 751 nm was performed in frame sequential mode. Time-gated detection with Hybrid detectors was set from 0.3–6 ns for both dyes. Raw data were deconvolved with Huygens Professional software (Scientific Volume Imaging) using a theoretical point spread function automatically computed based on pulsed-wave STED optimized function and the specific microscope parameters. Default deconvolution settings were applied. Images for figures and for finding high-intensity clusters (see below) were processed with ImageJ software to remove obvious background or neighboring AZs (if required), enhance brightness/contrast and smoothened (0.5 pixel Sigma radius) using the Gauss blur function.

#### Classification and Alignment of Single AZs

All analysis described below was done using MATLAB R2016b (Mathworks Inc., MA, USA) with optional toolboxes, which are indicated in the respective sections. Classification of individual AZs was achieved by using a custom script to detect the position of cluster centers (local intensity maxima) in the images where obvious background was removed (only for Unc13A, BRP, RBP), while the averaging procedure described hereafter was performed on unretouched images. This code retains only pixels above a defined grey value threshold. For the analysis of Unc13A stainings shown in Fig. 2f, a threshold value of 25 was used (18 for BRP, RBP, Syx-1A and Unc-18). All pixel values in the image below this threshold value were set to zero to remove background noise. We then identified the positions of high-intensity pixel clusters as local intensity maxima in the images freed from neighboring AZ signal and obvious noise. This was achieved by finding local maxima in the vertical and horizontal pixel lines. First, the function searched the first derivative (using the function *diff*) of all pixel columns for zero values and the changes in the surrounding slopes were considered to identify local maxima. The same procedure was then applied in pixel rows, but only for those pixel column values associated with a local maximum in the previous step. All single pixels that were associated with a maximum in both a row and a column were detected using the function *intersect*. To prevent detection of the same cluster more than once, a defined minimum distance of clusters was used (50 nm for BRP, Unc13A, Syx-1A and RBP, 20 nm for RBP) and only the local maximum with the highest intensity value was considered. All subsequent translation and averaging procedures were performed on the non-cleaned images of the same AZs using the determined classification and positions of clusters. The non-cleaned AZs were sorted by the number of protein clusters detected this way. To calculate the center of mass of all coordinates, the means of all x- and y-coordinates were taken according to equations (1) and (2).

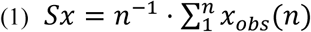

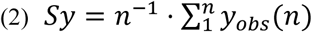

Where (Sx,Sy) is the (x,y)-coordinate center of mass in the initial image, n is the number of identified clusters, and x_obs_(n) and y_obs_(n) are the positions of the n^th^ cluster in the present image. To align the center of mass to the center of the image, the necessary shift (Δx and Δy) of the original coordinates was calculated according to equations (3) and (4) and used subsequently in equations (5) and (6).

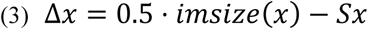

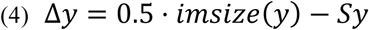

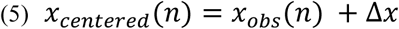

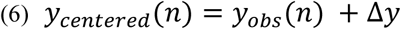

In equations (3) and (4), “imsize” refers to the size of the image in x or y dimension. The resulting coordinates x_centered_ and y_centered_ represent cluster coordinates after shifting the original coordinates x_obs_ and y_obs_. The same translation was applied to the corresponding AZ image (using the function *imtranslate*, part of the ‘Image Processing’ toolbox). Clusters were ranked in a counter-clockwise sequence in all images (see illustration in Fig. 2b; Supplementary Fig. 3c) by sorting them for increasing angle between the image center and the cluster location in relation to the vertical midline (x=26).

To align protein cluster coordinates between all investigated AZs, central rotation (around the image center, x_middle_=y_middle_=26), was done using the operation in equation (7).

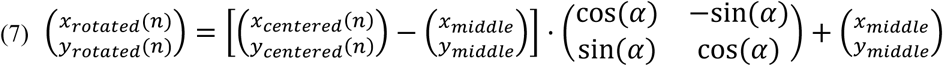

To find the optimal angle to overlay all AZs, a cost reflecting the sum of distances between cluster positions of the same rank in all images was minimized. The cost function was defined as described in equation (8).

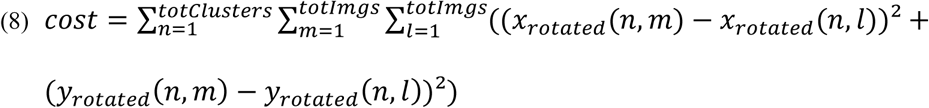

In equation (8), *n* is the cluster number, *totClusters* is the total cluster number of the respective category of images, *m* and *l* are particular AZ images in the stack of images from one category, and *totImgs* is the total number of AZ images in that category. The squared Euclidean distances were found by using the function *pdist*.

The optimal rotation angle was found using a genetic algorithm function (*ga*, part of the ‘Global Optimization’ toolbox). The rotation angles evaluated were constrained in a range from - 180 to 180 degrees. For faster optimization, parallelization (setting the option ‘UseParallel’ to ‘true’; this requires the ‘Parallel Computing’ toolbox) was employed to evaluate 500 individual cost functions per generation. Cluster coordinates from all images in one category were rotated simultaneously. A single individual in the genetic algorithm represented a set of rotation angles for each image. The convergence criterion (*TolFun*) was left at the default value (a relative cost value change of less than 10^-6^ over 50 generations). The output of this optimization was a vector containing all rotation angles that led to the best overlap of cluster coordinates. Finally, these rotations were then applied to the centered original AZ images. All images aligned this way were then combined in a stack and an average image was generated by calculating the mean intensity of all image pixels. For better illustration of the AZ structure, pixel intensities were linearly scaled such that the highest intensity pixel had a value of 255. The procedure was only performed if more than two images existed in the same cluster number class for at least 5 consecutive cluster number classes. The histograms shown in Fig. 2c,d,e and supplementary Fig. 2a were generated by counting the number of images in each cluster number class, and dividing each value by the total amount of images detected in all classes. The mean cluster intensity was calculated by taking the differences between the mean image intensities in consecutive classes, and averaging them.

To investigate the AZ structure in an approach independent of the cluster-distance minimization procedure described above, we repeated the averaging in a different way as follows. We developed a MATLAB code for AZ centering and alignment by rotation of the highest intensity pixel to identical angles and therefore similar positions, which yielded qualitatively similar results (Supplementary Fig. 3a). Again, high-intensity clusters were found in AZ images cleaned from obvious noise and bordering AZs, and all translations and rotations were then performed on unretouched images. The center of mass of the found cluster coordinates was calculated and the image shifted so that the center of mass of the coordinates was in the center of the 51 by 51 pixel space (x=y=26; see equations (1) to (6)). The brightest point in each image was then found by sorting the list of intensities of all clusters found. Only this point was then considered for rotation. To determine the angle by which to rotate each image to place this highest intensity region to the same fixed position (on the vertical midline between the two top image quadrants), the x and y distances to the center of rotation (equal to the center pixel of the image x=y=26) were calculated to find the length (*l*) of the hypotenuse and the opposite side of the right triangle. The angle α in degrees was then calculated by taking the inverse sine in degrees of this value (MATLAB function *asind*), as shown in equation (9).

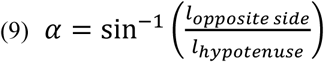

In cases where the brightest peak was located above the horizontal midline, the adjacent side of the triangle was the vertical midline. In cases where the brightest peak was located below the horizontal midline, the angle was calculated with the horizontal midline being the adjacent side, and 90° were added to the final angle value. Additionally, in cases where the brightest peak was located to the right of the vertical midline, the angle was multiplied with -1. To generate the results shown in Supplementary Fig. 3b, which shows the averaging of AZ images within randomly assigned categories, we generated a number of random category values and then proceeded with the averaging procedure described above. For this, we reproduced the distribution of category values from the AZ dataset according to the histogram values of the category vector as follows. A histogram of the category vector was generated (MATLAB function *histogram*) with a bin width of 1, yielding the absolute amount of AZs per category. A cumulative sum vector was calculated from these histogram values (MATLAB function *cumsum)*. For each position in the category vector, we then chose a random number between 0 and 1 and multiplied it by the number of images considered in the category vector. We then found the first position in the cumulative sum vector that was larger than this random value.

The position found was equal to the assigned category. This resulted in a randomly assigned category vector with a similar distribution of categories as the original vector.

#### *In vivo* live imaging and analysis

*In vivo* imaging of intact *Drosophila* larvae was performed as previously described^91^. Briefly, 3^rd^ instar larvae were put into a drop of Voltalef H10S oil (Arkema, Inc., France) within an airtight imaging chamber. Before imaging, the larvae were anaesthetized with 20 short pulses of a desflurane (Baxter,IL, USA) air mixture until the heartbeat completely stopped. For assessing axonal transport, axons immediately after exiting the ventral nerve cord were imaged for 5 min using timelapse confocal microscopy. Kymographs were plotted using a custom-written ImageJ script.

#### Dissection, Induction of Homeostatic Plasticity and Electrophysiology

Third-instar larvae were selected and placed individually on a Sylgard block. Using a very sharp pin, the tail of the larva was pinned between the posterior spiracles, in the absence of solution. The head was pinned, making sure not to stretch the larva, so that the animal was relatively loose between the two pins. A small horizontal incision was made in the dorsal cuticle at the tail with a sharp scissors. The larva was cut vertically from the tail incision in an anterior direction (towards the head), continuing beyond the head pin. Great care was taken not to stretch the cuticle or animal during this process. 40 µl of a 20 µM PhTx in modified hemolymph-like solution (HL3; ^92^ composition (in mM): NaCl 70, KCl 5, MgCl_2_ 10, NaHCO_3_ 10, trehalose 5, sucrose 115, HEPES 5, CaCl_2_ 0, pH adjusted to 7.2) was pipetted into the abdominal cavity with minimal force, making sure to fill the abdomen. After 10 minutes incubation, the preparation was completed without rinsing. The cuticle was gently pinned twice on each side (without stretching). The connection of the intestines and trachea to the body at the posterior were cut. Holding the now free ends of the intestines and trachea with a fine forceps, remaining connections were cut moving in an anterior direction (towards the head). The intestines and trachea could then be gently removed without stretching the larva. Finally, the brain was held firmly and slightly raised above the body so that the scissors could be placed underneath to cut the segmental nerves. Care was taken not to touch the underlying muscle and to avoid excessive pulling of the nerves before they were cut. The completed preparation was rinsed 3 times with PhTx-free HL3 (0 mM CaCl_2_, 10 mM MgCl_2_). PhTx-free HL3 solution was used in control treatments. Sylgard blocks were kept separate for PhTx and control treatments and all implements were rinsed after each recording.

The Sylgard block and completed larval preparation was placed in the recording chamber which was filled with 2 ml HL3 (0.4 mM CaCl_2_, 10 mM MgCl_2_). Recordings were performed at room temperature (∼22°C) in current clamp mode at muscle 6 in segments A2/A3 as previously described^93^ using an Axon Digidata 1550A digitizer, Axoclamp 900A amplifier with HS-9A x0.1 headstage (Molecular Devices, CA, USA) and on a BX51WI Olympus microscope with a 40X LUMPlanFL/IR water immersion objective. Sharp intracellular recording electrodes were pulled using a Flaming Brown Model P-97 micropipette puller (Sutter Instrument, CA, USA) with a resistance of 20-35 MΩ, back-filled with 3 M KCl. Cells were only considered with a membrane potential less than -60 mV and membrane resistances greater than 4 MΩ. All recordings were acquired using Clampex software (v10.5) and sampled at 10-50 kHz, filtering with a 5 kHz low-pass filter. mEPSPs were recorded for 1 minute. eEPSPs were recorded by stimulating the appropriate nerve at 0.1 Hz, 5 times (8 V, 300 µs pulse) using an ISO-STIM 01D stimulator (NPI Electronic, Germany). Stimulating suction electrodes were pulled on a DMZ-Universal Puller (Zeitz-Instruments GmbH, Germany) and fire polished using a CPM-2 microforge (ALA Scientific, NY, USA). A maximum of two cells were recorded per animal.

Analysis was performed with Clampfit 10.5 and Graphpad Prism 6 software. mEPSPs were further filtered with a 500 Hz Gaussian low-pass filter. Using a single template for all cells, mEPSPs were identified and analyzed, noting the mean mEPSP amplitude per cell. For Fig. 5 and Supplementary Fig. 7, templates were generated for each cell and the first 30 mEPSPs were identified and taken into account for further analysis. An average trace was generated from the 5 eEPSP traces per cell. The amplitude of the average eEPSP trace was divided by the mean mEPSP amplitude, for each respective cell, to determine the quantal content.

Dissection and current clamp recordings of w1118 vs *gluRIIA*^Null^ were performed as above in male third-instar larvae. Cells with an initial membrane potential greater than -55 mV, resistances less than 5 MΩ or multiple responses to a single stimulus were rejected. eEPSPs were recorded by stimulating the appropriate nerve at 0.2 Hz, 10 times (6 V, 300 µs pulse). An average eEPSP amplitude was calculated from the 10 traces. mEPSPs were analysed with a genotype specific template. Quantal contents were calculated by dividing the mean eEPSP by mean mEPSP for each cell.

#### In Vivo Two-Photon Live Calcium Imaging and Analysis

Two-photon imaging of odor-evoked calcium responses was conducted in 3 to 5-day-old mixed-sex flies expressing LexAop-GCaMP6f in VT1211-LexA. For imaging, flies were briefly anesthetized on ice and mounted in a custom made chamber by immobilizing wings, head and proboscis with wax. The head capsule was opened in sugar-free HL3-like extracellular saline^94^. Odor stimulation consisted of a 1.5 s OCT pulse followed by a 30 s break and then a 1.5 s MCH pulse followed again by another 30 s break. This alternating odor pulse protocol was consecutively repeated five times (odor dilution in mineral oil 1/1000). Odors were delivered on a clean air carrier stream and image acquisition and odor stimulation was synchronized temporally using a custom-designed system. Fluorescence was centered on 910 nm generated by a Ti-Sapphire laser (Chameleon Ultra II, Coherent, CA, USA). Images with a pixel size of 0.3 x 0.3µm were acquired at 70 Hz using two-photon microscopy (Femto2D-Resonant by Femtonics Ltd., Hungary) with a 20X, 1.0 NA water-immersion objective, controlled by MESc v3.5 software (Femtonics Ltd., Hungary). For each animal, a single hemisphere was analyzed. All OCT and MCH responses of a fly were averaged respectively and resulting traces were averaged between flies. Mean intensity values of two-photon fluorescence were calculated, while F0 was defined as the mean F from 0 to 1.5s at the beginning of a recording (MESc v 3.5 software). Image processing for single frames were manually performed using ImageJ (images did not require registration).

#### Odor avoidance conditioning

All flies were 3 to 5 days old, raised in 12 h:12 h, light:dark cycle and at 65% relative humidity. One day before the experiment, the flies were transferred to fresh food vials. One hour prior to the experiment, flies were pre-conditioned to experimental conditions (dim red light, 25°C, humidity of 80%). The aversive odors 3-Octanol (OCT) and Methylcyclohexanol (MCH) were diluted 1:100 in paraffin oil and presented in 14 mm cups. A current of 120V AC was used as a behavioral reinforcer. The associative training was performed as previously described^95^. In a single-cycle training, nearly 100 flies were presented with one odor (CS^+^) paired with electrical shock (US; 12 times for one minute). After one minute of pure air-flow, the second odor was presented without the shock (CS^-^) for another minute. The flies were then immediately tested for short-term memory performance by presenting them the two odors together. A performance index (PI) was calculated as the number of flies choosing the odor without shock (CS^-^), minus the number of flies choosing the odor paired with shock (CS^+^), divided by the total number of flies, multiplied by 100. The values of PI ranges from 0 to 100 where 0 means no learning (50:50 distribution of flies) and a value of 100 means complete learning (all flies avoided the conditioned odor). The final learning index was calculated as the average of both reciprocal indices for the two odors. Odor Avoidance experiments were used to test innate behavior where each odor was presented to the flies without conditioning. The PIs were calculated as stated above.

#### Quantification and Statistical Analysis

Data were analyzed using Prism (GraphPad Software, CA, USA). Per default Student’s t test was performed to compare the means of two groups unless the data were either non-normally distributed (as assessed D’Agostino-Pearson omnibus normality test) or if variances were unequal (assessed by F test) in which case they were compared by a Mann-Whitney U Test. However, in supplementary table 1 both tests are provided for all relevant cases. For comparison of more than two groups, one-way analysis of variance (ANOVA) tests were used, followed by a Tukey’s multiple comparison test. P values and N values are given in Supplementary table 1. Means are annotated ± s.e.m.. Asterisks are used to denote significance: *, p < 0.05; **, p < 0.01; ***, p < 0.001; n.s. (not significant), p > 0.05.

#### Data and Software Availability

The data that support the findings of this study as well as MATLAB and ImageJ codes used in this study are available from Alexander M. Walter upon request.

#### Contact for Reagent and Resource Sharing

Informations and requests for resources and reagents should be directed to and will be fulfilled by the Lead Contact, Alexander M. Walter.

## Acknowledgements

We thank S. Waddell and Y. Huang for the VT1211-LexA fly line. M.A. Böhme was supported by the SFB 740. A.T. Grasskamp was supported by a NeuroCure Ph.D. fellowship funded by the Deutsche Forschungsgemeinschaft (Exc 257) within the International Graduate Program Medical Neurosciences. M. Jusyte was supported by an Einstein Center for Neurosciences Ph.D. fellowship. U. Rey was supported by a fellowship of the International Max Planck Research School by the Max-Planck-Gesellschaft. This work was supported by grants from the Deutsche Forschungsgemeinschaft to S.J. Sigrist (Exc 257, TP A3 and A6 SFB 958, TP B9/SFB665; TP09/SFB740), A.T. Grasskamp (TRR 186), A.M. Walter (Emmy Noether programme, TRR 186), D. Owald (Emmy Noether programme) and from the National Institutes of Health to D. Dickman (NS091546).

## Author Contributions

M.A.B, S.J.S and A.M.W conceived the project. M.A.B., A.W.M., C.B.B., P.G. and A.G.P. performed fly husbandry and maintenance. M.A.B., M.J., M.L. and P.G. performed confocal and/or STED imaging experiments and M.A.B., P.G. and A.T.G. analyzed the data. A.W.M., M.J. and P.G. performed electrophysiological experiments and analyzed the data. U.R. performed live-imaging experiments and analysis. P.H. provided reagents and suggested additional experiments. A.T.G and A.M.W. developed software codes for image alignment and averaging and A.T.G analyzed the data. P.G. and D.D. conceived and performed translation block experiments. C.B.B. performed adult *Drosophila* brain antibody staining and imaging. C.B.B. and S.H. performed behavioral experiments and data analysis. S.H. performed Western blots. D.L. and D.O. planned and performed in vivo two-photon live calcium imaging. F.G. and S.W.H. developed and built the STED microscope. M.A.B., S.J.S, and A.M.W wrote the paper with input from all co-authors.

