## Supplementary material for "Rapid active zone remodeling consolidates presynaptic potentiation"

**Supplemental figures, titles and legends**

**Supplementary Figure 1 – related to Figure 1**

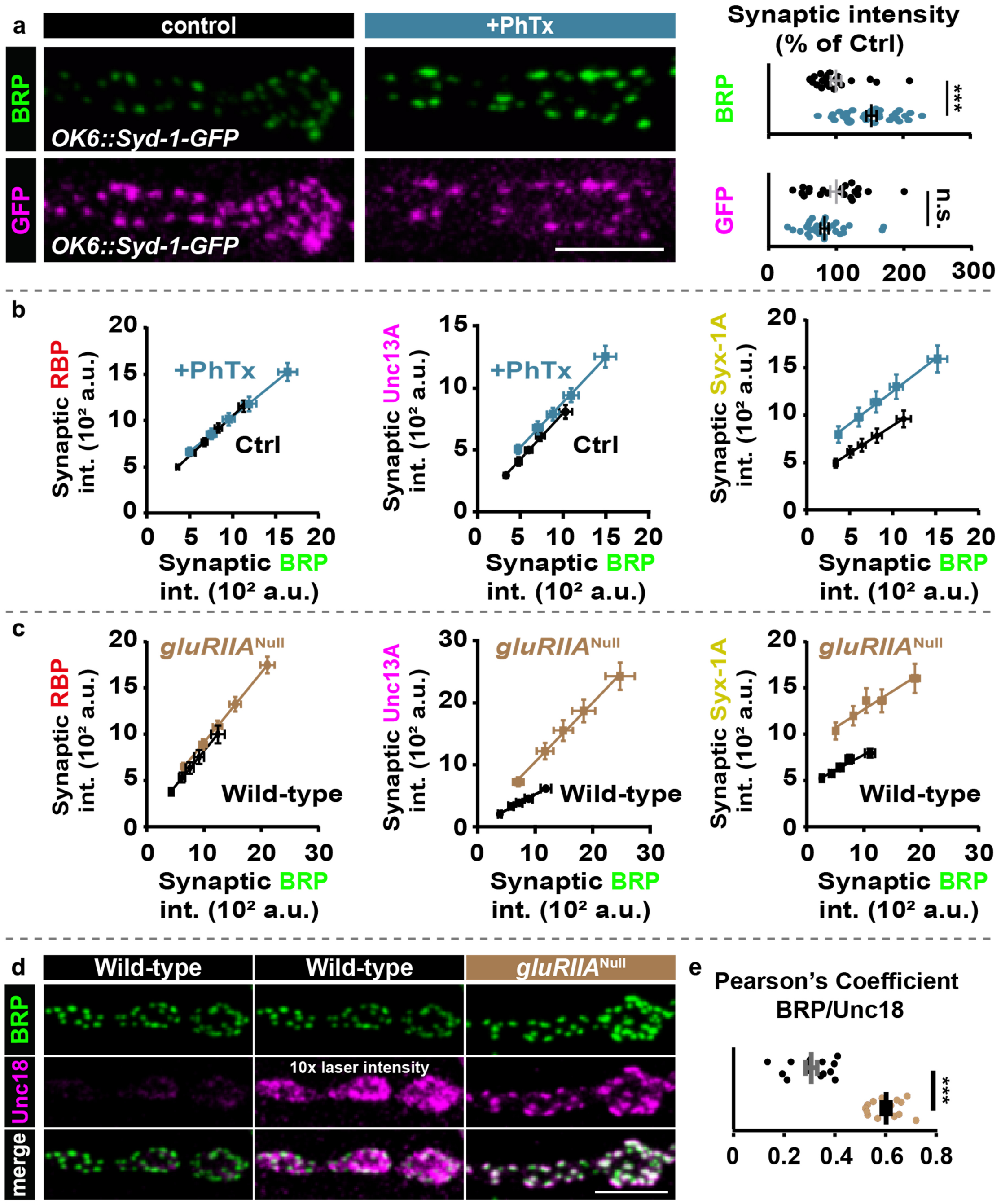


**Supplementary Figure 1: Rapid and chronic homeostatic potentiation enhances AZ protein levels.**

**(a)** Confocal images (left) and quantification (right) of synaptic BRP and GFP AZ-levels in motoneurons overexpressing Syd-1-GFP, labelled with the indicated antibodies. **(b,c)** Scatter plot of average BRP/RBP-, BRP/Unc13A- and BRP/Syx-1A-intensity levels of AZs binned by their BRP intensity fit with regression lines for Ctrl (black) and PhTx (blue) treated (b) and in Wild-type (black) and *gluRIIA*$^{Null}$ mutant (light brown) (c) larvae. **(d)** Confocal scans of muscle 4 NMJs of segment A2-4 from 3$^{rd}$ instar Wild-type and *gluRIIA*$^{Null}$ larvae labelled with the indicated antibodies. In the second Wild-type column, the laser intensity for Unc18 was increased tenfold. **(e)** Quantification of Pearson's correlation coefficient for BRP and Unc18 in Wild-type (black) and *gluRIIA*$^{Null}$ (light brown) larvae. Exact values, detailed statistics including sample sizes and P values are listed in Supplementary Table 1. Scale bars: (a,d) 5 µm. Statistics: Mann-Whitney U test. ***P ≤ 0.001; n.s., not significant, P > 0.05. All panels show mean ± s.e.m..

### Supplementary Figure 2 – related to Figure 2

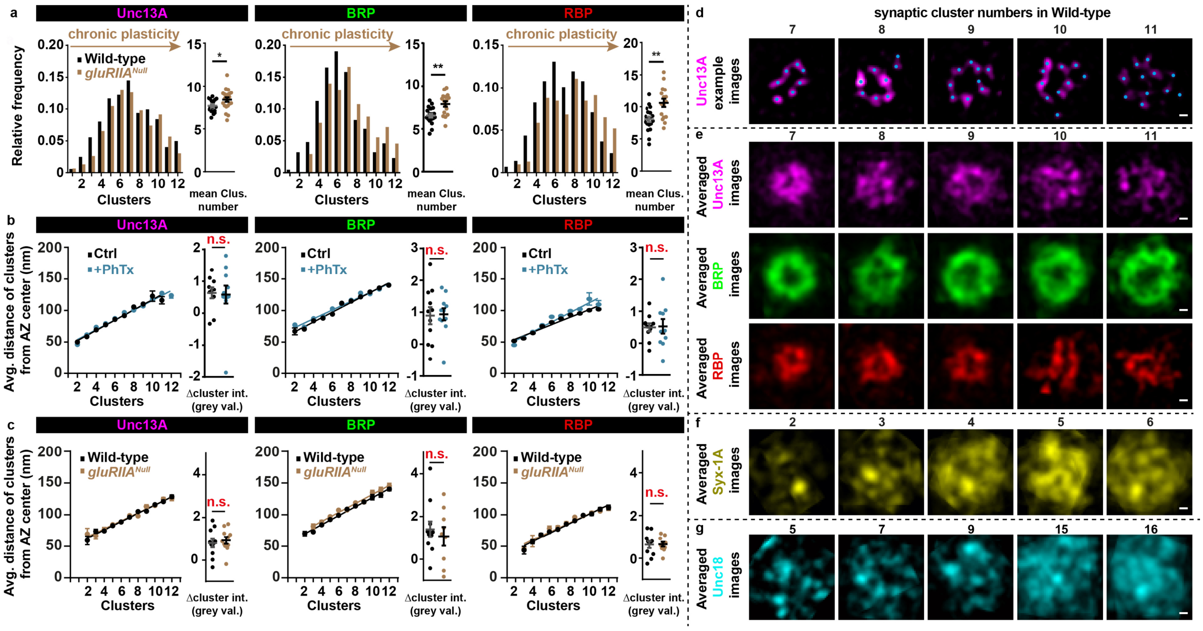


### Supplementary Figure 2: Analysis of AZ nano-architecture

**(a)** Frequency distribution of clusters (left) and average cluster number (right) of Unc13A, BRP and RBP in Wild-type (black) or *gluRIIA*$^{Null}$ (light brown) larvae. **(b,c)** Average AZ radius (left) and cluster intensity change (right) of Unc13A, BRP and RBP without (Ctrl; black) and with 10 minutes PhTx (+PhTx; blue) treatment (b) and in Wild-type (black) or *gluRIIA*$^{Null}$ (light brown) (c) larvae. **(d)** Example Unc13A STED-images of AZs with 7-11 clusters. **(e)** Average of rotated STED images stained against Unc13A (magenta), BRP (green) and RBP (red) with 7-11 clusters. **(f,g)** Average of rotated STED images stained against Syx-1A with 2-6 clusters (e) or Unc18 with 5, 7, 9, 15 and 16 clusters (f). Exact values, detailed statistics including sample sizes and P values are listed in Supplementary Table 1. Scale bars: (d-g) 50 nm. Statistics: Mann-Whitney U test. **P ≤ 0.01; n.s., not significant, P > 0.05. All panels show mean ± s.e.m.

### Supplementary Figure 3 – related to Figure 2

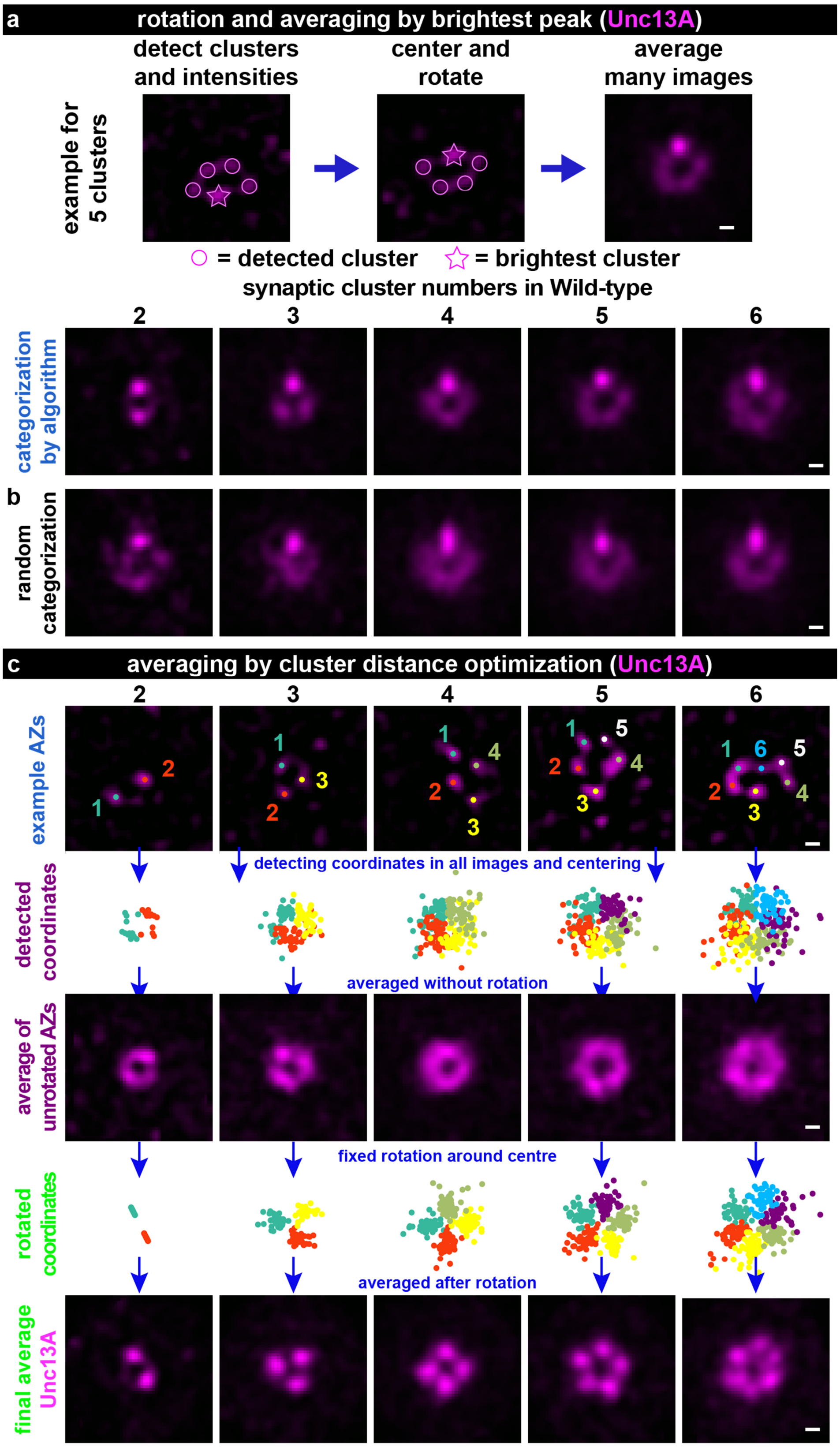


**Supplementary Figure 3: AZ protein averaging procedures exemplified for Unc13A.**

**(a,b)** Procedure 1: Single AZ STED images of Unc13A after categorization via peak finder algorithm (a) or after random categorization (b) are centered and only the highest intensity peak is considered for AZ rotation. Whole images are rotated to align the highest intensity peak on the vertical midline between the top quadrants. Images are shown for 2 to 6 clusters. **(c)** Averaging procedure of Unc13A-labelled single AZ images. The top row contains example AZ with 2-6 Unc13A clusters, marked by colored dots. In the second row, corresponding cluster positions from all AZ images containing the same number of clusters are shown together; in each AZ, clusters are counted in a counter-clockwise manner and all detected clusters from all images are shown by their rank in identical colors. Without further processing, averaging these images reveal a circular fluorescence pattern, because of the random localization of clusters in the different images (third row). Fixed rotation of cluster positions per image optimizes the overlap between the clusters of all images, and averaging of images then reveals a geometrical pattern (bottom row). Images are 510x510 nm. Scale bars: 50 nm.

**Supplementary Figure 4 – related to Figure 2**

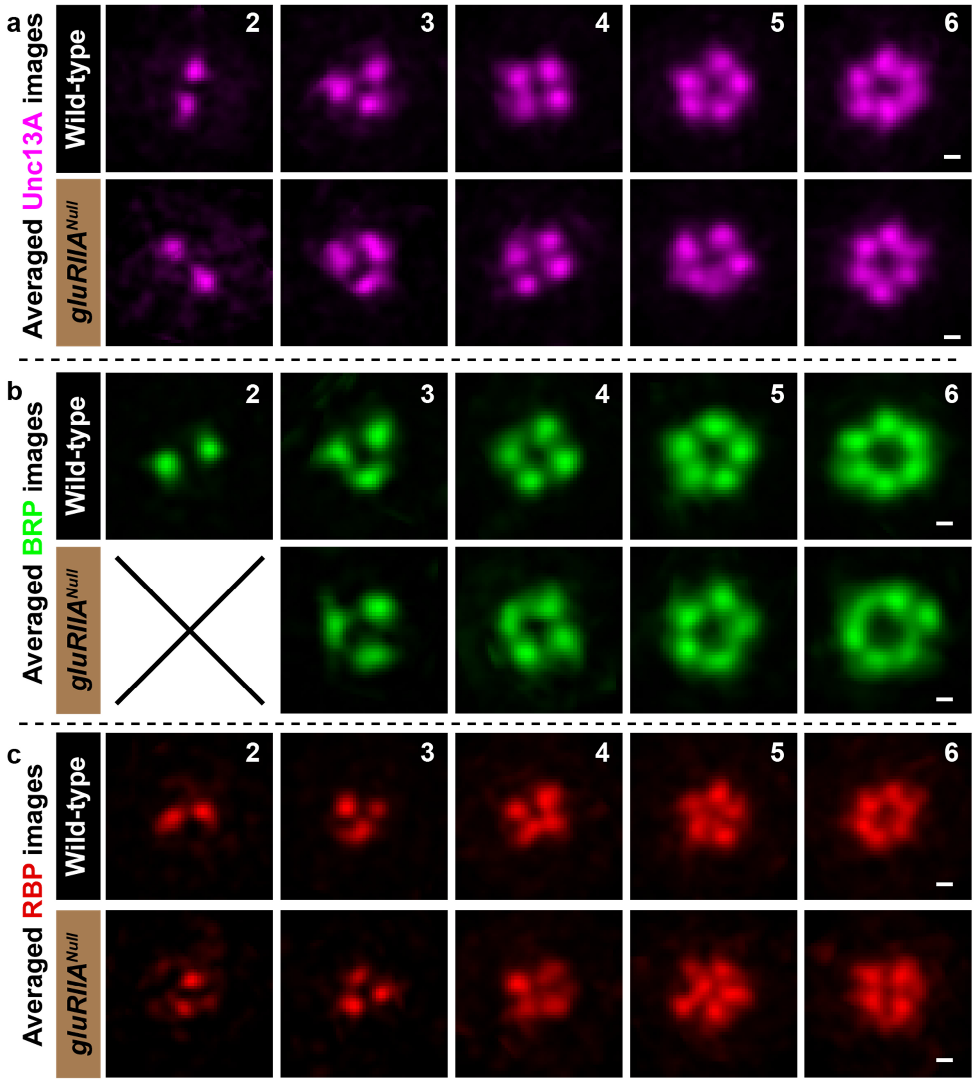


**Supplementary Figure 4: Chronic plasticity does not alter principle Unc13A/BRP/RBP single AZ protein architecture.**

**(a-c)** Averages of rotated STED images stained against Unc13A (a), BRP (b) and RBP (c) with 2-6 modules in Wild-type or *gluRIIA*$^{\text{Null}}$ larvae. Scale bars: 50 nm.

**Supplementary Figure 5 – related to Figure 3**

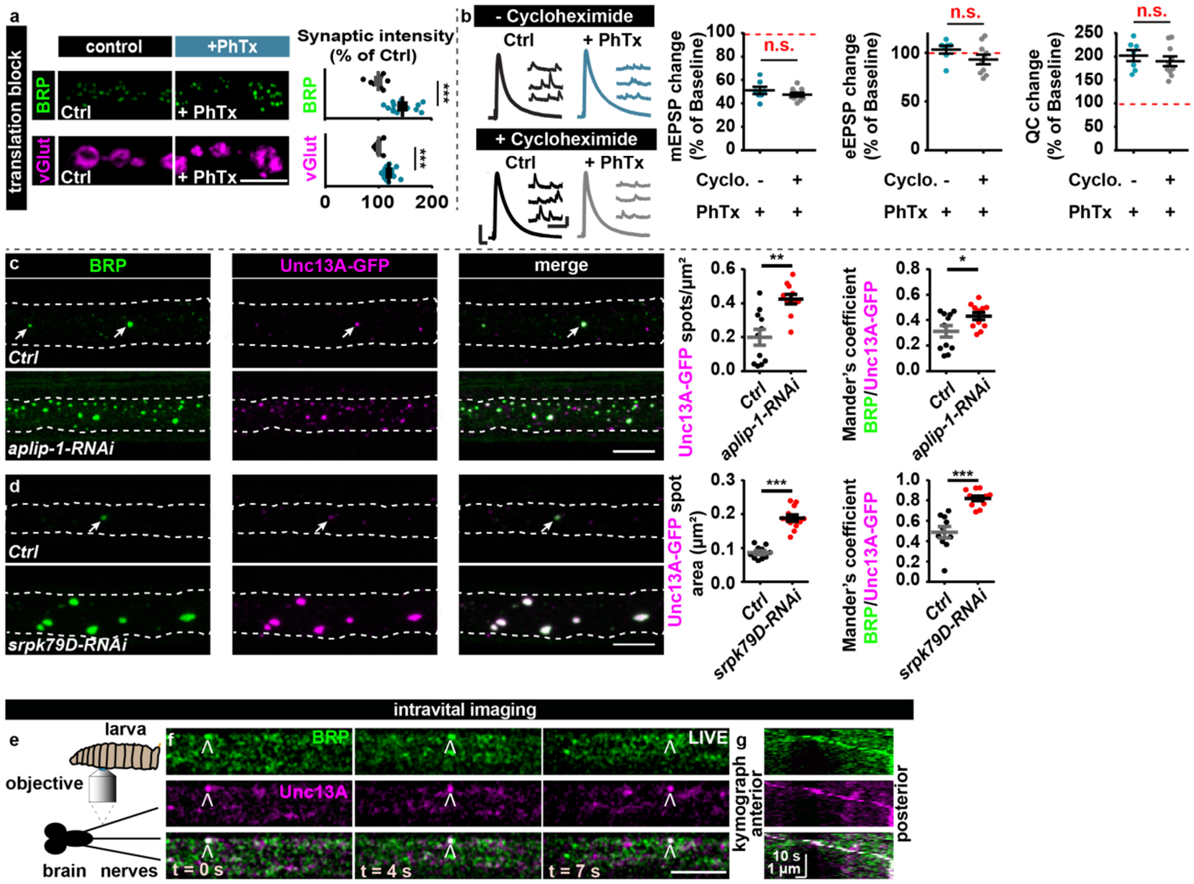


**Supplementary Figure 5: Translation block maintains PHP and AZ-remodelling, Unc13A co-accumulates in the axons of 3rd instar larvae upon *aplip-1*-KD and *srpk79d*-KD, BRP and Unc13A were seen to co-transport in motoneuronal axons.**

**(a)** Confocal images and quantification of AZ-fluorescence intensities at Wild-type NMJs labelled with the indicated antibodies after translation blockage with Cycloheximide without (Ctrl; black) and with 10 minutes PhTx (+PhTx; blue) treatment. Note that BRP AZ-levels were not changed after 10 minutes of Cycloheximide treatment in comparison to untreated NMJs (single BRP-AZ-levels without Cycloheximide: 1912 ± 109.3 a.u. vs. with Cycloheximide: 1943 ± 57.88 a.u.; P-value 0.7945, Student's t-test). **(b)** (left) Representative traces of eEPSP (evoked) and mEPSP (spont.) with or without the translation blocker Cycloheximide, and after control (Ctrl; black) or PhTx (+PhTx; blue) treatment for 10 minutes. (right) Quantifications of percentage change of mEPSP amplitude, eEPSP amplitude and quantal content (QC) in PhTx-treated Wild-type (blue) cells in the presence (+) or absence (-) of Cycloheximide compared to baseline of each control treatment (without PhTx, dashed red line corresponds to 100%/no change). **(c)** Nerve bundles of segments A1–A3 from third instar larvae of the respective genotypes labeled with the antibodies indicated. Arrows indicate axonal spots co-positive for BRP and Unc13A-GFP in the control situation. Quantification of axonal Unc13A-GFP spots per $\mu m^2$ and Mander's overlap coefficient for BRP and Unc13A-GFP. **(d)** Nerve bundles of segments A1–A3 from third instar larvae of the genotypes indicated labeled with the antibodies indicated. Arrows point to axonal spots co-positive for BRP and Unc13A-GFP in the control situation. Quantification of axonal Unc13A-GFP spot area in $\mu m^2$ and Mander's overlap coefficient for BRP and Unc13A-GFP. Dashed lines outline the axonal area. Exact values, detailed statistics including sample sizes and P values are listed in Supplementary Table 1. **(e)** Intravital imaging procedure. **(f,g)** Live imaging in intact third instar larvae (f) and dual-color kymograph (g) of axonal BRP (green) and Unc13A (magenta) showed anterograde co-transport of both proteins. See also **Movie 1**. Exact values, detailed statistics including sample sizes and P values are listed in Supplementary Table 1. Scale bars: (b) eEPSP: 50 ms, 4 mV ; mEPSP: 250 ms, 1 mV ; (a,c,d) 5 µm. Statistics: Mann-Whitney U test. $^*P \leq 0.05$; $^{**}P \leq 0.01$; $^{***}P \leq 0.001$. n.s., not significant, $P > 0.05$. All panels show mean ± s.e.m.

**Supplementary Figure 6 – related to Figure 3**

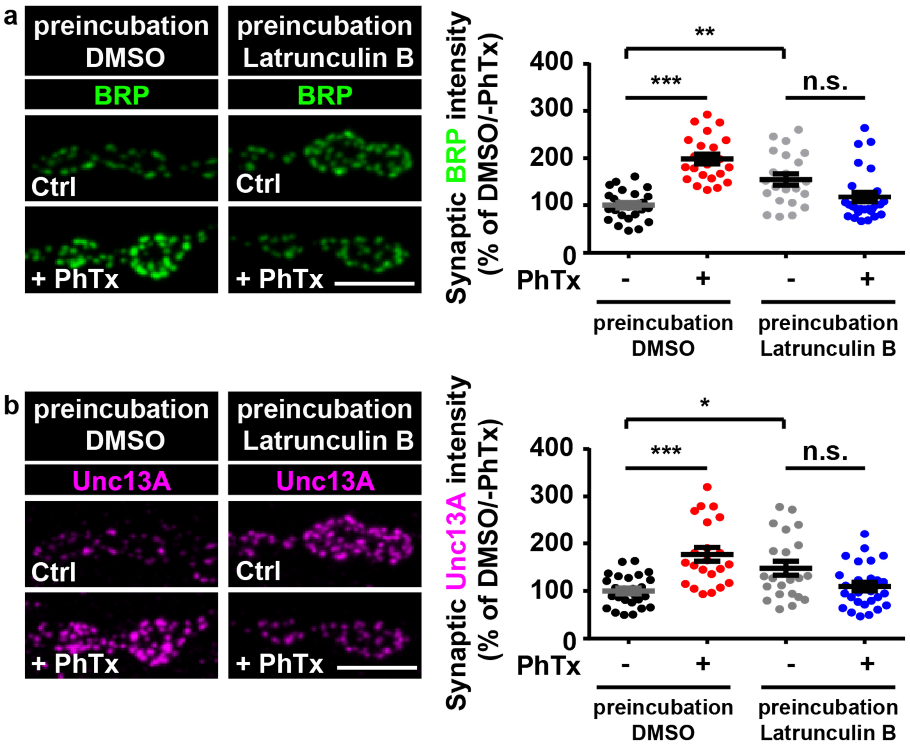


**Supplementary Figure 6: Actin polymerization is required for rapid AZ-remodeling.** **(a,b)** Confocal images and quantification of synaptic BRP or Unc13A intensities in % of DMSO/-PhTx without (preincubation DMSO) and with (preincubation Latrunculin B) labelled with the indicated antibodies without (Ctrl) and with 10 minutes PhTx (+PhTx) treatment. Exact values, detailed statistics including sample sizes and P values are listed in Supplementary Table 1. Scale bar: 5 µm. Statistics: nonparametric one-way analysis of variance (ANOVA) test, followed by a Tukey's multiple comparison test. $*P \leq 0.05$; $**P \leq 0.01$; n.s., not significant, $P > 0.05$. All panels show mean ± s.e.m..

**Supplementary Figure 7 – related to Figure 5**

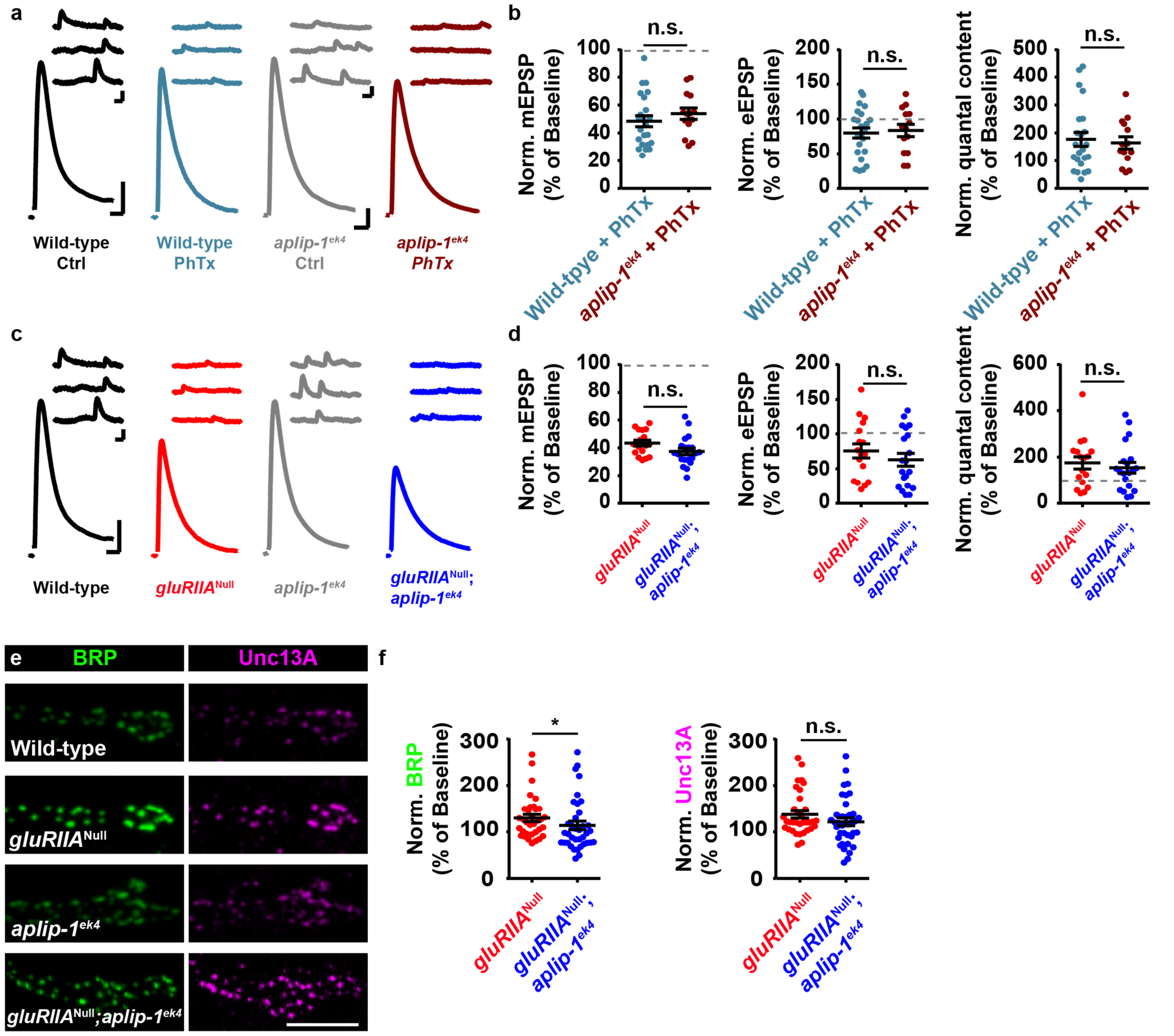


**Supplementary Figure 7: Loss of Aplip-1 in *gluRIIA*$^{\text{Null}}$ mutants moderately (i.e. non-significantly at 5% level) impairs the expression of chronic homeostatic plasticity.**

**(a)** Representative traces of eEPSP (evoked) and mEPSP (spont.) of the indicated genotypes with and without PhTx-treatment. **(b)** Quantifications of percentage change of mEPSP amplitude, eEPSP amplitude and quantal content (QC) in PhTx-treated genotypes compared to baseline (no PhTx) for each genotype (dashed red line corresponds to 100%/no change). **(c,d)** Same as in (a,b) but for Wild-type (black), *gluRIIA*$^{\text{Null}}$ (red), *aplip-1*$^{\text{ek4}}$ (grey) and *gluRIIA*$^{\text{Null}}$;*aplip-1*$^{\text{ek4}}$ (blue) but compared to baseline values of Wild-type for *gluRIIA*$^{\text{Null}}$ and *aplip-1*$^{\text{ek4}}$ for *gluRIIA*$^{\text{Null}}$;*aplip-1*$^{\text{ek4}}$ (dashed red line corresponds to 100%/no change). **(e)**

Confocal of muscle 4 NMJs of abdominal segment 2-5 from 3rd instar larvae at Wild-type, *gluRIIA*$^{\mathrm{Null}}$*, aplip-1*$^{ek4}$ and *gluRIIA*$^{\mathrm{Null}}$*,aplip-1*$^{ek4}$ NMJs labelled with the indicated antibodies. **(f)** Quantification of percentage change of synaptic BRP and Unc13A levels in *gluRIIA*$^{\mathrm{Null}}$ (red) and *gluRIIA*$^{\mathrm{Null}}$;*aplip-1*$^{\mathrm{ek4}}$ (blue) mutants compared to baseline fluorescence values of Wild-type for *gluRIIA*$^{\mathrm{Null}}$ and *aplip-1*$^{\mathrm{ek4}}$ for *gluRIIA*$^{\mathrm{Null}}$;*aplip-1*$^{\mathrm{ek4}}$ (dashed grey line corresponds to 100%/no change). Exact normalized and raw values, detailed statistics including sample sizes and P values are listed in Supplementary table 1. See also Supplementary figure 10 and 11 for non-normalized values. Scale bars: (a,c) eEPSP: 25 ms, 5 mV; mEPSP: 50 ms, 1 mV; (e) 5 µm. Statistics: Student's T-test for all comparisons except (d) quantal content and (f) where a Mann-Whitney U Test was performed. * P ≤ 0.05; **P ≤ 0.01; ***P ≤ 0.001; n.s., not significant, P > 0.05. All panels show mean ± s.e.m..

**Supplementary Figure 8 – related to Figure 6**

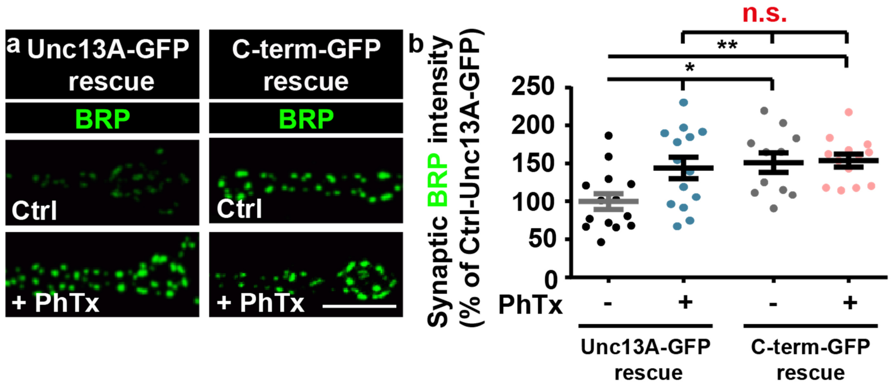


**Supplementary Figure 8: Enhanced synaptic BRP levels and loss of AZ-remodeling at *unc13*$^{Null}$ NMJs expressing the C-term GFP construct.**

**(a,b)** Confocal images and quantification of synaptic BRP intensities in % of Unc13A-GFP rescue at NMJs pan-neuronally re-expressing Unc13A-GFP or C-term-GFP in the *unc13*$^{Null}$ background labelled with the indicated antibodies without (Ctrl; black (Unc13A-GFP rescue); grey (C-term-GFP rescue)) and with 10 minutes PhTx (+PhTx; blue (Unc13A-GFP rescue); light red (C-term-GFP rescue)) treatment. Exact values, detailed statistics including sample sizes and P values are listed in Supplementary Table 1. Scale bar: 5 µm. Statistics: nonparametric one-way analysis of variance (ANOVA) test, followed by a Tukey's multiple comparison test. *$P \leq 0.05$; **$P \leq 0.01$; n.s., not significant, $P > 0.05$. All panels show mean ± s.e.m..

**Supplementary Figure 9 – related to Figure 8**

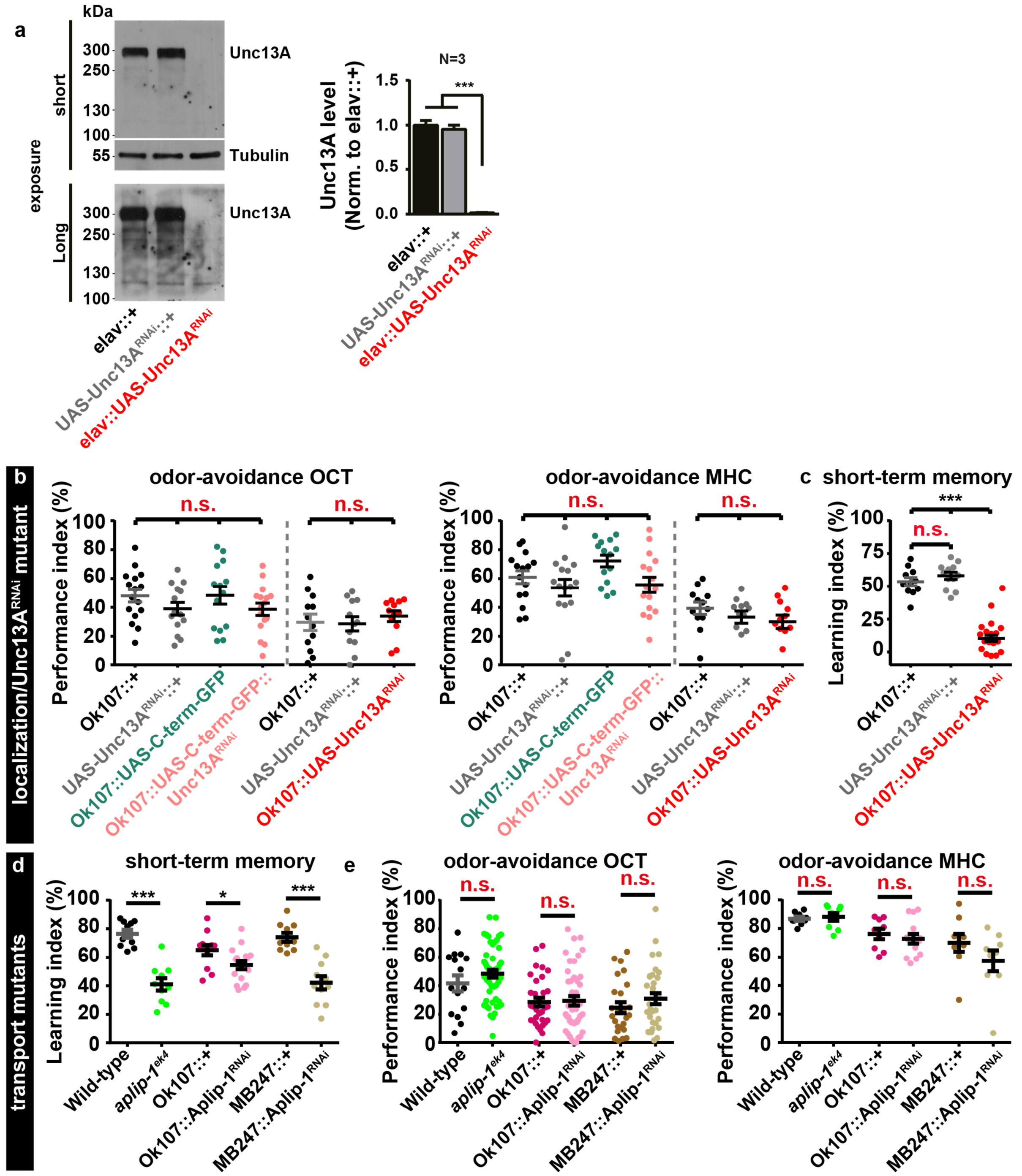


**Supplementary Figure 9: Efficiency of Unc13A RNAi, odor avoidance controls for behavioral analyses and short-term memory impairment upon Unc13A KD and in Aplip-1 mutants.**

**(a)** Whole brain western blot and quantification of Unc13A-levels normalized to driver-control of pan-neuronally expressed Unc13A$^{RNAi}$ and control flies with short (top) and longer exposure (bottom). **(b)** Quantification of odor-avoidance performance index of OCT (left) or MHC (right) in the indicated genotypes. **(c)** Short-term memory scores after mushroom body-specific Unc13A downregulation via locally driven RNAi expression (Ok107:: Unc13ARNAi) compared to controls expressing the driver, but not the RNAi (Ok107::+) or the RNAi without driver (UAS-Unc13ARNAi::+). **(d)** Short-term memory scores in *aplip-1*$^{ek4}$ mutant flies compared to Wild-type and ones after mushroom body-specific Aplip-1 downregulation via locally driven RNAi expression (Ok107::Aplip-1$^{RNAi}$) or (MB247::Aplip-1$^{RNAi}$) compared to controls expressing the driver, but not the RNAi (Ok107::+ or MB247::+). **(e)** Quantification of odor-avoidance performance index of OCT (left) or MHC (right) in the indicated genotypes. Exact values, detailed statistics including sample sizes and P values are listed in Supplementary Table 1. Statistics: (a-c) nonparametric one-way analysis of variance (ANOVA) test, followed by a Tukey's multiple comparison test. (d,e) Mann-Whitney U test. *P ≤ 0.05; ***P ≤ 0.001; n.s., not significant, P > 0.05. All panels show mean ± s.e.m..

### Supplementary Figure 10 – related to Figure 5, 6 and Supplementary Fig. 7

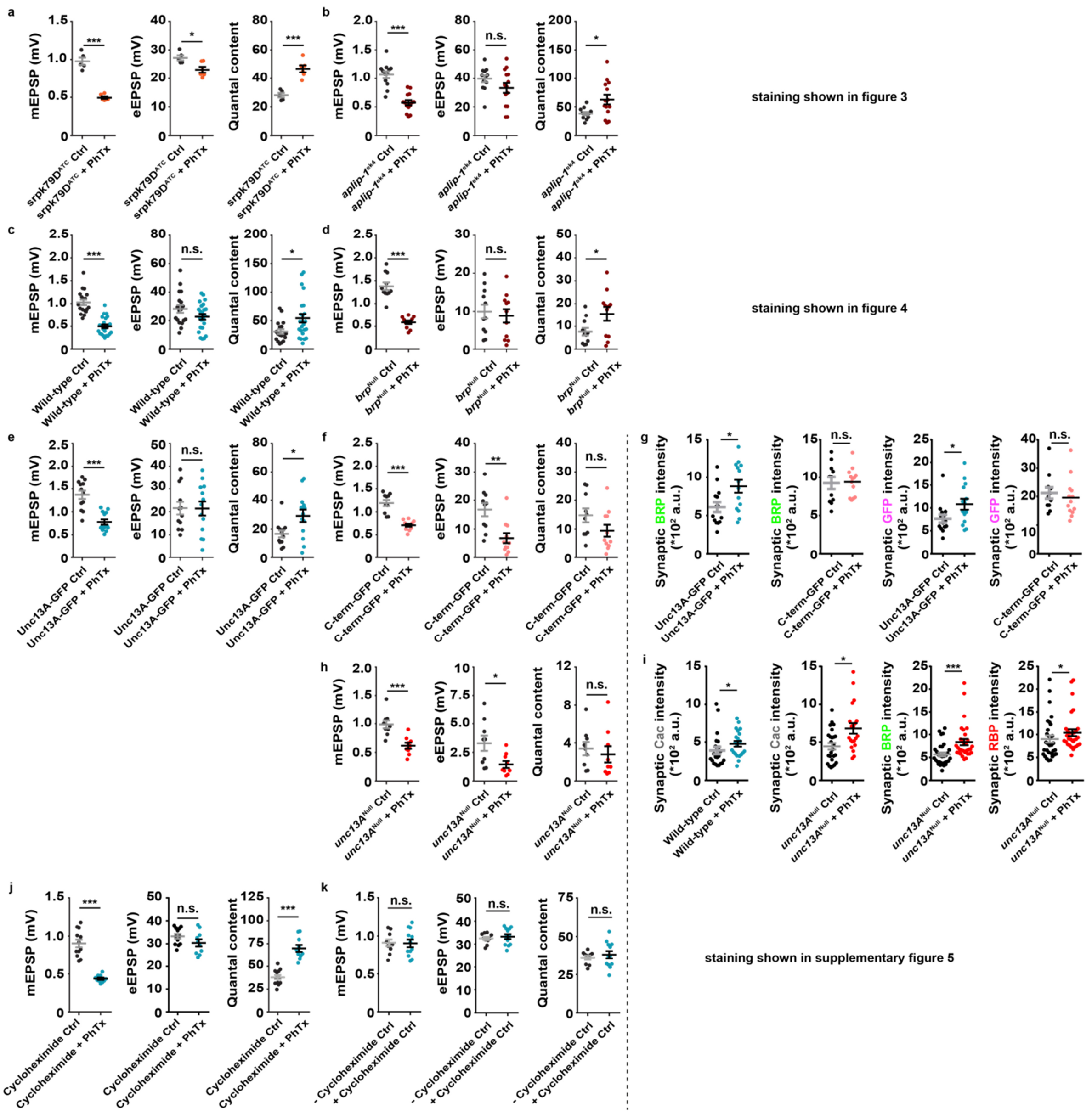


**Supplementary Figure 10: Non-normalized values of electrophysiological and single AZ imaging experiments**

(a-f,h,j) Non-normalized values of mEPSP amplitude, eEPSP amplitude and quantal content without (Ctrl) and with (+PhTx) Phtx-treatment in the genotypes indicated. (g,i) Quantification of BRP, GFP, Cac or RBP AZ-levels in Ctrl and PhTx (+PhTx) treated animals of the indicated genotypes. (k) Non-normalized values of mEPSP amplitude, eEPSP amplitude and quantal content of cells without (-Cycloheximide) and with (+Cycloheximide)-

treatment. Statistics: Student's t test for all comparisons except for (e) quantal content, (f) eEPSP amplitude, (g), (h) quantal content and (i) where a Mann-Whitney U test was performed. * $P \leq 0.05$; **$P \leq 0.01$; ***$P \leq 0.001$; n.s., not significant, $P > 0.05$. All panels show mean ± s.e.m..

**Supplementary Figure 11 – related to Figure 5 and Supplementary Fig. 7**

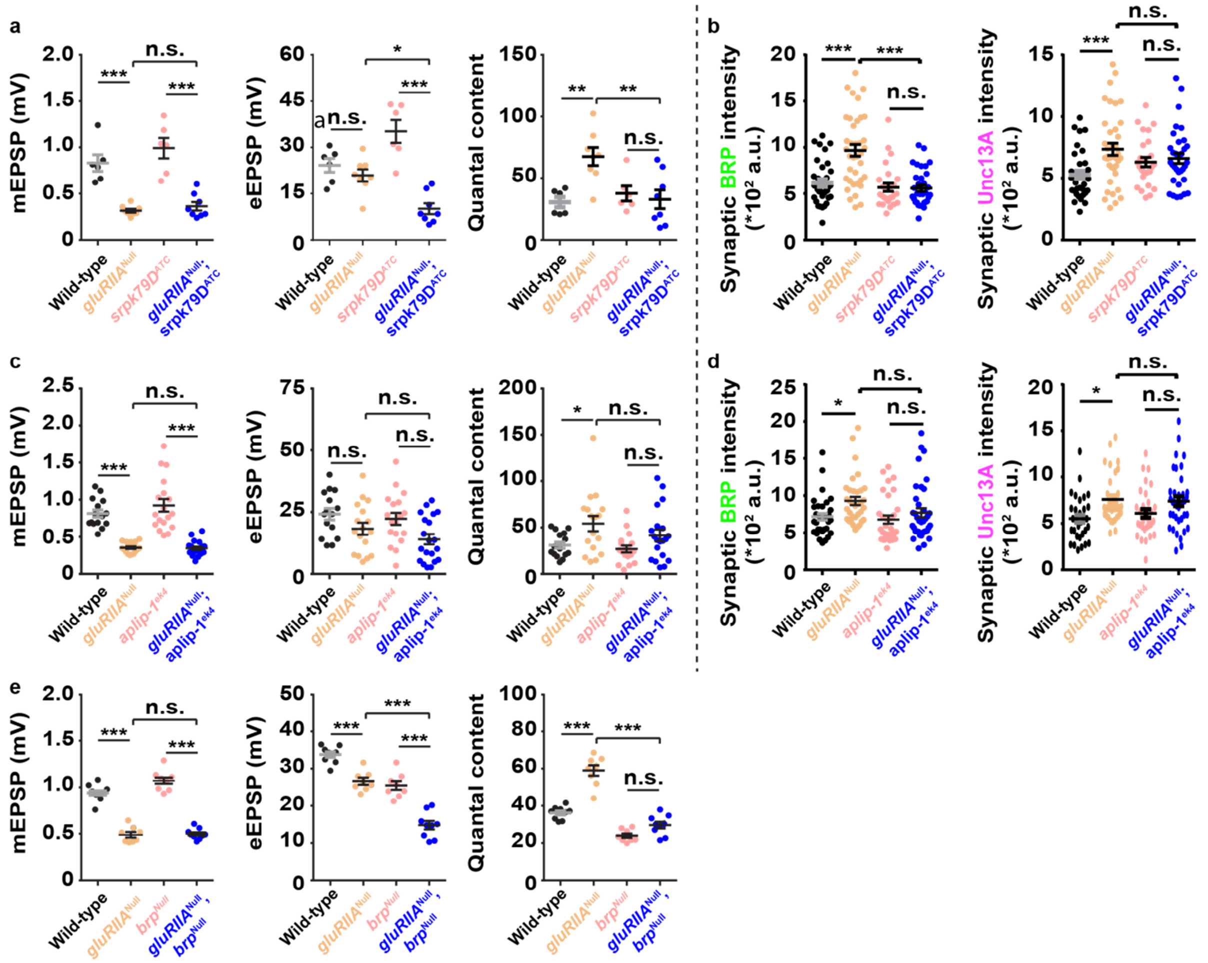


**Supplementary Figure 11: Non-normalized values of electrophysiological and single AZ imaging experiments**

(a,c,e) Non-normalized values of mEPSP amplitude, eEPSP amplitude and quantal content in the genotypes indicated. (b,d) Quantification BRP and Unc13A AZ-levels in the genotypes indicated. Statistics: nonparametric one-way analysis of variance (ANOVA) test, followed by a Tukey's multiple comparison test. * $P \leq 0.05$; **$P \leq 0.01$; ***$P \leq 0.001$; n.s., not significant, $P > 0.05$. All panels show mean ± s.e.m..

### Additional titles and legends

**Movie 1: Axonal Co-transport of BRP and Unc13A.**

Live imaging in intact third instar larvae of axonal (segment A2) BRP (green) and Unc13A (magenta) showed anterograde co-transport of both proteins.

**Supplementary Table 1 – Figure source data, number of experiments and P values – related to all figures including supplementary Figures**

Supplementary Table 1 provides all figure source data (including raw and normalized values), number of experiments or data points, P values, tests of normality and the nature of the statistical comparisons used in this study.
