## Supplementary material for "Rapid active zone remodeling consolidates presynaptic potentiation"

Supplementary table 1 - related to all figures of the manuscript

**Summary of all obtained parameters in this study**

**Current clamp recordings:**

| | **mean ± SEM** | | | | | | | |
|---|---|---|---|---|---|---|---|---|
| **Parameter** | **control treatment (n)** | **PhTx treatment (n)** | **Mann-Whitney U-test (P values)** | **test statistic** | **Unpaired t test (P values)** | **test statistic** | **D'Agostino & Pearson omnibus normality test** | |
| **(Figure)** | | | **two-tailed** | | **two-tailed** | | **control treatment** | **PhTx treatment** |
| **Representative PhTx traces and quantification** | | | | | | | **P values and passed normality test (α=0.05)?** | |
| **Induction of Rapid Homeostatic Plasticity** | | | | | | | *** indicates normality test failed** | |
| **(Figure 1b and 6d)** | | | | | | | | |
| **Wild-type** | 8 cells from 6 animals | 8 cells from 5 animals | | | | | | |
| mEPSP amplitude (mV) | 0.8074 ± 0.03153 (8) | 0.4539 ± 0.03266 (8) | 0.0002*** | U = 0 | < 0.0001*** | t=7.786 df=14 | 0.1272 | 0.506 |
| eEPSP amplitude (mV) | 17.32 ± 2.681 (8) | 15.57 ± 1.609 (8) | 0.7209 | U = 28 | 0.5844 | t=0.5599 df=14 | 0.2912 | 0.6423 |
| quantal content | 21.15 ± 2.758 (8) | 34.40 ± 2.616 (8) | 0.0047** | U = 6 | 0.0036** | t=3.486 df=14 | 0.5648 | 0.4318 |
| **(Figure 6d)** | | | | | | | | |
| ***unc13A*** $^{Null}$ | 9 cells from 5 animals | 9 cells from 5 animals | | | | | | |
| mEPSP amplitude (mV) | 0.9881 ± 0.06958 (9) | 0.6169 ± 0.05206, (9) | 0.0005*** | U = 4 | 0.0006*** | t=4.272 df=16 | 0.0873 | 0.835 |
| eEPSP amplitude (mV) | 3.296 ± 0.6671 (9) | 1.453 ± 0.3050 (9) | 0.0188* | U = 14 | 0.023* | t=2.514 df=16 | 0.6709 | 0.4974 |
| quantal content | 3.395 ± 0.6906, (9) | 2.795 ± 0.8457, (9) | 0.3865 | U = 30 | 0.5901 | t=0.5497 df=16 | 0.3651 | 0.0433* |
| **(Normalized)** | | | | | | | | |
| **Wild-type** | 8 cells from 6 animals | 8 cells from 5 animals | | | | | | |
| mEPSP amplitude (normalized) | 100.0 ± 3.906, (8) | 56.22 ± 4.045, (8) | 0.0002*** | U = 0 | < 0.0001*** | t=7.786 df=14 | 0.1272 | 0.506 |
| eEPSP amplitude (normalized) | 100.0 ± 15.48, (8) | 89.89 ± 9.288, (8) | 0.7209 | U = 28 | 0.5844 | t=0.5599 df=14 | 0.2912 | 0.6423 |
| quantal content (normalized) | 100.0 ± 13.04, (8) | 162.7 ± 12.37, (8) | 0.0047** | U = 6 | 0.0036** | t=3.486 df=14 | 0.5648 | 0.4318 |
| ***unc13A*** $^{Null}$ | 9 cells from 5 animals | 9 cells from 5 animals | | | | | | |
| mEPSP amplitude (normalized) | 100.0 ± 7.042, (9) | 62.43 ± 5.269, (9) | 0.0005*** | U = 4 | 0.0006* | t=4.272 df=16 | 0.0873 | 0.835 |
| eEPSP amplitude (normalized) | 100.0 ± 20.24, (9) | 44.07 ± 9.253, (9) | 0.0188* | U = 14 | 0.023* | t=2.514 df=16 | 0.6709 | 0.4974 |
| quantal content (normalized) | 100.0 ± 20.34, (9) | 82.32 ± 24.91, (9) | 0.3865 | U = 30 | 0.5901 | t=0.5497 df=16 | 0.3651 | 0.0433* |
| **Normalized comparison of PhTx treatment** | | | **two-tailed** | | **two-tailed** | | **D'Agostino & Pearson omnibus normality test** | |
| **Figure 6d** | **Wild-type +PhTx (n)** | ***unc13A*** $^{Null}$ **+ PhTx (n)** | **Wild-type +PhTx vs. *unc13A*** $^{Null}$ **PhTx** | | **Wild-type +PhTx vs. *unc13A*** $^{Null+}$**PhTx** | | **Wild-type +PhTx** | ***unc13A*** $^{Null}$ **+PhTx** |
| mEPSP amplitude (normalized) | 56.22 ± 4.045, (8) | 62.43 ± 5.269, (9) | 0.4807 | U = 28 | 0.3736 | t=0.9171 df=15 | 0.506 | 0.835 |
| eEPSP amplitude (normalized) | 89.89 ± 9.288, (8) | 44.07 ± 9.253, (9) | 0.0079** | U = 9 | 0.0033** | t=3.483 df=15 | 0.6423 | 0.4974 |
| quantal content (normalized) | 162.7 ± 12.37, (8) | 82.32 ± 24.91, (9) | 0.0152* | U = 11 | 0.0141* | t=2.775 df=15 | 0.4318 | 0.0433 |
| **Parameter** | **control treatment (n)** | **PhTx treatment (n)** | **Mann-Whitney U-test (P values)** | **test statistic** | **Unpaired t test (P values)** | **test statistic** | **D'Agostino & Pearson omnibus normality test** | |
| **(Figure 6i)** | | | **two-tailed** | | **two-tailed** | | **control treatment** | **PhTx treatment** |
| **Unc13A-GFP rescue** | 12 cells from 7 animals | 13 cells from 8 animals | | | | | | |
| mEPSP amplitude (mV) | 1.372 ± 0.09085 (12) | 0.7788 ± 0.05711, (13) | < 0.0001*** | U = 9.5 | < 0.0001*** | t=5.617 df=23 | 0.4425 | 0.1114 |
| eEPSP amplitude (mV) | 21.59 ± 2.756 (12) | 21.44 ± 3.101 (13) | 0.9787 | U = 77 | 0.9714 | t=0.03624 df=23 | 0.4724 | 0.4572 |
| quantal content | 16.63 ± 2.498 (12) | 29.20 ± 4.383 (13) | 0.0398* | U = 40 | 0.023* | t=2.436 df=23 | 0.0168* | 0.9666 |
| **C-term-GFP rescue** | 10 cells from 6 animals | 11 cells from 7 animals | | | | | | |
| mEPSP amplitude (mV) | 1.194 ± 0.06731 (10) | 0.7063 ± 0.03215 (11) | < 0.0001*** | U = 0 | < 0.0001*** | t=6.731 df=19 | 0.285 | 0.8552 |
| eEPSP amplitude (mV) | 16.77 ± 2.326 (10) | 6.800 ± 1.744 (11) | 0.0028** | U = 14 | 0.0026** | t=3.469 df=19 | 0.8914 | 0.0147* |
| quantal content | 14.77 ± 2.405 (10) | 9.416 ± 2.115 (11) | 0.0845 | U = 30 | 0.1096 | t=1.678 df=19 | 0.4915 | 0.2951 |
| **(Normalized)** | | | | | | | | |
| **Unc13A-GFP rescue** | 12 cells from 7 animals | 13 cells from 8 animals | | | | | | |
| mEPSP amplitude (normalized) | 100.0 ± 6.624, (12) | 56.78 ± 4.163 (13) | < 0.0001*** | U = 9.5 | < 0.0001*** | t=5.617 df=23 | 0.4425 | 0.1114 |
| eEPSP amplitude (normalized) | 100.0 ± 12.77, (12) | 99.30 ± 14.37 (13) | 0.9787 | U = 77 | 0.9714 | t=0.03624 df=23 | 0.4724 | 0.4572 |
| quantal content (normalized) | 100.0 ± 15.02, (12) | 175.6 ± 26.35 (13) | 0.0398* | U = 40 | 0.023* | t=2.436 df=23 | 0.0168* | 0.9666 |
| **C-term-GFP rescue** | 10 cells from 6 animals | 11 cells from 7 animals | | | | | | |
| mEPSP amplitude (normalized) | 100.0 ± 5.639 (10) | 59.18 ± 2.694 (11) | < 0.0001*** | U = 0 | < 0.0001*** | t=6.731 df=19 | 0.285 | 0.8552 |
| eEPSP amplitude (normalized) | 100.0 ± 13.87 (10) | 40.55 ± 10.40 (11) | 0.0028** | U = 14 | 0.0026** | t=3.469 df=19 | 0.8914 | 0.0147* |
| quantal content (normalized) | 100.0 ± 16.28 (10) | 63.75 ± 14.32 (11) | 0.0845 | U = 30 | 0.1096 | t=1.678 df=19 | 0.4915 | 0.2951 |
| **Normalized comparison of PhTx treatment** | | | **Mann-Whitney U-test (P values)** | **test statistic** | **Unpaired t test (P values)** | **test statistic** | | |
| **Figure 6i** | | | **two-tailed** | | **two-tailed** | | **D'Agostino & Pearson omnibus normality test** | |
| | **Unc13A-GFP rescue +PhTx (n)** | **C-term-GFP rescue +PhTx (n)** | **Unc13A-GFP rescue +PhTx vs. C-term-GFP rescue +PhTx** | | **Unc13A-GFP rescue +PhTx vs. C-term-GFP rescue +PhTx** | | **Unc13A-GFP rescue PhTx** | **C-term-GFP rescue PhTx** |
| mEPSP amplitude (normalized) | 56.78 ± 4.163 (13) | 59.18 ± 2.694 (11) | 0.494 | U = 59 | 0.6475 | t=0.4636 df=22 | 0.1114 | 0.8552 |
| eEPSP amplitude (normalized) | 99.30 ± 14.37 (13) | 40.55 ± 10.40 (11) | 0.0048** | U = 24 | 0.0041** | t=3.203 df=22 | 0.4572 | 0.0147* |
| quantal content (normalized) | 175.6 ± 26.35 (13) | 63.75 ± 14.32 (11) | 0.0031** | U =22 | 0.0018** | t=3.538 df=22 | 0.9666 | 0.2951 |

| Parameter (Figure) | control treatment (n) | PhTx treatment (n) | Mann-Whitney U-test (P values) two-tailed | test statistic | Unpaired t test (P values) two-tailed | test statistic | D'Agostino & Pearson omnibus normality test control treatment | PhTx treatment |
|---|---|---|---|---|---|---|---|---|
| **(Figure 5b)** | | | | | | | | |
| **Wild-type** | 18 cells from 10 animals | 23 cells from 12 animals | | | | | **control treatment** | **PhTx treatment** |
| mEPSP amplitude (mV) | 1.031 ± 0.06045 (18) | 0.4999 ± 0.04065 (23) | < 0.0001*** | U = 17 | < 0.0001*** | t=7.537 df=39 | 0.1633 | 0.296 |
| eEPSP amplitude (mV) | 28.41 ± 2.743 (18) | 22.77 ± 2.075 (23) | 0.2146 | U = 159 | 0.1024 | t=1.673 df=39 | 0.3592 | 0.3596 |
| quantal content | 30.66 ± 4.253 (18) | 54.34 ± 7.698 (23) | 0.0219* | U = 120 | 0.017* | t=2.494 df=39 | 0.0884 | 0.1113 |
| ***brp* Null** | 11 cells from 6 animals | 11 cells from 6 animals | | | | | | |
| mEPSP amplitude (mV) | 1.381 ± 0.08217 (11) | 0.5881 ± 0.03854 (11) | < 0.0001*** | U = 0 | < 0.0001*** | t=8.738 df=20 | 0.7995 | 0.5946 |
| eEPSP amplitude (mV) | 9.931 ± 1.850 (11) | 8.793 ± 1.804 (11) | 0.5619 | U = 51 | 0.6643 | t=0.4405 df=20 | 0.3924 | 0.6004 |
| quantal content | 7.632 ± 1.679 (11) | 15.54 ± 3.108 (11) | 0.1014 | U = 35 | 0.0368* | t=2.237 df=20 | 0.364 | 0.8273 |
| **(Supplementary Figure 7b)** | | | | | | | | |
| ***aplip-1* ek4** | 12 cells from 6 animals | 14 cells from 7 animals | | | | | | |
| mEPSP amplitude (mV) | 1.065 ± 0.06161 (12) | 0.5736 ± 0.04377 (14) | < 0.0001*** | U = 6 | < 0.0001*** | t=6.635 df=24 | 0.7467 | 0.8734 |
| eEPSP amplitude (mV) | 39.69 ± 2.450 (12) | 33.23 ± 3.482 (14) | 0.3217 | U = 64 | 0.155 | t=1.468 df=24 | 0.1089 | 0.6028 |
| quantal content | 38.19 ± 2.803 (12) | 62.76 ± 8.425 (14) | 0.0234 | U = 40 | 0.0161 | t=2.589 df=24 | 0.6746 | 0.6049 |
| **(Normalized) (Figure 5b)** | | | | | | | | |
| **Wild-type** | 18 cells from 10 animals | 23 cells from 12 animals | | | | | | |
| mEPSP amplitude (normalized) | 100.0 ± 5.864 (18) | 48.50 ± 3.944 (23) | < 0.0001*** | U = 17 | < 0.0001*** | t=7.537 df=39 | 0.1633 | 0.296 |
| eEPSP amplitude (normalized) | 100.0 ± 9.655 (18) | 80.14 ± 7.303 (23) | 0.2146 | U = 159 | 0.1024 | t=1.673 df=39 | 0.3592 | 0.3596 |
| quantal content (normalized) | 100.0 ± 13.87 (18) | 177.3 ± 25.11 (23) | 0.0219* | U = 120 | 0.017* | t=2.494 df=39 | 0.0884 | 0.1113 |
| ***brp* Null** | 11 cells from 6 animals | 11 cells from 6 animals | | | | | | |
| mEPSP amplitude (normalized) | 100.0 ± 5.949 (11) | 42.58 ± 2.790 (11) | < 0.0001*** | U = 0 | < 0.0001*** | t=8.738 df=20 | 0.7995 | 0.5946 |
| eEPSP amplitude (normalized) | 100.0 ± 18.63 (11) | 88.54 ± 18.16 (11) | 0.5619 | U = 51 | 0.6643 | t=0.4405 df=20 | 0.3924 | 0.6004 |
| quantal content (normalized) | 100.0 ± 22.00 (11) | 203.5 ± 40.72 (11) | 0.1014 | U = 35 | 0.0368* | t=2.237 df=20 | 0.364 | 0.8273 |
| **(Supplementary Figure 7b)** | | | | | | | | |
| ***aplip-1* ek4** | 12 cells from 6 animals | 14 cells from 7 animals | | | | | | |
| mEPSP amplitude (normalized) | 100.0 ± 5.786 (12) | 53.87 ± 4.111 (14) | < 0.0001*** | U = 6 | < 0.0001*** | t=6.635 df=24 | 0.7467 | 0.8734 |
| eEPSP amplitude (normalized) | 100.0 ± 6.173 (12) | 83.73 ± 8.773 (14) | 0.3217 | U = 64 | 0.155 | t=1.468 df=24 | 0.1089 | 0.6028 |
| quantal content (normalized) | 100.0 ± 7.340 (12) | 164.3 ± 22.06 (14) | 0.0234 | U = 40 | 0.0161* | t=2.589 df=24 | 0.6746 | 0.6049 |

| Normalized comparison of PhTx treatment Figure 5b | | | Mann-Whitney U-test (P values) two-tailed | test statistic | Unpaired t test (P values) two-tailed | test statistic | D'Agostino & Pearson omnibus normality test | |
|---|---|---|---|---|---|---|---|---|
| | **Wild-type +PhTx** | ***brp* Null +PhTx** | **Wild-type +PhTx vs. *brp* Null +PhTx** | | **Wild-type +PhTx vs. *brp* Null +PhTx** | | **Wild-type +PhTx** | ***brp* Null +PhTx** |
| mEPSP amplitude (normalized) | 48.50 ± 3.944 (23) | 42.58 ± 2.790 (11) | 0.6312 | U = 113 | 0.3356 | t=0.9775 df=32 | 0.296 | 0.5946 |
| eEPSP amplitude (normalized) | 80.14 ± 7.303 (23) | 88.54 ± 18.16 (11) | 0.717 | U = 116 | 0.6099 | t=0.5152 df=32 | 0.3596 | 0.6004 |
| quantal content (normalized) | 177.3 ± 25.11 (23) | 203.5 ± 40.72 (11) | 0.5372 | U = 109 | 0.5707 | t=0.5729 df=32 | 0.1113 | 0.8273 |
| **Suplementary Fig. 7b** | **Wild-type +PhTx** | ***aplip-1* ek4 + PhTx** | **Wild-type +PhTx vs. *aplip-1* ek4 +PhTx** | | **Wild-type +PhTx vs. *aplip-1* ek4 +PhTx** | | **Wild-type +PhTx** | ***aplip-1* ek4 +PhTx** |
| mEPSP amplitude (normalized) | 48.50 ± 3.944 (23) | 53.87 ± 4.111 (14) | 0.1282 | U = 112 | 0.2453 | t=1.182 df=35 | 0.296 | 0.8734 |
| eEPSP amplitude (normalized) | 80.14 ± 7.303 (23) | 83.73 ± 8.773 (14) | 0.7928 | U = 152 | 0.7589 | t=0.3094 df=35 | 0.3596 | 0.6028 |
| quantal content (normalized) | 177.3 ± 25.11 (23) | 164.3 ± 22.06 (14) | 0.8167 | U = 153 | 0.7257 | t=0.3537 df=35 | 0.1113 | 0.6049 |

| Parameter (Figure) | control treatment (n) | PhTx treatment (n) | Mann-Whitney U-test (P values) two-tailed | test statistic | Unpaired t test (P values) two-tailed | test statistic | D'Agostino & Pearson omnibus normality test control treatment | PhTx treatment |
|---|---|---|---|---|---|---|---|---|
| **(Figure 5c)** | | | | | | | | |
| **Wild-type** | 6 cells from 3 animals | 6 cells from 3 animals | | | | | | |
| mEPSP amplitude (mV) | 0.9678 ± 0.04384 (6) | 0.4950 ± 0.02273 (6) | 0.0022** | U = 0 | < 0.0001*** | t=9.574 df=10 | Not determined | Not determined |
| eEPSP amplitude (mV) | 27.84 ± 1.666 (6) | 22.03 ± 1.582 (6) | 0.0411* | U = 5 | 0.0301* | t=2.527 df=10 | Not determined | Not determined |
| quantal content | 28.74 ± 1.054 (6) | 44.79 ± 3.143 (6) | 0.0022** | U = 0 | 0.0007*** | t=4.842 df=10 | Not determined | Not determined |
| ***srpk79D* ATC** | 6 cells from 3 animals | 6 cells from 3 animals | | | | | | |
| mEPSP amplitude (mV) | 0.9774 ± 0.04854 (5) | 0.4953 ± 0.01627 (6) | 0.0043** | U = 0 | < 0.0001*** | t=10.18 df=9 | Not determined | Not determined |
| eEPSP amplitude (mV) | 27.26 ± 0.8913 (5) | 23.06 ± 1.050 (6) | 0.0519 | U = 4 | 0.0156* | t=2.974 df=9 | Not determined | Not determined |
| quantal content | 28.15 ± 1.585 (5) | 46.77 ± 2.493 (6) | 0.0043** | U = 0 | 0.0002*** | t=5.997 df=9 | Not determined | Not determined |
| **(Normalized)** | | | | | | | | |
| **Wild-type** | 6 cells from 3 animals | 6 cells from 3 animals | | | | | | |
| mEPSP amplitude (normalized) | 100.0 ± 4.530 (6) | 51.15 ± 2.349 (6) | 0.0022** | U = 0 | < 0.0001*** | t=9.574 df=10 | Not determined | Not determined |
| eEPSP amplitude (normalized) | 100.0 ± 5.986 (6) | 79.15 ± 5.682 (6) | 0.0411* | U = 5 | 0.0301* | t=2.527 df=10 | Not determined | Not determined |
| quantal content (normalized) | 100.0 ± 3.668 (6) | 155.9 ± 10.94 (6) | 0.0022** | U = 0 | 0.0007*** | t=4.842 df=10 | Not determined | Not determined |
| ***srpk79D* ATC** | 6 cells from 3 animals | 6 cells from 3 animals | | | | | | |
| mEPSP amplitude (normalized) | 100.0 ± 4.966 (5) | 50.68 ± 1.664 (6) | 0.0043** | U = 0 | < 0.0001*** | t=10.18 df=9 | Not determined | Not determined |
| eEPSP amplitude (normalized) | 100.0 ± 3.269 (5) | 84.59 ± 3.852 (6) | 0.0519 | U = 4 | 0.0156 | t=2.974 df=9 | Not determined | Not determined |
| quantal content (normalized) | 100.0 ± 5.632 (5) | 166.2 ± 8.857 (6) | 0.0043** | U = 0 | 0.0002*** | t=5.997 df=9 | Not determined | Not determined |

| Normalized comparison of PhTx treatment Figure 5c | | | Mann-Whitney U-test (P values) two-tailed | test statistic | Unpaired t test (P values) two-tailed | test statistic | D'Agostino & Pearson omnibus normality test | |
|---|---|---|---|---|---|---|---|---|
| | **Wild-type +PhTx (n)** | ***srpk79D* ATC +PhTx (n)** | **Wild-type +PhTx vs. *srpk79D* ATC +PhTx** | | **Wild-type +PhTx vs. *srpk79D* ATC +PhTx** | | **Wild-type +PhTx** | ***srpk79D* ATC +PhTx** |
| mEPSP amplitude (normalized) | 51.15 ± 2.349 (6) | 50.68 ± 1.664 (6) | 0.5584 | U = 14 | 0.8745 | t=0.1620 df=10 | Not determined | Not determined |
| eEPSP amplitude (normalized) | 79.15 ± 5.682 (6) | 84.59 ± 3.852 (6) | 0.4848 | U = 13 | 0.446 | t=0.7933 df=10 | Not determined | Not determined |
| quantal content (normalized) | 155.9 ± 10.94 (6) | 166.2 ± 8.857 (6) | 0.6991 | U =15 | 0.4805 | t=0.7327 df=10 | Not determined | Not determined |

| Parameter<br>(Figure) | | | Mann-Whitney U-test (P values)<br>two-tailed | test statistic | Unpaired t test (P values)<br>two-tailed | test statistic | D'Agostino & Pearson omnibus normality test | |
|---|---|---|---|---|---|---|---|---|
| **Chronic Homeostatic Plasticity**<br>**(Figure 5f)** | | | | | | | | |
| **Wild-type vs. *gluRIIA* $^{Null}$** | **Wild-type (n)** | ***gluRIIA* $^{Null}$ (n)** | | | | | **Wild-type** | ***gluRIIA* $^{Null}$** |
| mEPSP amplitude (mV) | 0.9406 ± 0.03027 (9) | 0.4883 ± 0.02990 (8) | < 0.0001*** | U = 0 | < 0.0001*** | t=10.58 df=15 | 0.3935 | 0.527 |
| eEPSP amplitude (mV) | 33.82 ± 0.7550 (9) | 26.67 ± 0.9317 (8) | 0.0002 *** | U = 1 | < 0.0001*** | t=6.017 df=15 | 0.5113 | 0.447 |
| quantal content | 36.18 ± 1.147 (9) | 59.04 ± 2.815 (8) | < 0.0001*** | U = 0 | < 0.0001*** | t=7.853 df=15 | 0.844 | 0.4393 |
| ***brp* $^{Null}$ vs. *brp* $^{Null}$;*gluRIIA* $^{Null}$** | ***brp* $^{Null}$ (n)** | ***brp* $^{Null}$;*gluRIIA* $^{Null}$ (n)** | | | | | ***brp* $^{Null}$** | ***brp* $^{Null}$;*gluRIIA* $^{Null}$** |
| mEPSP amplitude (mV) | 1.073 ± 0.04019 (8) | 0.4976 ± 0.01917 (9) | < 0.0001*** | U = 0 | < 0.0001*** | t=13.42 df=15 | 0.4563 | 0.5221 |
| eEPSP amplitude (mV) | 25.52 ± 1.166 (8) | 14.79 ± 1.163 (9) | < 0.0001*** | U = 0 | < 0.0001*** | t=6.492 df=15 | 0.5182 | 0.8105 |
| quantal content | 23.92 ± 1.157 (8) | 29.61 ± 1.792 (9) | 0.0274* | U = 13 | 0.0204* | t=2.593 df=15 | 0.4079 | 0.9408 |
| **(Normalized)** | | | | | | | | |
| **Wild-type vs. *gluRIIA* $^{Null}$** | **Wild-type (n)** | ***gluRIIA* $^{Null}$ (n)** | | | | | | |
| mEPSP amplitude (normalized) | 100.0 ± 3.218 (9) | 51.91 ± 3.179 (8) | < 0.0001*** | U = 0 | < 0.0001*** | t=10.58 df=15 | 0.3935 | 0.527 |
| eEPSP amplitude (normalized) | 100.0 ± 2.233 (9) | 78.86 ± 2.755 (8) | 0.0002 *** | U = 1 | < 0.0001*** | t=6.017 df=15 | 0.5113 | 0.447 |
| quantal content (normalized) | 100.0 ± 3.169 (9) | 163.2 ± 7.781 (8) | < 0.0001*** | U = 0 | < 0.0001*** | t=7.853 df=15 | 0.844 | 0.4393 |
| ***brp* $^{Null}$ vs. *brp* $^{Null}$;*gluRIIA* $^{Null}$** | ***brp* $^{Null}$ (n)** | ***brp* $^{Null}$;*gluRIIA* $^{Null}$ (n)** | | | | | ***brp* $^{Null}$** | ***brp* $^{Null}$;*gluRIIA* $^{Null}$** |
| mEPSP amplitude (normalized) | 100.0 ± 3.745 (8) | 46.37 ± 1.786 (9) | < 0.0001*** | U = 0 | < 0.0001*** | t=13.42 df=15 | 0.4563 | 0.5221 |
| eEPSP amplitude (normalized) | 100.0 ± 4.567 (8) | 57.96 ± 4.559 (9) | < 0.0001*** | U = 0 | < 0.0001*** | t=6.492 df=15 | 0.5182 | 0.8105 |
| quantal content (normalized) | 100.0 ± 4.837 (8) | 123.8 ± 7.494 (9) | 0.0274* | U = 13 | 0.0204* | t=2.593 df=15 | 0.4079 | 0.9408 |

| Normalized comparison | | | Mann-Whitney U-test (P values)<br>two-tailed | test statistic | Unpaired t test (P values)<br>two-tailed | test statistic | D'Agostino & Pearson omnibus normality test | |
|---|---|---|---|---|---|---|---|---|
| **(Figure 5f)** | ***gluRIIA* $^{Null}$ (n)** | ***brp* $^{Null}$;*gluRIIA* $^{Null}$ (n)** | ***gluRIIA* $^{Null}$ vs. *brp* $^{Null}$;*gluRIIA* $^{Null}$** | | ***gluRIIA* $^{Null}$ vs. *brp* $^{Null}$;*gluRIIA* $^{Null}$** | | ***gluRIIA* $^{Null}$ (n)** | ***brp* $^{Null}$;*gluRIIA* $^{Null}$** |
| mEPSP amplitude (normalized) | 51.91 ± 3.179 (8) | 46.37 ± 1.786 (9) | 0.2359 | U = 23 | 0.1378 | t=1.568 df=15 | 0.527 | 0.5221 |
| eEPSP amplitude (normalized) | 78.86 ± 2.755 (8) | 57.96 ± 4.559 (9) | 0.0079** | U = 9 | 0.0017** | t=3.801 df=15 | 0.447 | 0.8105 |
| quantal content (normalized) | 163.2 ± 7.781 (8) | 123.8 ± 7.494 (9) | 0.0055** | U = 8 | 0.0024** | t=3.641 df=15 | 0.4393 | 0.9409 |

| Parameter<br>(Figure) | | | Mann-Whitney U-test (P values)<br>two-tailed | test statistic | Unpaired t test (P values)<br>two-tailed | test statistic | D'Agostino & Pearson omnibus normality test<br>Wild-type | ***gluRIIA* $^{Null}$** |
|---|---|---|---|---|---|---|---|---|
| **(Figure 5g)** | | | | | | | | |
| **Wild-type vs. *gluRIIA* $^{Null}$** | **Wild-type (n)** 6 cells from 4 animals | ***gluRIIA* $^{Null}$ (n)** 8 cells from 4 animals | | | | | | |
| mEPSP amplitude (mV) | 0.8308 ± 0.09016 (6) | 0.3159 ± 0.01962 (8) | 0.0007*** | U = 0 | < 0.0001*** | t=6.410 df=12 | Not determined | 0.43 |
| eEPSP amplitude (mV) | 24.17 ± 2.252 (6) | 20.90 ± 1.989 (8) | 0.345 | U = 16 | 0.3003 | t=1.082 df=12 | Not determined | 0.3339 |
| quantal content | 30.58 ± 4.134 (6) | 67.60 ± 7.382 (8) | 0.0047** | U = 3 | 0.0018** | t=3.978 df=12 | Not determined | 0.8133 |
| ***srpk79D* $^{ATC}$ vs. *srpk79D* $^{ATC}$;*gluRIIA* $^{Null}$** | ***srpk79D* $^{ATC}$ (n)** 6 cells from 4 animals | ***srpk79D* $^{ATC}$;*gluRIIA* $^{Null}$ (n)** 8 cells from 4 animals | | | | | ***srpk79D* $^{ATC}$** | ***srpk79D* $^{ATC}$;*gluRIIA* $^{Null}$** |
| mEPSP amplitude (mV) | 0.9919 ± 0.1107 (6) | 0.3638 ± 0.04587 (8) | 0.0007*** | U = 0 | < 0.0001*** | t=5.783 df=12 | Not determined | 0.3573 |
| eEPSP amplitude (mV) | 35.22 ± 3.728 (6) | 10.14 ± 1.743 (8) | 0.0007*** | U = 0 | < 0.0001*** | t=6.639 df=12 | Not determined | 0.4622 |
| quantal content | 37.68 ± 6.060 (6) | 32.84 ± 7.563 (8) | 0.5728 | U = 19 | 0.6448 | t=0.4728 df=12 | Not determined | 0.4653 |
| **(Normalized)** | | | | | | | | |
| **Wild-type vs. *gluRIIA* $^{Null}$** | **Wild-type (n)** 6 cells from 4 animals | ***gluRIIA* $^{Null}$ (n)** 8 cells from 4 animals | | | | | **Wild-type** | ***gluRIIA* $^{Null}$** |
| mEPSP amplitude (normalized) | 100.0 ± 10.85 (6) | 38.02 ± 2.362 (8) | 0.0007*** | U = 0 | < 0.0001*** | t=6.410 df=12 | Not determined | 0.43 |
| eEPSP amplitude (normalized) | 100.0 ± 9.321 (6) | 86.50 ± 8.230 (8) | 0.345 | U = 16 | 0.3003 | t=1.082 df=12 | Not determined | 0.3339 |
| quantal content (normalized) | 100.0 ± 13.52 (6) | 221.1 ± 24.14 (8) | 0.0047** | U = 3 | 0.0018** | t=3.978 df=12 | Not determined | 0.8133 |
| ***srpk79D* $^{ATC}$ vs. *srpk79D* $^{ATC}$;*gluRIIA* $^{Null}$** | ***srpk79D* $^{ATC}$ (n)** 6 cells from 4 animals | ***srpk79D* $^{ATC}$;*gluRIIA* $^{Null}$ (n)** 8 cells from 4 animals | | | | | ***srpk79D* $^{ATC}$** | ***srpk79D* $^{ATC}$;*gluRIIA* $^{Null}$** |
| mEPSP amplitude (normalized) | 100.0 ± 11.16 (6) | 36.68 ± 4.624 (8) | 0.0007*** | U = 0 | < 0.0001*** | t=5.783 df=12 | Not determined | 0.3573 |
| eEPSP amplitude (normalized) | 100.0 ± 10.59 (6) | 28.79 ± 4.950 (8) | 0.0007*** | U = 0 | < 0.0001*** | t=6.639 df=12 | Not determined | 0.4622 |
| quantal content (normalized) | 100.0 ± 16.08 (6) | 87.16 ± 20.07 (8) | 0.5728 | U = 19 | 0.6448 | t=0.4728 df=12 | Not determined | 0.4653 |

| Normalized comparison<br>Figure 5g | | | Mann-Whitney U-test (P values)<br>two-tailed | test statistic | Unpaired t test (P values)<br>two-tailed | test statistic | D'Agostino & Pearson omnibus normality test | |
|---|---|---|---|---|---|---|---|---|
| | ***gluRIIA* $^{Null}$ (n)** | ***srpk79D* $^{ATC}$;*gluRIIA* $^{Null}$ (n)** | ***gluRIIA* $^{Null}$ vs. *srpk79D* $^{ATC}$;*gluRIIA* $^{Null}$** | | ***gluRIIA* $^{Null}$ vs. *srpk79D* $^{ATC}$;*gluRIIA* $^{Null}$** | | ***gluRIIA* $^{Null}$** | ***srpk79D* $^{ATC}$;*gluRIIA* $^{Null}$** |
| mEPSP amplitude (normalized) | 38.02 ± 2.362 (8) | 36.68 ± 4.624 (8) | 0.3282 | U = 22 | 0.7997 | t=0.2586 df=14 | 0.43 | 0.3573 |
| eEPSP amplitude (normalized) | 86.50 ± 8.230 (8) | 28.79 ± 4.950 (8) | 0.0006*** | U = 2 | < 0.0001*** | t=6.010 df=14 | 0.3339 | 0.4622 |
| quantal content (normalized) | 221.1 ± 24.14 (8) | 87.16 ± 20.07 (8) | 0.0011** | U = 3 | 0.0008*** | t=4.266 df=14 | 0.8133 | 0.4653 |

| Parameter (Figure) | | | Mann-Whitney U-test (P values) two-tailed | test statistic | Unpaired t test (P values) two-tailed | test statistic | D'Agostino & Pearson omnibus normality test | |
|---|---|---|---|---|---|---|---|---|
| **(Supplementary Figure 7d)** | | | | | | | | |
| **Wild-type vs. *gluRIIA*$^{Null}$** | **Wild-type (n)** 15 cells from 10 animals | ***gluRIIA*$^{Null}$ (n)** 17 cells from 11 animals | | | | | **Wild-type** | ***gluRIIA*$^{Null}$** |
| mEPSP amplitude (mV) | 0.8135 ± 0.04911 (15) | 0.3533 ± 0.01729 (17) | < 0.0001*** | U = 0 | < 0.0001*** | t=9.281 df=30 | 0.498 | 0.2166 |
| eEPSP amplitude (mV) | 24.19 ± 2.321 (15) | 18.28 ± 2.455 (17) | 0.105 | U = 84 | 0.0929 | t=1.736 df=30 | 0.5692 | 0.6495 |
| quantal content | 31.03 ± 3.411 (15) | 54.32 ± 8.195 (17) | 0.0402* | U = 73 | 0.018* | t=2.503 df=30 | 0.1465 | 0.031* |
| ***aplip-1*$^{ek4}$ vs. *aplip-1*$^{ek4}$;*gluRIIA*$^{Null}$** | ***aplip-1*$^{ek4}$ (n)** 18 cells from 11 animals | ***aplip-1*$^{ek4}$;*gluRIIA*$^{Null}$ (n)** 20 cells from 13 animals | | | | | ***aplip-1*$^{ek4}$** | ***aplip-1*$^{ek4}$;*gluRIIA*$^{Null}$ (n)** |
| mEPSP amplitude (mV) | 0.9249 ± 0.08341 (18) | 0.3462 ± 0.02172 (20) | < 0.0001*** | U = 3 | < 0.0001*** | t=7.035 df=36 | 0.1945 | 0.1951 |
| eEPSP amplitude (mV) | 22.31 ± 2.441 (18) | 14.03 ± 2.040 (20) | 0.0173* | U = 99 | 0.0128* | t=2.619 df=36 | 0.7679 | 0.0783 |
| quantal content | 26.96 ± 3.742 (18) | 41.64 ± 6.220 (20) | 0.0873 | U = 121 | 0.0568 | t=1.968 df=36 | 0.0571 | 0.2157 |
| **(Normalized)** | | | | | | | | |
| **Wild-type vs. *gluRIIA*$^{Null}$** | **Wild-type (n)** 15 cells from 10 animals | ***gluRIIA*$^{Null}$ (n)** 17 cells from 11 animals | | | | | **Wild-type** | ***gluRIIA*$^{Null}$** |
| mEPSP amplitude (normalized) | 100.0 ± 6.037 (15) | 43.43 ± 2.125 (17) | < 0.0001*** | U = 0 | < 0.0001*** | t=9.281 df=30 | 0.498 | 0.2166 |
| eEPSP amplitude (normalized) | 100.0 ± 9.592 (15) | 75.57 ± 10.15 (17) | 0.105 | U = 84 | 0.0929 | t=1.736 df=30 | 0.5692 | 0.6495 |
| quantal content (normalized) | 100.0 ± 10.99 (15) | 175.1 ± 26.41 (17) | 0.0402* | U = 73 | 0.018* | t=2.503 df=30 | 0.1465 | 0.031* |
| ***aplip-1*$^{ek4}$ vs. *aplip-1*$^{ek4}$;*gluRIIA*$^{Null}$** | ***aplip-1*$^{ek4}$ (n)** 18 cells from 11 animals | ***aplip-1*$^{ek4}$;*gluRIIA*$^{Null}$ (n)** 20 cells from 13 animals | | | | | ***aplip-1*$^{ek4}$** | ***aplip-1*$^{ek4}$;*gluRIIA*$^{Null}$ (n)** |
| mEPSP amplitude (normalized) | 100.0 ± 9.018 (18) | 37.43 ± 2.348 (20) | < 0.0001*** | U = 3 | < 0.0001*** | t=7.035 df=36 | 0.1945 | 0.1951 |
| eEPSP amplitude (normalized) | 100.0 ± 10.94 (18) | 62.91 ± 9.145 (20) | 0.0173* | U = 99 | 0.0128* | t=2.619 df=36 | 0.7679 | 0.0783 |
| quantal content (normalized) | 100.0 ± 13.88 (18) | 154.5 ± 23.08 (20) | 0.0873 | U = 121 | 0.0568 | t=1.968 df=36 | 0.0571 | 0.2157 |

| Normalized comparison Supplementary Fig. 7d | | | Mann-Whitney U-test (P values) two-tailed | test statistic | Unpaired t test (P values) two-tailed | test statistic | D'Agostino & Pearson omnibus normality test | |
|---|---|---|---|---|---|---|---|---|
| | ***gluRIIA*$^{Null}$ (n)** | ***aplip-1*$^{ek4}$;*gluRIIA*$^{Null}$ (n)** | ***gluRIIA*$^{Null}$ vs. *aplip-1*$^{ek4}$;*gluRIIA*$^{Null}$** | | ***gluRIIA*$^{Null}$ vs. *aplip-1*$^{ek4}$;*gluRIIA*$^{Null}$** | | ***gluRIIA*$^{Null}$** | ***aplip-1*$^{ek4}$;*gluRIIA*$^{Null}$ (n)** |
| mEPSP amplitude (normalized) | 43.43 ± 2.125 (17) | 37.43 ± 2.348 (20) | 0.0416* | U = 103 | 0.0705 | t=1.866 df=35 | 0.2166 | 0.1951 |
| eEPSP amplitude (normalized) | 75.57 ± 10.15 (17) | 62.91 ± 9.145 (20) | 0.3413 | U = 138 | 0.3592 | t=0.9290 df=35 | 0.6495 | 0.0783 |
| quantal content (normalized) | 175.1 ± 26.41 (17) | 154.5 ± 23.08 (20) | 0.4975 | U = 147 | 0.5587 | t=0.5904 df=35 | 0.031* | 0.2157 |

| Parameter (Figure) | | | Mann-Whitney U-test (P values) two-tailed | test statistic | Unpaired t test (P values) two-tailed | test statistic | D'Agostino & Pearson omnibus normality test | |
|---|---|---|---|---|---|---|---|---|
| **(Figure 1f)** | **Wild-type (n)** | ***gluRIIA*$^{Null}$ (n)** | | | | | | |
| **Wild-type vs. *gluRIIA*$^{Null}$** | 10 cells from 6 animals | 9 cells from 5 animals | | | | | **Wild-type** | ***gluRIIA*$^{Null}$** |
| mEPSP amplitude (mV) | 1.144 ± 0.06038 (10) | 0.3680 ± 0.02385 (9) | < 0.0001*** | U = 0 | < 0.0001*** | t=11.47 df=17 | 0.4692 | 0.324 |
| eEPSP amplitude (mV) | 22.17 ± 1.759 (10) | 21.09 ± 1.771 (9) | 0.7197 | U = 40 | 0.671 | t=0.4323 df=17 | 0.6419 | 0.4868 |
| quantal content | 19.99 ± 2.005 (10) | 60.95 ± 8.605 (9) | < 0.0001*** | U = 0 | 0.0001*** | t=4.871 df=17 | 0.5887 | 0.1365 |

| Parameter (Figure) | control treatment (n) | PhTx treatment (n) | Mann-Whitney U-test (P values) two-tailed | test statistic | Unpaired t test (P values) two-tailed | test statistic | D'Agostino & Pearson omnibus normality test control treatment | PhTx treatment |
|---|---|---|---|---|---|---|---|---|
| **Induction of Rapid Homeostatic Plasticity after translation block (Supplementary Figure 5b)** | **control treatment (n)** | **PhTx treatment (n)** | | | | | **control treatment** | **PhTx treatment** |
| **Wild-type - Cycloheximide** | 10 cells from 5 animals | 14 cells from 7 animals | | | | | | |
| mEPSP amplitude (mV) | 0.912 ± 0.043 (10) | 0.465 ± 0.028 (7) | 0.0001*** | U = 0 | < 0.0001*** | t=7.750 df=15 | 0.9518 | Not determined |
| eEPSP amplitude (mV) | 32.50 ± 0.715 (10) | 33.55 ± 1.3 (7) | 0.228 | U = 22 | 0.4569 | t=0.7636 df=15 | 0.5691 | Not determined |
| quantal content | 36.15 ± 1.317 (10) | 73.22 ± 4.236 (7) | 0.0001*** | U = 0 | < 0.0001*** | t=9.659 df=15 | 0.5033 | Not determined |
| **Wild-type + Cycloheximide** | | | | | | | | |
| mEPSP amplitude (mV) | 0.9050 ± 0.05065 (12) | 0.4375 ± 0.01526 (10) | < 0.0001*** | U = 0 | < 0.0001*** | t=8.142 df=20 | 0.3425 | Not determined |
| eEPSP amplitude (mV) | 33.29 ± 1.061 (12) | 30.41 ± 1.593 (10) | 0.203 | u = 40 | 0.1368 | t=1.550 df=20 | 0.2946 | Not determined |
| quantal content | 38.09 ± 2.450 (12) | 69.92 ± 3.807 (10) | < 0.0001*** | U = 0 | < 0.0001*** | t=7.262 df=20 | 0.7881 | Not determined |
| **(Normalized)** | | | | | | | | |
| **Wild-type - Cycloheximide** | | | | | | | | |
| mEPSP amplitude (mV) | 100.0 ± 4.788 (10) | 51.05 ± 3.090 (7) | 0.0001*** | U = 0 | < 0.0001*** | t=7.750 df=15 | 0.9518 | Not determined |
| eEPSP amplitude (mV) | 100.0 ± 2.200 (10) | 103.2 ± 3.999 (7) | 0.228 | U = 22 | 0.4569 | t=0.7636 df=15 | 0.5691 | Not determined |
| quantal content | 100.0 ± 3.642 (10) | 202.5 ± 11.72 (7) | 0.0001*** | U = 0 | < 0.0001*** | t=9.659 df=15 | 0.5033 | Not determined |
| **Wild-type + Cycloheximide** | | | | | | | | |
| mEPSP amplitude (mV) | 100.0 ± 5.597 (12) | 48.34 ± 1.686 (10) | < 0.0001*** | U = 0 | < 0.0001*** | t=8.142 df=20 | 0.3425 | Not determined |
| eEPSP amplitude (mV) | 100.0 ± 3.187 (12) | 91.34 ± 4.787 (10) | 0.203 | u = 40 | 0.1368 | t=1.550 df=20 | 0.2946 | Not determined |
| quantal content | 100.0 ± 6.432 (12) | 183.6 ± 9.995 (10) | < 0.0001*** | U = 0 | < 0.0001*** | t=7.262 df=20 | 0.7881 | Not determined |

| Normalized comparison Supplementary Fig. 5b | | | Mann-Whitney U-test (P values) two-tailed | test statistic | Unpaired t test (P values) two-tailed | test statistic | D'Agostino & Pearson omnibus normality test | |
|---|---|---|---|---|---|---|---|---|
| | **-Cycloheximide; +PhTx** | **+Cycloheximide; +PhTx** | **-Cycloheximide; +PhTx vs. +Cycloheximide; +PhTx** | | | | **-Cycloheximide; +PhTx** | **+Cycloheximide; +PhTx** |
| mEPSP amplitude (mV) | 51.05 ± 3.090 (7) | 48.34 ± 1.686 (10) | 0.4599 | U = 27 | 0.4197 | t=0.8297 df=15 | Not determined | 0.6599 |
| eEPSP amplitude (mV) | 103.2 ± 3.999 (7) | 91.34 ± 4.787 (10) | 0.1329 | U = 19 | 0.0941 | t=1.787 df=15 | Not determined | 0.5259 |
| quantal content | 202.5 ± 11.72 (7) | 183.6 ± 9.995 (10) | 0.2393 | U = 23 | 0.2393 | t=1.226 df=15 | Not determined | 0.5787 |

| **(Supplementary Figure 10k)** | | | | | | | | |
|---|---|---|---|---|---|---|---|---|
| **Wild-type ± Cycloheximide without PhTx (absolute values)** | **-Cycloheximide; ctrl** | **+Cycloheximide; ctrl** | **-Cycloheximide; ctrl vs. +Cycloheximide; ctrl** | | | | **-Cycloheximide; ctrl** | **+Cycloheximide; ctrl** |
| mEPSP amplitude (mV) | 0.9123 ± 0.04368 (10) | 0.9050 ± 0.05065 (12) | 0.8588 | U = 57 | 0.9161 | t=0.1067 df=20 | 0.9518 | 0.3425 |
| eEPSP amplitude (mV) | 32.50 ± 0.7150 (10) | 33.29 ± 1.061 (12) | 0.5387 | U = 50 | 0.5634 | t=0.5876 df=20 | 0.5691 | 0.2946 |
| quantal content | 36.15 ± 1.317 (10) | 38.09 ± 2.450 (12) | 0.6744 | U = 53 | 0.5196 | t=0.6556 df=20 | 0.5033 | 0.7881 |

| Parameter (Figure) | control treatment (n) (n = number of NMJs from min. 5 animals) | PhTx treatment (n) | Mann-Whitney U-test (P values) two-tailed | test statistic | Unpaired t test (P values) two-tailed | test statistic | D'Agostino & Pearson omnibus normality test control treatment | PhTx treatment |
|---|---|---|---|---|---|---|---|---|
| **Synaptic intensity (% of Ctrl)** | | | | | | | | |
| **(Figure 1d)** | | | | | | | | |
| **Wild-type** | | | | | | | | |
| BRP | 100 ± 7.142 (21) | 146.5 ± 11.60 (23) | 0.0044** | U = 122 | 0.0018** | t=3.335 df=42 | 0.0058* | 0.5625 |
| RBP | 100 ± 5.376 (44) | 130.9 ± 8.438 (39) | 0.0075** | U = 567 | 0.0022** | t=3.157 df=81 | 0.1419 | 0.1156 |
| Unc13A | 100 ± 6.549 (21) | 158.9 ± 10.27 (23) | 0.0002*** | U = 88 | < 0.0001*** | t=4.730 df=42 | < 0.0001* | 0.8957 |
| Syx-1A | 100 ± 9.299 (43) | 164.2 ± 15.87 (36) | < 0.0001*** | U = 377 | 0.0005*** | t=3.625 df=77 | < 0.0001* | 0.0001* |
| Unc18 | 100 ± 7.119 (16) | 105.2 ± 6.831 (24) | 0.6835 | U = 177 | 0.6152 | t=0.5069 df=38 | 0.0328* | 0.0112* |
| **Synaptic intensity ((a.u.); not normalized)** | | | | | | | | |
| **(Figure 1d)** | | | | | | | | |
| **Wild-type** | | | | | | | | |
| BRP | 641.9 ± 45.84 (21) | 940.3 ± 74.49 (23) | 0.0044** | U = 122 | 0.0018** | t=3.335 df=42 | 0.0058* | 0.5625 |
| RBP | 797.9 ± 42.90 (44) | 1044 ± 67.33 (39) | 0.0075** | U = 567 | 0.0022** | t=3.157 df=81 | 0.1419 | 0.1156 |
| Unc13A | 530.5 ± 34.74 (21) | 842.7 ± 54.48 (23) | 0.0002*** | U = 88 | < 0.0001*** | t=4.730 df=42 | < 0.0001* | 0.8957 |
| Syx-1A | 708.7 ± 65.90 (43) | 1164 ± 112.5 (36) | < 0.0001*** | U = 377 | 0.0005*** | t=3.625 df=77 | < 0.0001* | 0.0001* |
| Unc18 | 1247 ± 88.76 (16) | 1311 ± 85.16 (24) | 0.6835 | U = 177 | 0.6152 | t=0.5069 df=38 | 0.0328* | 0.0112* |

| Synaptic intensity (% of Ctrl) (Supplementary Figure 1a) | control treatment (n) (n = number of NMJs from min. 5 animals) | PhTx treatment (n) | Mann-Whitney U-test (P values) two-tailed | test statistic | Unpaired t test (P values) two-tailed | test statistic | D'Agostino & Pearson omnibus normality test control treatment | PhTx treatment |
|---|---|---|---|---|---|---|---|---|
| **Ok6::Syd-1-GFP** | | | | | | | | |
| BRP | 100 ± 7.809 (21) | 153 ± 7.676 (29) | < 0.0001*** | U = 93 | < 0.0001*** | t=4.726 df=48 | 0.0004*** | 0.2676 |
| GFP | 100 ± 8.853 (21) | 82.58 ± 6.056 (29) | 0.107 | U = 222 | 0.099 | t=1.682 df=48 | 0.6238 | 0.0096** |

| | | |
|---|---|---|
| **(Figure Supplementary 1b)** | | |
| **BRP vs RBP** | | |
| **BRP** | | |
| Bin1 | 360.754 ± 18.436 (44) | 498.595 ± 39.363 (39) |
| Bin2 | 540.761 ± 27.753 (44) | 755.233 ± 57.095 (39) |
| Bin3 | 672.746 ± 34.168 (44) | 954.003 ± 70.367 (39) |
| Bin4 | 833.531 ± 43.065 (44) | 1191.625 ± 84.539 (39) |
| Bin5 | 1130.9 ± 60.515 (44) | 1639.203 ± 109.971 (39) |
| **RBP** | | |
| Bin1 | 499.437 ± 28.644 (44) | 662.895 ± 45.753 (39) |
| Bin2 | 651.425 ± 34.880 (44) | 854.929 ± 57.835 (39) |
| Bin3 | 766.809 ± 42.111 (44) | 1013.391 ± 67.368 (39) |
| Bin4 | 919.854 ± 48.940 (44) | 1180.286 ± 79.251 (39) |
| Bin5 | 1155.559 ± 63.923 (44) | 1525.522 ± 97.061 (39) |
| **BRP vs Unc13A** | | |
| **BRP** | | |
| Bin1 | 331.889 ± 24.125 (21) | 472.314 ± 35.164 (23) |
| Bin2 | 479.109 ± 35.612 (21) | 704.125 ± 55.224 (23) |
| Bin3 | 598.144 ± 42.557 (21) | 880.377 ± 69.689 (23) |
| Bin4 | 732.988 ± 52.437 (21) | 1092.994 ± 87.141 (23) |
| Bin5 | 331.889 ± 24.125 (21) | 472.314 ± 35.164 (23) |
| **Unc13A** | | |
| Bin1 | 295.701 ± 22.436 (21) | 505.049 ± 37.489 (23) |
| Bin2 | 406.786 ± 34.036 (21) | 679.722 ± 49.715 (23) |
| Bin3 | 499.974 ± 28.362 (21) | 787.825 ± 50.854 (23) |
| Bin4 | 613.045 ± 41.181 (21) | 942.936 ± 56.869 (23) |
| Bin5 | 807.655 ± 55.735 (21) | 1253.008 ± 86.463 (23) |
| **BRP vs Syx-1A** | | |
| **BRP** | | |
| Bin1 | 336.700 ± 21.098 (43) | 369.202 ± 21.468 (36) |
| Bin2 | 506.696 ± 34.442 (43) | 607.061 ± 41.536 (36) |
| Bin3 | 642.368 ± 44.858 (43) | 808.261 ± 58.731 (36) |
| Bin4 | 817.806 ± 58.853 (43) | 1043.762 ± 76.979 (36) |
| Bin5 | 1129.353 ± 83.030 (43) | 1523.703 ± 113.665 (36) |
| **Syx-1A** | | |
| Bin1 | 494.859 ± 45.419 (43) | 797.197 ± 86.019 (36) |
| Bin2 | 613.611 ± 59.468 (43) | 980.762 ± 99.145 (36) |
| Bin3 | 684.983 ± 65.945 (43) | 1141.57 ± 113.877 (36) |
| Bin4 | 784.569 ± 74.178 (43) | 1303.114 ± 129.372 (36) |
| Bin5 | 960.462 ± 87.523 (43) | 1594.776 ± 140.524 (36) |

| Parameter (Figure) | Wild-type (n) (n = number of NMJs from at least 4 animals) | *gluRIIA* $^{Null}$ (n) | Mann-Whitney U-test (P values) two-tailed | test statistic | Unpaired t test (P values) two-tailed | test statistic | D'Agostino & Pearson omnibus normality test Wild-type | gluRIIA $^{Null}$ |
|---|---|---|---|---|---|---|---|---|
| **Synaptic intensity (% of Ctrl) (Figure 1h)** | | | | | | | | |
| BRP | 100 ± 7.692 (9) | 203.1 ± 22.17 (12) | 0.0007*** | U = 9 | 0.001** | t=3.876 df=19 | 0.7022 | 0.5131 |
| RBP | 100 ± 9.730 (19) | 172.3 ± 9.602 (21) | < 0.0001*** | U = 43 | < 0.0001*** | t=5.279 df=38 | 0.0139* | 0.0559 |
| Unc13A | 100 ± 7.737 (9) | 387.8 ± 36.78 (12) | < 0.0001*** | U = 1 | < 0.0001*** | t=6.652 df=19 | 0.4247 | 0.7757 |
| Syx-1A | 100 ± 5.708 (19) | 200.7 ± 18.39 (22) | < 0.0001*** | U = 25 | < 0.0001*** | t=4.909 df=39 | 0.4026 | 0.0139* |
| Unc18 | 100 ± 8.317 (14) | 694.9 ± 54.63 (15) | < 0.0001*** | U = 0 | < 0.0001*** | t=10.40 df=27 | 0.465 | 0.5875 |
| **Synaptic intensity ((a.u.); not normalized) (Figure 1h)** | | | | | | | | |
| BRP | 769.5 ± 59.19 (9) | 1563 ± 170.6 (12) | 0.0007*** | U = 9 | 0.001** | t=3.876 df=19 | 0.7022 | 0.5131 |
| RBP | 661.6 ± 64.38 (19) | 1140 ± 63.53 (21) | < 0.0001*** | U = 43 | < 0.0001*** | t=5.279 df=38 | 0.0139* | 0.0559 |
| Unc13A | 407.5 ± 31.53 (9) | 1580 ± 149.9 (12) | < 0.0001*** | U = 1 | < 0.0001*** | t=6.652 df=19 | 0.4247 | 0.7757 |
| Syx-1A | 656.8 ± 37.49 (19) | 1318 ± 120.8 (22) | < 0.0001*** | U = 25 | < 0.0001*** | t=4.909 df=39 | 0.4026 | 0.0139* |
| Unc18 | 200.5 ± 16.68 (14) | 1394 ± 109.5 (15) | < 0.0001*** | U = 0 | < 0.0001*** | t=10.40 df=27 | 0.465 | 0.5875 |
| **(Supplementary Figure 1c)** | | | | | | | | |
| **BRP vs RBP** | | | | | | | | |
| **BRP** | | | | | | | | |
| Bin1 | 427.129 ± 40.291 (19) | 642.663 ± 42.667 (21) | | | | | | |
| Bin2 | 615.213 ± 54.816 (19) | 983.311 ± 65.926 (21) | | | | | | |
| Bin3 | 751.216 ± 68.602 (19) | 1235.075 ± 82.487 (21) | | | | | | |
| Bin4 | 912.767 ± 85.687 (19) | 1543.867 ± 99.662 (21) | | | | | | |
| Bin5 | 1243.825 ± 119.252 (19) | 2106.347 ± 124.351 (21) | | | | | | |
| **RBP** | | | | | | | | |
| Bin1 | 381.347 ± 40.370 (19) | 650.847 ± 42.166 (21) | | | | | | |
| Bin2 | 534.336 ± 53.122 (19) | 890.064 ± 55.963 (21) | | | | | | |
| Bin3 | 636.233 ± 61.847 (19) | 1084.65 ± 66.285 (21) | | | | | | |
| Bin4 | 754.413 ± 74.695 (19) | 1325.438 ± 77.998 (21) | | | | | | |
| Bin5 | 1000.198 ± 97.377 (19) | 1746.62 ± 89.139 (21) | | | | | | |
| **BRP vs Unc13A** | | | | | | | | |
| **BRP** | | | | | | | | |
| Bin1 | 385.266 ± 31.484 (9) | 706.975 ± 89.879 (12) | | | | | | |
| Bin2 | 580.696 ± 47.507 (9) | 1172.313 ± 144.838 (12) | | | | | | |
| Bin3 | 725.251 ± 56.315 (9) | 1489.847 ± 167.055 (12) | | | | | | |
| Bin4 | 886.716 ± 68.038 (9) | 1843.5 ± 201.659 (12) | | | | | | |
| Bin5 | 1187.099 ± 91.663 (9) | 2478.65 ± 253.168 (12) | | | | | | |
| **Unc13A** | | | | | | | | |
| Bin1 | 205.651 ± 19.890 (9) | 727.054 ± 72.383 (12) | | | | | | |
| Bin2 | 333.052 ± 30.748 (9) | 1221.429 ± 132.941 (12) | | | | | | |
| Bin3 | 390.111 ± 25.176 (9) | 1555.63 ± 167.858 (12) | | | | | | |
| Bin4 | 456.629 ± 33.371 (9) | 1873.615 ± 179.756 (12) | | | | | | |
| Bin5 | 619.255 ± 55.144 (9) | 2425.303 ± 218.619 (12) | | | | | | |
| **BRP vs Syx-1A** | | | | | | | | |
| **BRP** | | | | | | | | |
| Bin1 | 266.204 ± 35.960 (19) | 505.015 ± 24.888 (22) | | | | | | |
| Bin2 | 435.113 ± 49.320 (19) | 814.358 ± 39.593 (22) | | | | | | |
| Bin3 | 583.609 ± 60.060 (19) | 1040.342 ± 51.430 (22) | | | | | | |
| Bin4 | 748.473 ± 69.771 (19) | 1312.701 ± 63.141 (22) | | | | | | |
| Bin5 | 1105.538 ± 89.345 (19) | 1889.354 ± 84.390 (22) | | | | | | |
| **Syx-1A** | | | | | | | | |
| Bin1 | 525.350 ± 37.825 (19) | 1037.027 ± 94.198 (22) | | | | | | |
| Bin2 | 576.731 ± 29.333 (19) | 1202.446 ± 103.949 (22) | | | | | | |
| Bin3 | 642.284 ± 40.114 (19) | 1366.013 ± 132.185 (22) | | | | | | |
| Bin4 | 732.650 ± 46.241 (19) | 1363.597 ± 123.058 (22) | | | | | | |
| Bin5 | 795.836 ± 46.768 (19) | 1602.309 ± 156.237 (22) | | | | | | |
| **Pearson's Coefficient (Supplementary Figure 1e)** | | | | | | | | |
| **BRP/Unc18** | 0.3069 ± 0.023 (14) | 0.602 ± 0.016 (15) | < 0.0001*** | U = 0 | < 0.0001*** | t=10.48 df=27 | 0.4942 | 0.4498 |

| Parameter<br>(Figure) | control treatment (n)<br>(n = number of NMJs from at least 6 animals) | PhTx treatment (n) | Mann-Whitney U-test (P values)<br>two-tailed | test statistic | Unpaired t test (P values)<br>two-tailed | test statistic | D'Agostino & Pearson omnibus normality test<br>control treatment | PhTx treatment |
|---|---|---|---|---|---|---|---|---|
| **Synaptic intensity (% of Ctrl)** | | | | | | | | |
| **(Figure 4a)** | | | | | | | | |
| **BRP** | | | | | | | | |
| Wild-type (replotted from Figure 1d) | 100 ± 7.142 (21) | 146.5 ± 11.60 (23) | 0.0044** | U = 122 | 0.0018** | t=3.335 df=42 | 0.0058* | 0.5625 |
| Wild-type (replotted from Figure 1d) | 100 ± 7.069 (30) | 85.98 ± 6.311 (29) | 0.1656 | U = 343 | 0.1453 | t=1.476 df=57 | 0.049* | < 0.0001* |
| *rbp*$^{Null}$ | 100 ± 7.761 (23) | 99.68 ± 6.178 (28) | 0.794 | U = 308 | 0.974 | t=0.03235 df=49 | 0.0048** | < 0.0001*** |
| *brp*$^{Null}$/+ | 100 ± 6.837 (35) | 137.5 ± 10.28 (34) | 0.0014** | U = 333 | 0.0032** | t=3.054 df=67 | 0.049* | < 0.0001* |
| *rim*$^{Null}$ | 100 ± 7.461 (29) | 133.5 ± 10.82 (30) | 0.021* | U = 284 | 0.014* | t=2.531 df=57 | < 0.0001*** | 0.2342 |
| *rim*$^{Null}$,*fife*$^{Null}$ | 100 ± 6.559 (23) | 126.2 ± 10.91 (24) | 0.0681 | U = 190 | 0.0473* | t=2.040 df=45 | 0.0271* | 0.0006* |
| liprin-α$^{Null}$ | 100 ± 10.02 (25) | 145.7 ± 13.39 (26) | 0.0042** | U = 175 | 0.0091** | t=2.718 df=49 | < 0.0001* | 0.2422 |
| *syd-1*$^{Null}$ | | | | | | | | |
| **Syx-1A** | 100 ± 9.545 (21) | 95.16 ± 6.788 (28) | 0.8925 | U = 287 | 0.6724 | t=0.4255 df=47 | 0.3757 | 0.4226 |
| *brp*$^{Null}$ | | | | | | | | |
| **(Figure 4b)** | | | | | | | | |
| **Unc13A** | | | | | | | | |
| Wild-type (replotted from Figure 1D) | 100 ± 6.549 (21) | 158.9 ± 10.27 (23) | 0.0002*** | U = 88 | < 0.0001*** | t=4.730 df=42 | < 0.0001* | 0.8957 |
| *rbp*$^{Null}$ | 100 ± 6.234 (30) | 105.4 ± 10.91 (29) | 0.8115 | U = 419 | 0.6656 | t=0.4345 df=57 | 0.6723 | < 0.0001* |
| *brp*$^{Null}$/+ | 100 ± 7.761 (23) | 80.46 ± 5.507 (28) | 0.07 | U = 227 | 0.04* | t=2.012 df=49 | 0.0079** | 0.0029** |
| *rim*$^{Null}$ | 100 ± 8.543 (35) | 140.4 ± 10.13 (34) | 0.0014** | U = 332 | 0.0032** | t=3.057 df=67 | 0.0007* | 0.0753 |
| *rim*$^{Null}$,*fife*$^{Null}$ | 100 ± 6.023 (29) | 139.8 ± 13.15 (30) | 0.056 | U = 309 | 0.008** | t=2.722 df=57 | 0.4633 | 0.2304 |
| liprin-α$^{Null}$ | 100 ± 7.143 (23) | 148.2 ± 15.65 (24) | 0.035* | U = 177 | 0.0083** | t=2.761 df=45 | 0.7522 | 0.0224* |
| *syd-1*$^{Null}$ | 100 ± 9.929 (25) | 145.7 ± 11.73 (26) | 0.001*** | U = 154 | 0.0047** | t=2.965 df=49 | < 0.0001* | 0.2276 |
| *brp*$^{Null}$ | 100 ± 6.369 (21) | 102.6 ± 5.985 (28) | 0.72 | U = 276 | 0.7682 | t=0.2965 df=47 | 0.238 | 0.5113 |
| **Synaptic intensity ((a.u.); not normalized)** | | | | | | | | |
| **(Figure 4a)** | | | | | | | | |
| **BRP** | | | | | | | | |
| Wild-type (replotted from Figure 1d) | 769.5 ± 59.19 (9) | 1563 ± 170.6 (12) | 0.0007*** | U = 122 | 0.0018** | t=3.335 df=42 | 0.0058* | 0.5625 |
| *rbp*$^{Null}$ | 618.5 ± 43.72 (30) | 531.7 ± 39.03 (29) | 0.1656 | U = 343 | 0.1453 | t=1.476 df=57 | 0.0242* | 0.3501 |
| *brp*$^{Null}$/+ | 497.8 ± 38.63 (23) | 496.2 ± 30.75 (28) | 0.794 | U = 308 | 0.974 | t=0.03235 df=49 | 0.0048** | < 0.0001*** |
| Wild-type (*brp*$^{Null}$/+ control) | 662.5 ± 41.75 (21) | 1054 ± 83.75 (24) | 0.0005*** | U = 104 | 0.0002*** | t=4.000 df=43 | 0.3477 | 0.1549 |
| *rim*$^{Null}$ | 371.6 ± 25.40 (35) | 510.9 ± 38.21 (34) | 0.0014** | U = 333 | 0.0032** | t=3.054 df=67 | 0.049* | < 0.0001* |
| *rim*$^{Null}$,*fife*$^{Null}$ | 1020 ± 76.11 (29) | 1362 ± 110.4 (30) | 0.021* | U = 284 | 0.014* | t=2.531 df=57 | < 0.0001*** | 0.2342 |
| Wild-type (*rim*$^{Null}$,*fife*$^{Null}$ control) | 979.9 ± 86.39 (22) | 1532 ± 100.3 (30) | 0.0001*** | U = 128 | 0.0002*** | t=3.981 df=50 | 0.0034** | 0.263 |
| liprin-α$^{Null}$ | 721.1 ± 47.30 (23) | 910.3 ± 78.68 (24) | 0.0681 | U = 190 | 0.0473* | t=2.040 df=45 | 0.0271* | 0.0006* |
| *syd-1*$^{Null}$ | 450.0 ± 45.09 (25) | 655.7 ± 60.24 (26) | 0.0042** | U = 175 | 0.0091** | t=2.718 df=49 | < 0.0001* | 0.2422 |
| **Syx-1A** | | | | | | | | |
| *brp*$^{Null}$ | 792.3 ± 75.62 (21) | 753.9 ± 53.78 (28) | 0.8925 | U = 287 | 0.6724 | t=0.4255 df=47 | 0.3757 | 0.4226 |
| Wild-type (*brp*$^{Null}$ control) | 648.1 ± 158 (23) | 1159 ± 168.8 (28) | 0.0002*** | U = 132 | 0.03* | t=2.173 df=49 | < 0.0001*** | 0.0001*** |
| **(Figure 4b)** | | | | | | | | |
| **Unc13A** | | | | | | | | |
| Wild-type (replotted from Figure 1d) | 407.5 ± 31.53 (9) | 1580 ± 149.9 (12) | < 0.0001*** | U = 88 | < 0.0001*** | t=4.730 df=42 | < 0.0001* | 0.8957 |
| *rbp*$^{Null}$ | 272.7 ± 17 (30) | 287.5 ± 29.76 (29) | 0.8115 | U = 419 | 0.6656 | t=0.4345 df=57 | 0.6723 | < 0.0001* |
| *brp*$^{Null}$/+ | 658.1 ± 55.08 (23) | 568.3 ± 38.90 (28) | 0.07 | U = 227 | 0.04* | t=2.012 df=49 | 0.0079** | 0.0029** |
| Wild-type (*brp*$^{Null}$/+ control) | 706.4 ± 44.68 (21) | 1068 ± 84.23 (24) | 0.0012** | U = 113 | 0.0007*** | t=3.641 df=43 | 0.2251 | 0.5199 |
| *rim*$^{Null}$ | 398.7 ± 34.06 (35) | 559.8 ± 40.38 (43) | 0.0014** | U = 332 | 0.0032** | t=3.057 df=67 | 0.0007* | 0.0753 |
| *rim*$^{Null}$,*fife*$^{Null}$ | 1053 ± 63.44 (29) | 1473 ± 138.5 (30) | 0.056 | U = 309 | 0.008** | t=2.722 df=57 | 0.4633 | 0.2304 |
| Wild-type (*rim*$^{Null}$,*fife*$^{Null}$ control) | 1170 ± 113.7 (22) | 1544 ± 106.9 (30) | 0.017* | U = 202 | 0.02* | t=2.360 df=50 | 0.0062** | 0.3182 |
| liprin-α$^{Null}$ | 450.3 ± 32.17 (23) | 667.3 ± 70.48 (24) | 0.035* | U = 177 | 0.0083** | t=2.761 df=45 | 0.7522 | 0.0224* |
| *syd-1*$^{Null}$ | 534.2 ± 53.04 (25) | 778.5 ± 62.66 (26) | 0.001*** | U = 154 | 0.0047** | t=2.965 df=49 | < 0.0001* | 0.2276 |
| *brp*$^{Null}$ | 589.1 ± 37.52 (21) | 604.6 ± 35.26 (28) | 0.72 | U = 276 | 0.7682 | t=0.2965 df=47 | 0.238 | 0.5113 |
| Wild-type (*brp*$^{Null}$ control) | 777 ± 75.80 (23) | 1194 ±109 (28) | 0.005** | U = 178 | 0.004** | t=3.006 df=49 | 0.0042** | 0.2019 |

| Parameter<br>(Figure) | control treatment (n)<br>(n = single AZ from min. 13 NMJs from min. 3 animals) | PhTx treatment (n) | Mann-Whitney U-test (P values)<br>two-tailed | test statistic | Unpaired t test (P values)<br>two-tailed | test statistic | D'Agostino & Pearson omnibus normality test<br>control treatment | <br>PhTx treatment |
|---|---|---|---|---|---|---|---|---|
| **Relative frequency of Unc13A clusters<br>(Figure 2c)** | | | | | | | | |
| 1 cluster | 0.02807 (8) | 0.007463 (2) | | | | | | |
| 2 cluster | 0.045614 (13) | 0.022388 (6) | | | | | | |
| 3 cluster | 0.14386 (41) | 0.070896 (19) | | | | | | |
| 4 cluster | 0.217544 (62) | 0.115672 (31) | | | | | | |
| 5 cluster | 0.178947 (51) | 0.160448 (43) | | | | | | |
| 6 cluster | 0.168421 (48) | 0.156716 (42) | | | | | | |
| 7 cluster | 0.087719 (25) | 0.145522 (39) | | | | | | |
| 8 cluster | 0.042105 (12) | 0.119403 (32) | | | | | | |
| 9 cluster | 0.045614 (13) | 0.08209 (22) | | | | | | |
| 10 cluster | 0.024561 (7) | 0.048507 (13) | | | | | | |
| 11 cluster | 0.010526 (3) | 0.022388 (6) | | | | | | |
| 12 cluster | 0 | 0.018657 (5) | | | | | | |
| | **(n = number of NMJs)** | | | | | | | |
| **Mean cluster number** | 5.093 ± 0.3386 (13) | 6.353 ± 0.3079 (15) | | | 0.0105* | t=2.757 df=26 | 0.4705 | 0.8034 |
| **Relative frequency of BRP clusters<br>(Figure 2d)** | **(n = single AZ from min. 12 NMJs from min. 3 animals)** | | | | | | | |
| 1 cluster | 0.007916 (3) | 0.005871 (3) | | | | | | |
| 2 cluster | 0.039578 (15) | 0.019569 (10) | | | | | | |
| 3 cluster | 0.07124 (27) | 0.035225 (18) | | | | | | |
| 4 cluster | 0.150396 (57) | 0.113503 (58) | | | | | | |
| 5 cluster | 0.163588 (62) | 0.127202 (65) | | | | | | |
| 6 cluster | 0.116095 (44) | 0.113503 (58) | | | | | | |
| 7 cluster | 0.131926 (50) | 0.150685 (77) | | | | | | |
| 8 cluster | 0.058047 (22) | 0.099804 (51) | | | | | | |
| 9 cluster | 0.084433 (32) | 0.088063 (45) | | | | | | |
| 10 cluster | 0.058047 (22) | 0.074364 (38) | | | | | | |
| 11 cluster | 0.036939 (14) | 0.058708 (30) | | | | | | |
| 12 cluster | 0.023747 (9) | 0.043053 (22) | | | | | | |
| | **(n = number of NMJs)** | | | | | | | |
| **Mean cluster number** | 6.532 ± 0.2102 (12) | 7.428 ± 0.2211 (13) | | | 0.0076** | t=2.926 df=23 | 0.3753 | 0.6579 |
| **Relative frequency of RBP clusters<br>(Figure 2e)** | **(n = single AZ from min. 13 NMJs from min. 3 animals)** | | | | | | | |
| 1 cluster | 0.057554 (16) | 0.018727 (5) | | | | | | |
| 2 cluster | 0.107914 (30) | 0.014981 (4) | | | | | | |
| 3 cluster | 0.176259 (49) | 0.116105 (31) | | | | | | |
| 4 cluster | 0.197842 (55) | 0.168539 (45) | | | | | | |
| 5 cluster | 0.169065 (47) | 0.191011 (51) | | | | | | |
| 6 cluster | 0.115108 (32) | 0.164794 (44) | | | | | | |
| 7 cluster | 0.071942 (20) | 0.138577 (37) | | | | | | |
| 8 cluster | 0.05036 (14) | 0.05618 (15) | | | | | | |
| 9 cluster | 0.035971 (10) | 0.059925 (16) | | | | | | |
| 10 cluster | 0.010791 (3) | 0.029963 (8) | | | | | | |
| 11 cluster | 0.007194 (2) | 0.029963 (8) | | | | | | |
| | **(n = number of NMJs)** | | | | | | | |
| **Mean cluster number** | 4.660 ± 0.3381 (n=13) | 5.776 ± 0.2243 (n=15) | | | 0.0091** | t=2.816 df=26 | 0.0131* | 0.1084 |

| Parameter (Figure) | control treatment (n) (n = number of NMJs from min. 6 animals) | PhTx treatment (n) | Mann-Whitney U-test (P values) two-tailed | test statistic | Unpaired t test (P values) two-tailed | test statistic | D'Agostino & Pearson omnibus normality test control treatment | PhTx treatment |
|---|---|---|---|---|---|---|---|---|
| **Synaptic intensity (% of Ctrl)** | | | | | | | | |
| **(Figure 6b)** | | | | | | | | |
| **Wild-Type** | | | | | | | | |
| BRP | 100 ± 5.412 (25) | 175.0 ± 11.38 (25) | < 0.0001*** | U = 77 | < 0.0001*** | t=5.952 df=48 | 0.0113* | 0.9712 |
| RBP | 100 ± 6.029 (25) | 149.7 ± 10.44 (25) | 0.0003*** | U = 130 | 0.0001*** | t=4.122 df=48 | 0.0488* | 0.5387 |
| Cac | 100 ± 11.99 (22) | 122.3 ± 9.20 (22) | 0.0325* | U = 151 | 0.148 | t=1.473 df=42 | 0.0006*** | 0.5758 |
| **(Figure 6b)** | | | | | | | | |
| **$unc13A^{Null}$** | | | | | | | | |
| BRP | 100 ± 7.551 (31) | 147.9 ± 12.24 (31) | 0.0001*** | U = 216 | 0.0015** | t=3.329 df=60 | 0.0401* | < 0.0001* |
| RBP | 100 ± 8.856 (31) | 115.4 ± 8.147 (31) | 0.0419* | U = 336 | 0.2057 | t=1.279 df=60 | 0.0013* | 0.0001* |
| Cac | 100 ± 9.554 (25) | 152.4 ± 15.46 (20) | 0.0121* | U = 141 | 0.0044** | t=3.004 df=43 | 0.4118 | 0.074 |
| **Synaptic intensity ((a.u.); not normalized)** | | | | | | | | |
| **(Figure 6b)** | | | | | | | | |
| **Wild-Type** | | | | | | | | |
| BRP | 500.7 ± 27.10 (25) | 876.3 ± 57.00 (25) | < 0.0001*** | U = 77 | < 0.0001*** | t=5.952 df=48 | 0.0113* | 0.9712 |
| RBP | 671.5 ± 40.49 (25) | 1005 ± 70.09 (25) | 0.0003*** | U = 130 | 0.0001*** | t=4.122 df=48 | 0.0488* | 0.5387 |
| Cac | 391.2 ± 46.92 (22) | 478.3 ± 36.00 (22) | 0.0325* | U = 151 | 0.148 | t=1.473 df=42 | 0.0006*** | 0.5758 |
| **(Figure 6b)** | | | | | | | | |
| **$unc13A^{Null}$** | | | | | | | | |
| BRP | 563.7 ± 42.57 (31) | 833.7 ± 69.01 (31) | 0.0001*** | U = 216 | 0.0015** | t=3.329 df=60 | 0.0401* | < 0.0001* |
| RBP | 900.9 ± 79.78 (31) | 1040 ± 73.39 (31) | 0.0419* | U = 336 | 0.2057 | t=1.279 df=60 | 0.0013* | 0.0001* |
| Cac | 446 ± 42.62 (25) | 679.9 ± 68.94 (20) | 0.0121* | U = 141 | 0.0044** | t=3.004 df=43 | 0.4118 | 0.074 |

| Normalized comparison | Wild-type + PhTx | $unc13A^{Null}$ + PhTx | | | | | | |
|---|---|---|---|---|---|---|---|---|
| **(Figure 6b)** | | | | | | | | |
| **BRP change** | | | | | | | | |
| Wild-type + PhTx vs. $unc13A^{Null}$ + PhTx | 175.0 ± 11.38 (25) | 147.9 ± 12.24 (31) | 0.03* | U = 256 | 0.117 | t=1.592 df=54 | 0.9712 | < 0.0001*** |
| **RBP change** | | | | | | | | |
| Wild-type + PhTx vs. $unc13A^{Null}$ + PhTx | 149.7 ± 10.44 (25) | 115.4 ± 8.147 (31) | 0.0084** | U = 229 | 0.01* | t=2.629 df=54 | 0.5387 | 0.0001*** |
| **Cac change** | | | | | | | | |
| Wild-type + PhTx vs. $unc13A^{Null}$ + PhTx | 122.3 ± 9.20 (22) | 152.4 ± 15.46 (20) | 0.21 | U = 170 | 0.09 | t=1.712 df=40 | 0.5758 | 0.074 |

| | | | | | | | | |
|---|---|---|---|---|---|---|---|---|
| **Synaptic intensity (% of Ctrl)** | | | | | | | | |
| **(Figure 6g)** | | | | | | | | |
| **Unc13A-GFP rescue** | | | | | | | | |
| BRP | 100 ± 10.68 (14) | 144.3 ± 13.91 (14) | 0.0273* | U = 50 | 0.0179* | t=2.527 df=26 | 0.3376 | 0.2847 |
| GFP | 100 ± 11.39 (14) | 141.1 ± 15.51 (14) | 0.044* | U = 54 | 0.0423* | t=2.136 df=26 | 0.0003* | 0.6019 |
| **(Figure 6g)** | | | | | | | | |
| **C-term-GFP rescue** | | | | | | | | |
| BRP | 100 ± 8.526 (11) | 101.8 ± 5.563 (13) | 0.9362 | U = 70 | 0.8605 | t=0.1778 df=22 | 0.4189 | 0.7988 |
| GFP | 100 ± 9.350 (11) | 92.42 ± 8.918 (13) | 0.3877 | U = 56 | 0.5649 | t=0.5844 df=22 | 0.0209 | 0.0297 |
| **Synaptic intensity ((a.u.); not normalized)** | | | | | | | | |
| **(Figure 6g)** | | | | | | | | |
| **Unc13A-GFP rescue** | | | | | | | | |
| BRP | 612.0 ± 65.38 (14) | 883.3 ± 85.14 (14) | 0.0273* | U = 50 | 0.0179* | t=2.527 df=26 | 0.3376 | 0.2847 |
| GFP | 766.0 ± 87.28 (14) | 1081 ± 118.8 (14) | 0.044* | U = 54 | 0.0423* | t=2.136 df=26 | 0.0003* | 0.6019 |
| **(Figure 6g)** | | | | | | | | |
| **C-term-GFP rescue** | | | | | | | | |
| BRP | 925.6 ± 78.91 (11) | 941.9 ± 51.49 (13) | 0.9362 | U = 70 | 0.8605 | t=0.1778 df=22 | 0.4189 | 0.7988 |
| GFP | 2120 ±198.2 (11) | 1959 ± 189.1 (13) | 0.3877 | U = 56 | 0.5649 | t=0.5844 df=22 | 0.0209 | 0.0297 |

| Normalized comparison | Unc13A-GFP rescue (PhTx) | C-term-GFP rescue (PhTx) | | | | | | |
|---|---|---|---|---|---|---|---|---|
| **(Figure 6g)** | | | | | | | | |
| **BRP change** | | | | | | | | |
| Unc13A-GFP rescue (PhTx) vs. C-term-GFP rescue (PhTx) | 144.3 ± 13.91 (14) | 101.8 ± 5.563 (13) | 0.06 | U = 53 | 0.01* | t=2.761 df=25 | 0.2847 | 0.7988 |
| **GFP change** | | | | | | | | |
| Unc13A-GFP rescue (PhTx) vs. C-term-GFP rescue (PhTx) | 141.1 ± 15.51 (14) | 92.42 ± 8.918 (13) | 0.01* | U = 53 | 0.01* | t=2.666 df=25 | 0.6019 | 0.0297* |

| Parameter (Figure) | control or PhTx treatment (n) (n = number of NMJs from min. 6 animals) | Ctrl or PhTx treatment (n) | One way ANOVA, followed by a Turkey's multiple comparison test (P values) | test statistic F (DFn, DFd) | | | D'Agostino & Pearson omnibus normality test | |
|---|---|---|---|---|---|---|---|---|
| **(Supplementary Figure 8b) BRP** | | | | | | | | |
| Unc13A-GFP rescue (Ctrl) vs. Unc13A-GFP rescue (PhTx) | 100 ± 10.68 (14) | 144.3 ± 13.91 (14) | 0.0374* | F (3, 48) = 4.844 | | | 0.3376 | 0.2847 |
| Unc13A-GFP rescue (Ctrl) vs. C-term-GFP rescue (Ctrl) | 100 ± 10.68 (14) | 151.2 ± 12.89 (11) | 0.0206* | F (3, 48) = 4.844 | | | 0.3376 | 0.4189 |
| Unc13A-GFP rescue (Ctrl) vs. C-term-GFP rescue (PhTx) | 100 ± 10.68 (14) | 153.9 ± 8.41 (13) | 0.0091** | F (3, 48) = 4.844 | | | 0.3376 | 0.7988 |
| Unc13A-GFP rescue (PhTx) vs. C-term-GFP rescue (Ctrl) | 144.3 ± 13.91 (14) | 151.2 ± 12.89 (11) | 0.977 | F (3, 48) = 4.844 | | | 0.2847 | 0.4189 |
| Unc13A-GFP rescue (PhTx) vs. C-term-GFP rescue (PhTx) | 144.3 ± 13.91 (14) | 153.9 ± 8.41 (13) | 0.934 | F (3, 48) = 4.844 | | | 0.2847 | 0.7988 |
| C-term-GFP rescue (Ctrl) vs. C-term-GFP rescue (PhTx) | 151.2 ± 12.89 (11) | 153.9 ± 8.41 (13) | 0.998 | F (3, 48) = 4.844 | | | 0.4189 | 0.7988 |

| Parameter (Figure) | control (n) (n = number of axons from min. 3 animals) | RNAi (n) | Mann-Whitney U-test (P values) two-tailed | test statistic | Unpaired t test (P values) two-tailed | test statistic | D'Agostino & Pearson omnibus normality test control | RNAi |
|---|---|---|---|---|---|---|---|---|
| **Unc13A-GFP spots/µm² (Supplementary Figure 5c)** | | | | | | | | |
| Ctrl vs. aplip-1-RNAi | 0.198 ± 0.046 (11) | 0.424 ± 0.028 (11) | 0.0014** | U = 14 | 0.0005*** | t=4.181 df=20 | 0.3636 | 0.3853 |
| **Mander's coefficient BRP/Unc13A-GFP (Supplementary Figure 5c)** | | | | | | | | |
| Ctrl vs. aplip-1-RNAi | 0.311 ± 0.044 (11) | 0.432 ± 0.029 (11) | 0.039* | U = 29 | 0.0357* | t=2.253 df=20 | 0.0265 | 0.3364 |
| **Unc13A-GFP spot area (µm²) (Supplementary Figure 5d)** | | | | | | | | |
| Ctrl vs. srpk79D-RNAi | 0.086 ± 0.005 (10) | 0.188 ± 0.009 (12) | < 0.0001*** | U = 0 | < 0.0001*** | t=8.685 df=20 | 0.5049 | 0.9543 |
| **Mander's coefficient BRP/Unc13A-GFP (Supplementary Figure 5d)** | | | | | | | | |
| Ctrl vs. srpk79D-RNAi | 0.487 ± 0.055 (10) | 0.819 ± 0.022 (12) | < 0.0001*** | U = 1 | < 0.0001*** | t=5.892 df=20 | 0.1914 | 0.7059 |

| Parameter | control treatment (n) | PhTx treatment (n) | Mann-Whitney U-test (P values) | test statistic | Unpaired t test (P values) | test statistic | D'Agostino & Pearson omnibus normality test | |
|---|---|---|---|---|---|---|---|---|
| (Figure) | (n = number of NMJs from min. 5 animals) | | two-tailed | | two-tailed | | control | PhTx |
| **Synaptic intensity (% of Ctrl)** | | | | | | | | |
| **(Supplementary Figure 5a)** | | | | | | | | |
| BRP | 100 ± 4.707 (9) | 145.5 ± 6.508 (14) | < 0.0001*** | U = 1 | < 0.0001*** | t=5.062 df=21 | 0.2992 | 0.3471 |
| vGlut | 100 ± 2.00 (9) | 119 ± 2.765 (14) | < 0.0001*** | U = 2.5 | < 0.0001*** | t=4.975 df=21 | 0.7411 | 0.3677 |
| **Synaptic intensity ((a.u.); not normalized)** | | | | | | | | |
| **(Supplementary Figure 5a)** | | | | | | | | |
| BRP | 25864 ± 1217 (9) | 37611 ± 1682 (14) | < 0.0001*** | U = 1 | < 0.0001*** | t=5.062 df=21 | 0.2992 | 0.3471 |
| vGlut | 1418 ± 28.36 (9) | 1688 ± 39.21 (14) | < 0.0001*** | U = 2.5 | < 0.0001*** | t=4.975 df=21 | 0.7411 | 0.3677 |
| **Synaptic intensity (% of Ctrl)** | | | | | | | | |
| **(Figure 3a)** | | | | | | | | |
| **BRP** | | | | | | | | |
| Wild-type-1 | 100 ± 8.963 (19) | 156.0 ± 11.70 (22) | 0.0003*** | U = 75 | 0.0006*** | t=3.708 df=39 | 0.2156 | 0.2301 |
| *aplip-1* $^{Null}$ | 100 ± 8.545 (20) | 88.95 ± 8.648 (14) | 0.47 | U = 119 | 0.3845 | t=0.8817 df=32 | 0.0562 | 0.0011* |
| *aplip-1* $^{ek4}$ | 100 ± 13.93 (16) | 99.31 ± 13.11 (16) | 0.9159 | U = 125 | 0.9716 | t=0.03590 df=30 | 0.0167* | 0.002* |
| Wild-type-2 | 100 ± 11.67 (17) | 134.2 ± 7.612 (17) | 0.0048** | U = 64 | 0.0197* | t=2.454 df=32 | 0.0589 | 0.7469 |
| *atg-1* $^{Null}$ | 100 ± 10.95 (11) | 160.6 ± 23.81 (14) | 0.043* | U = 40 | 0.0456* | t=2.114 df=23 | 0.2288 | 0.0141* |
| *srpk79D* $^{ATC}$ | 100 ± 13.15 (15) | 103.4 ± 13.49 (10) | 0.717 | U = 68 | 0.8652 | t=0.1717 df=23 | < 0.0001* | < 0.0001* |
| **(Figure 3b)** | | | | | | | | |
| **Unc13A** | | | | | | | | |
| Wild-type-1 | 100 ± 8.869 (19) | 140.6 ± 12.21 (22) | 0.014* | U = 116 | 0.0126* | t=2.616 df=39 | 0.3026 | 0.1937 |
| *aplip-1* $^{Null}$ | 100 ± 10.28 (20) | 83.22 ± 9.817 (14) | 0.335 | U = 112 | 0.2653 | t=1.134 df=32 | 0.3504 | 0.0002* |
| *aplip-1* $^{ek4}$ | 100 ± 14.05 (16) | 89.46 ± 11.71 (16) | 0.397 | U = 105 | 0.5686 | t=0.5764 df=30 | 0.0036* | 0.005* |
| Wild-type-2 | 100 ± 11.01 (17) | 129.7 ± 9.548 (17) | 0.014* | U = 74 | 0.0502 | t=2.035 df=32 | 0.1303 | 0.0235* |
| *atg-1* $^{Null}$ | 100 ± 10.27 (11) | 156.7 ± 19.20 (14) | 0.012* | U = 32 | 0.0244* | t=2.408 df=23 | 0.0009* | 0.0042* |
| *srpk79D* $^{ATC}$ | 100 ± 13.23 (15) | 109.1 ± 12.0 (10) | 0.394 | U = 59 | 0.6348 | t=0.4814 df=23 | 0.0023* | 0.0027* |
| **Synaptic intensity ((a.u.); not normalized)** | | | | | | | | |
| **(Figure 3a)** | | | | | | | | |
| **BRP** | | | | | | | | |
| Wild-type-1 | 769.4 ± 68.95 (19) | 1200 ± 90.05 (22) | 0.0003*** | U = 75 | 0.0006*** | t=3.708 df=39 | 0.2156 | 0.2301 |
| *aplip-1* $^{Null}$ | 830.6 ± 70.98 (20) | 738.8 ± 71.83 (14) | 0.47 | U = 119 | 0.3845 | t=0.8817 df=32 | 0.0562 | 0.0011* |
| *aplip-1* $^{ek4}$ | 819.7 ± 114.2 (16) | 814.0 ± 107.4 (16) | 0.9159 | U = 125 | 0.9716 | t=0.03590 df=30 | 0.0167* | 0.002* |
| Wild-type-2 | 607.8 ± 70.93 (17) | 815.7 ± 46.27 (17) | 0.0048** | U = 64 | 0.0197* | t=2.454 df=32 | 0.0589 | 0.7469 |
| *atg-1* $^{Null}$ | 648.7 ± 71.04 (11) | 1042 ± 154.5 (14) | 0.043* | U = 40 | 0.0456* | t=2.114 df=23 | 0.2288 | 0.0141* |
| *srpk79D* $^{ATC}$ | 631.1 ± 83.01 (15) | 652.3 ± 85.15 (10) | 0.717 | U = 68 | 0.8652 | t=0.1717 df=23 | < 0.0001* | < 0.0001* |
| **(Figure 3b)** | | | | | | | | |
| **Unc13A** | | | | | | | | |
| Wild-type-1 | 526.2 ± 46.67 (19) | 739.9 ± 64.27 (22) | 0.014* | U = 116 | 0.0126* | t=2.616 df=39 | 0.3026 | 0.1937 |
| *aplip-1* $^{Null}$ | 766.9 ± 78.85 (20) | 638.2 ± 75.29 (14) | 0.335 | U = 112 | 0.2653 | t=1.134 df=32 | 0.3504 | 0.0002* |
| *aplip-1* $^{ek4}$ | 672.6 ± 94.50 (16) | 601.7 ± 78.76 (16) | 0.397 | U = 105 | 0.5686 | t=0.5764 df=30 | 0.0036* | 0.005* |
| Wild-type-2 | 713 ± 78.52 (17) | 924.5 ± 68.08 (17) | 0.014* | U = 74 | 0.0502 | t=2.035 df=32 | 0.1303 | 0.0235* |
| *atg-1* $^{Null}$ | 669.3 ± 68.76 (11) | 1049 ± 128.5 (14) | 0.012* | U = 32 | 0.0244* | t=2.408 df=23 | 0.0009* | 0.0042* |
| *srpk79D* $^{ATC}$ | 907.8 ± 120.1 (15) | 990.7 ± 108.9 (10) | 0.394 | U = 59 | 0.6348 | t=0.4814 df=23 | 0.0023* | 0.0027* |
| **Normalized comparison** | **Wild-type + PhTx** | **srpk79DATC + PhTx** | | | | | | |
| **(Figure 5d)** | | | | | | | | |
| **BRP change** | | | | | | | | |
| Wild-type + PhTx vs. *srpk79D* $^{ATC}$ + PhTx | 134.2 ± 7.612 (17) | 103.4 ± 13.49 (10) | 0.006** | U = 22 | 0.04* | t=2.158 df=25 | 0.7469 | < 0.0001*** |
| **Unc13A change** | | | | | | | | |
| Wild-type + PhTx vs. *srpk79D* $^{ATC}$ + PhTx | 129.7 ± 9.548 (17) | 109.1 ± 12.0 (10) | 0.257 | U = 62 | 0.197 | t=1.325 df=25 | 0.0235* | 0.0027** |
| **Mean cluster number** | **(n = number of NMJs from min. 4 animals)** | | | | | | | |
| **(Figure 3c)** | | | | | | | | |
| **BRP** | | | | | | | | |
| *aplip-1* $^{ek4}$ | 2.766 ± 0.08 (17) | 2.498 ± 0.09 (15) | not determined | not determined | 0.037* | t=2.174 df=30 | 0.5622 | 0.3708 |
| *srpk79D* $^{ATC}$ | 2.393 ± 0.11 (25) | 1.831 ± 0.06 (20) | not determined | not determined | 0.0002*** | t=4.001 df=43 | 0.04815 | 0.09923 |
| **Unc13A** | | | | | | | | |
| *aplip-1* $^{ek4}$ | 2.205 ± 0.08 (17) | 1.908 ± 0.08 (15) | not determined | not determined | 0.021* | t=2.420 df=30 | 0.0018** | 0.2262 |
| *srpk79D* $^{ATC}$ | 1.929 ± 0.09 (25) | 1.411 ± 0.05 (20) | not determined | not determined | 0.0002*** | t=4.141 df=43 | 0.9032 | 0.0002*** |

| Parameter (Figure) | (n = number of NMJs from min. 6 animals) | | One way ANOVA, followed by a Turkey's multiple comparison test (P values) | test statistic F (DFn, DFd) | | | D'Agostino & Pearson omnibus normality test | |
|---|---|---|---|---|---|---|---|---|
| **Synaptic intensity ((a.u.); not normalized)** | | | | | | | | |
| **(Figure 5h)** | | | | | | | | |
| **BRP** | | | | | | | | |
| Wild-type vs. *gluRIIA*$^{Null}$ | 614.2 ± 40.07 (34) | 968.2 ± 63.72 (37) | < 0.0001*** | F (3, 126) = 17.23 | | | 0.2137 | 0.339 |
| *gluRIIA*$^{Null}$ vs *srpk79D*$^{ATC}$ | 968.2 ± 63.72 (37) | 568.3 ± 44.91 (26) | < 0.0001*** | F (3, 126) = 17.23 | | | 0.339 | 0.0003*** |
| *srpk79D*$^{ATC}$ vs. *gluRIIA*$^{Null}$;*srpk79D*$^{ATC}$ | 568.3 ± 44.91 (26) | 559.2 ± 34.41 (33) | 0.999 | F (3, 126) = 17.23 | | | 0.0003*** | 0.1639 |
| *gluRIIA*$^{Null}$ vs. *gluRIIA*$^{Null}$;*srpk79D*$^{ATC}$ | 968.2 ± 63.72 (37) | 559.2 ± 34.41 (33) | < 0.0001*** | F (3, 126) = 17.23 | | | 0.339 | 0.1639 |
| **Unc13A** | | | | | | | | |
| Wild-type vs. *gluRIIA*$^{Null}$ | 526.4 ± 32.68 (34) | 735.4 ± 49.34 (37) | 0.0021** | F (3, 126) = 4.567 | | | 0.0803 | 0.3614 |
| *gluRIIA*$^{Null}$ vs *srpk79D*$^{ATC}$ | 735.4 ± 49.34 (37) | 630.8 ± 39.02 (26) | 0.3279 | F (3, 126) = 4.567 | | | 0.3614 | 0.3526 |
| *srpk79D*$^{ATC}$ vs. *gluRIIA*$^{Null}$;*srpk79D*$^{ATC}$ | 630.8 ± 39.02 (26) | 661.2 ± 41.89 (33) | 0.9627 | F (3, 126) = 4.567 | | | 0.3526 | 0.0408* |
| *gluRIIA*$^{Null}$ vs. *gluRIIA*$^{Null}$;*srpk79D*$^{ATC}$ | 735.4 ± 49.34 (37) | 559.2 ± 34.41 (33) | 0.5723 | F (3, 126) = 4.567 | | | 0.3614 | 0.0408* |

| Normalized comparison (normalized to each control genotype) (Figure 5h) | *gluRIIA*$^{Null}$ | *gluRIIA*$^{Null}$;*srpk79D*$^{ATC}$ | Mann-Whitney U-test (P values) two-tailed | test statistic | Unpaired t test (P values) two-tailed | test statistic | D'Agostino & Pearson omnibus normality test *gluRIIA*$^{Null}$ | *gluRIIA*$^{Null}$,*srpk79D*$^{ATC}$ |
|---|---|---|---|---|---|---|---|---|
| **BRP change** | | | | | | | | |
| *gluRIIA*$^{Null}$ vs. *gluRIIA*$^{Null}$;*srpk79D*$^{ATC}$ | 157.6 ± 10.37 (37) | 98.4 ± 6.054 (33) | < 0.0001*** | U = 261 | < 0.0001*** | t=4.781 df=68 | 0.339 | 0.1639 |
| **Unc13A change** | | | | | | | | |
| *gluRIIA*$^{Null}$ vs. *gluRIIA*$^{Null}$;*srpk79D*$^{ATC}$ | 139.7 ± 9.373 (37) | 104.8 ± 6.641 (33) | 0.0038** | U = 367 | 0.0041** | t=2.968 df=68 | 0.3614 | 0.0408* |

| Parameter (Figure) | (n = number of NMJs from min. 6 animals) | | One way ANOVA, followed by a Turkey's multiple comparison test (P values) | test statistic F (DFn, DFd) | | | D'Agostino & Pearson omnibus normality test | |
|---|---|---|---|---|---|---|---|---|
| **Synaptic intensity ((a.u.); not normalized)** | | | | | | | | |
| **(Supplementary Figure 6a,b)** | | | | | | | | |
| **BRP** | | | | | | | | |
| (+DMSO/-PhTX) vs (+DMSO/+PhTx) | 543.3 ± 31.75 (27) | 1078 ± 56.23 (22) | < 0.0001*** | F (3, 96) = 17.11 | | | 0.8365 | 0.3093 |
| (+DMSO/-PhTX) vs (+LatrunculinB/+PhTx) | 543.3 ± 31.75 (27) | 670.4 ± 62.20 (29) | 0.314 | F (3, 96) = 17.11 | | | 0.8365 | 0.0014** |
| (+DMSO/-PhTX) vs (+LatrunculinB/-PhTx) | 543.3 ± 31.75 (27) | 843.2 ± 64.89 (22) | 0.0014 | F (3, 96) = 17.11 | | | 0.8365 | 0.3822 |
| (+LatrunculinB/-PhTX) vs (+LatrunculinB/+PhTx) | 843.2 ± 64.89 (22) | 670.4 ± 62.20 (29) | 0.124 | F (3, 96) = 17.11 | | | 0.3822 | 0.0014** |
| **Unc13A** | | | | | | | | |
| (+DMSO/-PhTX) vs (+DMSO/+PhTx) | 538.8 ± 36.46 (27) | 1033 ± 84.69 (22) | < 0.0001*** | F (3, 96) = 44.31 | | | 0.5341 | 0.2648 |
| (+DMSO/-PhTX) vs (+LatrunculinB/+PhTx) | 538.8 ± 36.46 (27) | 664.7 ± 55.01 (29) | 0.205 | F (3, 96) = 44.31 | | | 0.5341 | 0.1275 |
| (+DMSO/-PhTX) vs (+LatrunculinB/-PhTx) | 538.8 ± 36.46 (27) | 863.2 ± 85.18 (22) | > 0.9999 | F (3, 96) = 44.31 | | | 0.5341 | 0.2104 |
| (+LatrunculinB/-PhTX) vs (+LatrunculinB/+PhTx) | 863.2 ± 85.18 (22) | 664.7 ± 55.01 (29) | 0.249 | F (3, 96) = 44.31 | | | 0.2104 | 0.1275 |
| **Synaptic intensity (normalized to -DMSO/-PhTx)** | | | | | | | | |
| **(Supplementary Figure 6a,b)** | | | | | | | | |
| **BRP** | | | | | | | | |
| (+DMSO/-PhTX) vs (+DMSO/+PhTx) | 100 ± 5.843 (27) | 198.5 ± 10.35 (22) | < 0.0001*** | F (3, 96) = 17.11 | | | 0.8365 | 0.3093 |
| (+DMSO/-PhTX) vs (+LatrunculinB/+PhTx) | 100 ± 5.843 (27) | 117.8 ± 10.40 (29) | 0.314 | F (3, 96) = 17.11 | | | 0.8365 | 0.0014** |
| (+DMSO/-PhTX) vs (+LatrunculinB/-PhTx) | 100 ± 5.843 (27) | 155.2 ± 11.94 (22) | 0.0014 | F (3, 96) = 17.11 | | | 0.8365 | 0.3822 |
| (+LatrunculinB/-PhTX) vs (+LatrunculinB/+PhTx) | 155.2 ± 11.94 (22) | 117.8 ± 10.40 (29) | 0.124 | F (3, 96) = 17.11 | | | 0.3822 | 0.0014** |
| **Unc13A** | | | | | | | | |
| (+DMSO/-PhTX) vs (+DMSO/+PhTx) | 100 ± 6.246 (27) | 177 ± 14.41 (22) | < 0.0001*** | F (3, 96) = 44.31 | | | 0.5341 | 0.2648 |
| (+DMSO/-PhTX) vs (+LatrunculinB/+PhTx) | 100 ± 6.246 (27) | 109.4 ± 8.907 (29) | 0.205 | F (3, 96) = 44.31 | | | 0.5341 | 0.1275 |
| (+DMSO/-PhTX) vs (+LatrunculinB/-PhTx) | 100 ± 6.246 (27) | 147.9 ± 14.59 (22) | > 0.9999 | F (3, 96) = 44.31 | | | 0.5341 | 0.2104 |
| (+LatrunculinB/-PhTX) vs (+LatrunculinB/+PhTx) | 147.9 ± 14.59 (22) | 109.4 ± 8.907 (29) | 0.249 | F (3, 96) = 44.31 | | | 0.2104 | 0.1275 |

| Parameter (Figure) | (n = number of NMJs from min. 6 animals) | | One way ANOVA, followed by a Turkey's multiple comparison test (P values) | test statistic F (DFn, DFd) | | | D'Agostino & Pearson omnibus normality test | |
|---|---|---|---|---|---|---|---|---|
| **Synaptic intensity ((a.u.); not normalized)** | | | | | | | | |
| **(Supplementary Fig 7f)** | | | | | | | | |
| **BRP** | | | | | | | | |
| Wild-type vs. *gluRIIA*$^{Null}$ | 716.9 ± 45.52 (33) | 929.2 ± 55.15 (34) | 0.04* | F (3, 131) = 3.841 | | | 0.0003*** | 0.0005*** |
| *gluRIIA*$^{Null}$ vs *aplip-1*$^{ek4}$ | 929.2 ± 55.15 (34) | 678.8 ± 55.82 (31) | 0.01* | F (3, 131) = 3.841 | | | 0.0005*** | 0.0653 |
| *aplip-1*$^{ek4}$ vs. *gluRIIA*$^{Null}$;*aplip-1*$^{ek4}$ | 678.8 ± 55.82 (31) | 770.4 ± 62.67 (37) | 0.651 | F (3, 131) = 3.841 | | | 0.0653 | 0.002 |
| *gluRIIA*$^{Null}$ vs. *gluRIIA*$^{Null}$;*aplip-1*$^{ek4}$ | 929.2 ± 55.15 (34) | 770.4 ± 62.67 (37) | 0.17 | F (3, 131) = 3.841 | | | 0.0005*** | 0.002 |
| **Unc13A** | | | | | | | | |
| Wild-type vs. *gluRIIA*$^{Null}$ | 551.9 ± 40.71(33) | 758.7 ± 44.26 (34) | 0.011* | F (3, 131) = 4.702 | | | 0.0212* | 0.0199* |
| *gluRIIA*$^{Null}$ vs *aplip-1*$^{ek4}$ | 758.7 ± 44.26 (34) | 610.4 ± 46.23 (31) | 0.1247 | F (3, 131) = 4.702 | | | 0.0199* | 0.1698 |
| *aplip-1*$^{ek4}$ vs. *gluRIIA*$^{Null}$;*aplip-1*$^{ek4}$ | 610.4 ± 46.23 (31) | 741.2 ± 51.86 (37) | 0.1963 | F (3, 131) = 4.702 | | | 0.1698 | 0.0909 |
| *gluRIIA*$^{Null}$ vs. *gluRIIA*$^{Null}$;*aplip-1*$^{ek4}$ | 758.7 ± 44.26 (34) | 741.2 ± 51.86 (37) | 0.9928 | F (3, 131) = 4.702 | | | 0.0199* | 0.0909 |

| Normalized comparison (normalized to each control genotype) (Supplementary Fig 7f) | *gluRIIA*$^{Null}$ | *gluRIIA*$^{Null}$;*aplip-1*$^{ek4}$ | Mann-Whitney U-test (P values) two-tailed | test statistic | Unpaired t test (P values) two-tailed | test statistic | D'Agostino & Pearson omnibus normality test *gluRIIA*$^{Null}$ | *gluRIIA*$^{Null}$,*aplip-1*$^{ek4}$ |
|---|---|---|---|---|---|---|---|---|
| **BRP change** | | | | | | | | |
| *gluRIIA*$^{Null}$ vs. *gluRIIA*$^{Null}$;*aplip-1*$^{ek4}$ | 129.6 ± 7.694 (34) | 113.5 ± 9.232 (37) | 0.02* | U = 429 | < 0.0001*** | t=4.781 df=68 | 0.0005 | 0.002 |
| **Unc13A change** | | | | | | | | |
| *gluRIIA*$^{Null}$ vs. *gluRIIA*$^{Null}$;*aplip-1*$^{ek4}$ | 137.5 ± 8.020 (34) | 121.4 ± 104.2 (37) | 0.155 | U = 505 | 0.175 | t=1.368 df=69 | 0.0199* | 0.0909 |

| Parameter (Figure) | control (n) (n = number of independent experiments) | mutant (n) | Mann-Whitney U-test (P values) two-tailed | test statistic | Unpaired t test (P values) two-tailed | test statistic | D'Agostino & Pearson omnibus normality test control | RNAi/mutant |
|---|---|---|---|---|---|---|---|---|
| **Short-term memory (Learning index (%)) (Supplementary Figure 9d)** | | | | | | | | |
| Wild-type vs. *aplip-1* $^{ek4}$ | 76.55 ± 2.834 (10) | 41.02 ± 4.309 (10) | < 0.0001*** | U = 2 | < 0.0001*** | t=6.889 df=18 | 0.093 | 0.7013 |
| Ok107::+ vs. Ok107::Aplip-1$^{RNAi}$ | 65.09 ± 3.591 (12) | 54.54 ± 2.895 (17) | 0.0236* | U = 51 | 0.0291* | t=2.305 df=27 | 0.9087 | 0.877 |
| MB247::+ vs. MB247::Aplip-1$^{RNAi}$ | 74.15 ± 30.070 (10) | 42.10 ± 4.632 (11) | < 0.0001*** | U = 3 | < 0.0001*** | t=5.645 df=19 | 0.5628 | 0.9253 |
| **Odor-avoidance MHC (Supplementary Figure 9e)** | | | | | | | | |
| Wild-type vs. *aplip-1* $^{ek4}$ | 86.90 ± 1.418 (9) | 88.20 ± 2.660 (9) | 0.422 | U =31 | 0.672 | t=0.4313 df=16 | 0.9949 | 0.4028 |
| Ok107::+ vs. Ok107::Aplip-1$^{RNAi}$ | 76.28 ± 3.656 (9) | 72.90 ± 3.360 (14) | 0.473 | U = 51 | 0.5167 | t=0.6596 df=21 | 0.3497 | 0.5675 |
| MB247::+ vs. MB247::Aplip-1$^{RNAi}$ | 70.02 ± 6.238 (9) | 57.40 ± 7.355 (14) | 0.219 | U = 26 | 0.2091 | t=1.309 df=16 | 0.1137 | 0.0109* |
| **Odor-avoidance 3-Oct (Supplementary Figure 9e)** | | | | | | | | |
| Wild-type vs. *aplip-1* $^{ek4}$ | 41.57 ± 5.588 (15) | 48.33 ± 2.699 (52) | 0.364 | U = 329 | 0.2518 | t=1.156 df=65 | 0.3434 | 0.5985 |
| Ok107::+ vs. Ok107::Aplip-1$^{RNAi}$ | 28.54 ± 3.078 (32) | 29.41 ± 3.287 (45) | 0.871 | U = 704 | 0.8536 | t=0.1851 df=75 | 0.2487 | 0.1207 |
| MB247::+ vs. MB247::Aplip-1$^{RNAi}$ | 24.62 ± 3.815 (25) | 30.87 ± 3.841 (33) | 0.335 | U = 350.5 | 0.2631 | t=1.131 df=56 | 0.232 | 0.0871 |

| **Short-term memory (Learning index (%)) (Figure 7c)** | | | **One way ANOVA, followed by a Turkey's multiple comparison test (P values)** | **test statistic F (DFn, DFd)** | | | **D'Agostino & Pearson omnibus normality test** | |
|---|---|---|---|---|---|---|---|---|
| Ok107::+ vs. UAS-Unc13A$^{RNAi}$::+ | 45.03 ± 2.934 (19) | 50.00 ± 3.198 (20) | 0.761 | F (3, 63) = 23.31 | | | 0.7883 | 0.2376 |
| Ok107::+ vs. Ok107::UAS-C-term-GFP | 45.03 ± 2.934 (19) | 51.39 ± 7.731 (11) | 0.714 | F (3, 63) = 23.31 | | | 0.7883 | N too small |
| Ok107::+ vs. Ok107::UAS-C-term-GFP::Unc13A$^{RNAi}$ | 45.03 ± 2.934 (19) | 11.64 ± 2.963 (17) | < 0.0001*** | F (3, 63) = 23.31 | | | 0.7883 | N too small |
| UAS-Unc13A$^{RNAi}$::+ vs. Ok107::UAS-C-term-GFP | 50.00 ± 3.198 (20) | 51.39 ± 7.731 (11) | 0.995 | F (3, 63) = 23.31 | | | 0.2376 | N too small |
| UAS-Unc13A$^{RNAi}$::+ vs. Ok107::UAS-C-term-GFP::Unc13A$^{RNAi}$ | 50.00 ± 3.198 (20) | 11.64 ± 2.963 (17) | < 0.0001*** | F (3, 63) = 23.31 | | | 0.2376 | N too small |
| Ok107::UAS-C-term-GFP vs. Ok107::UAS-C-term-GFP::Unc13A$^{RNAi}$ | 51.39 ± 7.731 (11) | 11.64 ± 2.963 (17) | < 0.0001*** | F (3, 63) = 23.31 | | | N too small | N too small |
| **Odor-avoidance MHC (Supplementary Figure 9b)** | | | | | | | | |
| Ok107::+ vs. UAS-Unc13A$^{RNAi}$::+ | 60.91 ± 4.522 (15) | 53.61 ± 5.793 (16) | 0.73 | F (3, 57) = 2.621 | | | N too small | N too small |
| Ok107::+ vs. Ok107::UAS-C-term-GFP | 60.91 ± 4.522 (15) | 72.19 ± 4.080 (14) | 0.419 | F (3, 57) = 2.621 | | | N too small | N too small |
| Ok107::+ vs. Ok107::UAS-C-term-GFP::Unc13A$^{RNAi}$ | 60.91 ± 4.522 (15) | 55.68 ± 5.266 (16) | 0.88 | F (3, 57) = 2.621 | | | N too small | N too small |
| UAS-Unc13A$^{RNAi}$::+ vs. Ok107::UAS-C-term-GFP | 53.61 ± 5.793 (16) | 72.19 ± 4.080 (14) | 0.058 | F (3, 57) = 2.621 | | | N too small | N too small |
| UAS-Unc13A$^{RNAi}$::+ vs. Ok107::UAS-C-term-GFP::Unc13A$^{RNAi}$ | 53.61 ± 5.793 (16) | 55.68 ± 5.266 (16) | 0.99 | F (3, 57) = 2.621 | | | N too small | N too small |
| Ok107::UAS-C-term-GFP vs. Ok107::UAS-C-term-GFP::Unc13A$^{RNAi}$ | 72.19 ± 4.080 (14) | 55.68 ± 5.266 (16) | 0.111 | F (3, 57) = 2.621 | | | N too small | N too small |
| **Odor-avoidance 3-Oct (Supplementary Figure 9b)** | | | | | | | | |
| Ok107::+ vs. UAS-Unc13A$^{RNAi}$::+ | 48.03 ± 4.372 (16) | 39.01 ± 4.392 (14) | 0.55 | F (3, 56) = 1.263 | | | 0.9924 | N too small |
| Ok107::+ vs. Ok107::UAS-C-term-GFP | 48.03 ± 4.372 (16) | 48.3 ± 6.143 (14) | >0.999 | F (3, 56) = 1.263 | | | 0.9924 | N too small |
| Ok107::+ vs. Ok107::UAS-C-term-GFP::Unc13A$^{RNAi}$ | 48.03 ± 4.372 (16) | 38.65 ± 4.269 (16) | 0.488 | F (3, 56) = 1.263 | | | 0.9924 | N too small |
| UAS-Unc13A$^{RNAi}$::+ vs. Ok107::UAS-C-term-GFP | 39.01 ± 4.392 (14) | 48.3 ± 6.143 (14) | 0.552 | F (3, 56) = 1.263 | | | N too small | N too small |
| UAS-Unc13A$^{RNAi}$::+ vs. Ok107::UAS-C-term-GFP::Unc13A$^{RNAi}$ | 39.01 ± 4.392 (14) | 38.65 ± 4.269 (16) | >0.999 | F (3, 56) = 1.263 | | | N too small | N too small |
| Ok107::UAS-C-term-GFP vs. Ok107::UAS-C-term-GFP::Unc13A$^{RNAi}$ | 48.3 ± 6.143 (14) | 38.65 ± 4.269 (16) | 0.493 | F (3, 56) = 1.263 | | | N too small | N too small |

| **Short-term memory (Learning index (%)) (Supplementary Figure 9c)** | | | **One way ANOVA, followed by a Turkey's multiple comparison test (P values)** | **test statistic F (DFn, DFd)** | | | **D'Agostino & Pearson omnibus normality test Ctrl** | **Ctrl/RNAi** |
|---|---|---|---|---|---|---|---|---|
| Ok107::+ vs. UAS-Unc13A$^{RNAi}$::+ | 58.23 ± 2.866 (11) | 62.42 ± 2.565 (12) | 0.5187 | F (2, 38) = 107.2 | | | 0.8959 | 0.7033 |
| Ok107::+ vs. Ok107::UAS-Unc13A$^{RNAi}$ | 58.23 ± 2.866 (11) | 18.55 ± 2.126 (18) | < 0.0001*** | F (2, 38) = 107.2 | | | 0.8959 | 0.2177 |
| UAS-Unc13A$^{RNAi}$::+ vs. Ok107::UAS-Unc13A-RNAi | 62.42 ± 2.565 (12) | 18.55 ± 2.126 (18) | < 0.0001*** | F (2, 38) = 107.2 | | | 0.7033 | 0.2177 |
| **Odor-avoidance MHC (Supplementary Figure 9b)** | | | | | | | | |
| Ok107::+ vs. UAS-Unc13A$^{RNAi}$::+ | 39.76 ± 4.328 (12) | 33.58 ± 4.322 (12) | 0.593 | F (2, 33) = 1.168 | | | 0.1116 | 0.0197 |
| Ok107::+ vs. Ok107::Unc13A$^{RNAi}$ | 39.76 ± 4.328 (12) | 30.29 ± 4.674 (12) | 0.301 | F (2, 33) = 1.168 | | | 0.1116 | 0.3661 |
| UAS-Unc13A$^{RNAi}$::+ vs. Ok107::UAS-Unc13A-RNAi | 33.58 ± 4.322 (12) | 30.29 ± 4.674 (12) | 0.8607 | F (2, 33) = 1.168 | | | 0.0197 | 0.3661 |
| **Odor-avoidance 3-Oct (Supplementary Figure 9b)** | | | | | | | | |
| Ok107::+ vs. UAS-Unc13A$^{RNAi}$::+ | 29.76 ± 5.67 (12) | 28.69 ± 4.921 (12) | 0.986 | F (2, 33) = 0.3417 | | | 0.5197 | 0.5839 |
| Ok107::+ vs. Ok107::Unc13A$^{RNAi}$ | 29.76 ± 5.67 (12) | 34.03 ± 3.698 (12) | 0.808 | F (2, 33) = 0.3417 | | | 0.5197 | 0.0908 |
| UAS-Unc13A$^{RNAi}$::+ vs. Ok107::UAS-Unc13A-RNAi | 28.69 ± 4.921 (12) | 34.03 ± 3.698 (12) | 0.716 | F (2, 33) = 0.3417 | | | 0.5839 | 0.0908 |

| Parameter<br>(Figure) | Wild-type (n)<br>(n = single AZ from min. 16 NMJs from min. 3 animals) | *gluRIIA* $^{Null}$ (n) | Mann-Whitney U-test (P values)<br>two-tailed | test statistic | Unpaired t test (P values)<br>two-tailed | test statistic | D'Agostino & Pearson omnibus normality test<br>Wild-type | <br>*gluRIIA* $^{Null}$ |
|---|---|---|---|---|---|---|---|---|
| **Relative frequency of Unc13A clusters (Supplementary Figure 2a)** | | | | | | | | |
| 1 cluster | 0.005725191 (3) | 0.006521739 (3) | | | | | | |
| 2 cluster | 0.02480916 (13) | 0.01304348 (6) | | | | | | |
| 3 cluster | 0.05534351 (29) | 0.02608696 (12) | | | | | | |
| 4 cluster | 0.08015267 (42) | 0.06521739 (30) | | | | | | |
| 5 cluster | 0.1164122 (61) | 0.1043478 (48) | | | | | | |
| 6 cluster | 0.1221374 (64) | 0.1304348 (60) | | | | | | |
| 7 cluster | 0.1450382 (76) | 0.126087 (58) | | | | | | |
| 8 cluster | 0.09351145 (49) | 0.09782609 (45) | | | | | | |
| 9 cluster | 0.09923664 (52) | 0.07391305 (34) | | | | | | |
| 10 cluster | 0.08396947 (44) | 0.09130435 (42) | | | | | | |
| 11 cluster | 0.04007633 (21) | 0.05652174 (26) | | | | | | |
| 12 cluster | 0.04961832 (26) | 0.03043478 (14) | | | | | | |
| | **(n = number of NMJs)** | | | | | | | |
| **Mean cluster number (n = NMJ-wise)** | 7.686 ± 0.1928 (16) | 8.408 ± 0.2596 (21) | | | 0.0421* | t=2.109 df=35 | 0.3251 | 0.5923 |
| **Relative frequency of BRP clusters** | | | | | | | | |
| | **(n = single AZ from min. 16 NMJs from min. 3 animals)** | | | | | | | |
| 1 cluster | 0.004587156 (2) | | | | | | | |
| 2 cluster | 0.03211009 (14) | | | | | | | |
| 3 cluster | 0.04816514 (21) | 0.02931596 (9) | | | | | | |
| 4 cluster | 0.1123853 (49) | 0.07817589 (24) | | | | | | |
| 5 cluster | 0.1651376 (72) | 0.1400651 (43) | | | | | | |
| 6 cluster | 0.190367 (83) | 0.1302932 (40) | | | | | | |
| 7 cluster | 0.1582569 (69) | 0.1661238 (51) | | | | | | |
| 8 cluster | 0.08256881 (36) | 0.1074919 (33) | | | | | | |
| 9 cluster | 0.06880734 (30) | 0.08794788 (27) | | | | | | |
| 10 cluster | 0.03211009 (14) | 0.05537459 (17) | | | | | | |
| 11 cluster | 0.04587156 (20) | 0.07166124 (22) | | | | | | |
| 12 cluster | 0.02293578 (10) | 0.0456026 (14) | | | | | | |
| | **(n = number of NMJs)** | | | | | | | |
| **Mean cluster number (n = NMJ-wise)** | 6.530 ± 0.2238, n=19 | 7.890 ± 0.3165, n=16 | | | 0.0011** | t=3.588 df=33 | 0.7664 | 0.6754 |
| **Relative frequency of RBP clusters** | | | | | | | | |
| | **(n = single AZ from min. 16 NMJs from min. 3 animals)** | | | | | | | |
| 1 cluster | 0.006880734 (3) | | | | | | | |
| 2 cluster | 0.01376147 (6) | 0.009771987 (3) | | | | | | |
| 3 cluster | 0.04357798 (19) | 0.01302932 (4) | | | | | | |
| 4 cluster | 0.08256881 (36) | 0.05863192 (18) | | | | | | |
| 5 cluster | 0.1077982 (47) | 0.08794788 (27) | | | | | | |
| 6 cluster | 0.130734 (57) | 0.08469056 (26) | | | | | | |
| 7 cluster | 0.1009174 (44) | 0.06840391 (21) | | | | | | |
| 8 cluster | 0.1192661 (52) | 0.1107492 (34) | | | | | | |
| 9 cluster | 0.1077982 (47) | 0.08469056 (26) | | | | | | |
| 10 cluster | 0.07110092 (31) | 0.09120521 (28) | | | | | | |
| 11 cluster | 0.03669725 (16) | 0.06514658 (20) | | | | | | |
| 12 cluster | 0.02293578 (10) | 0.05211726 (16) | | | | | | |
| | **(n = number of NMJs)** | | | | | | | |
| **Mean cluster number (n = NMJ-wise)** | 8.050 ± 0.4255, n=19 | 10.60 ± 0.6310, n=16 | | | 0.0016** | t=3.438 df=33 | 0.751 | 0.7439 |

| Parameter (Figure) | control (n) (n = single AZ from min. 10 NMJs from min. 3 animals) | PhTx (n) | Mann-Whitney U-test (P values) two-tailed | test statistic | Unpaired t test (P values) two-tailed | test statistic | D'Agostino & Pearson omnibus normality test control | PhTx |
|---|---|---|---|---|---|---|---|---|
| **Average distance of Unc13A clusters from active zone center (nm)** | | | | | | | | |
| **(Supplementary Figure 2b)** | **mean ± SEM** | **mean ± SEM** | | | | | | |
| 1 cluster | | | | | | | | |
| 2 cluster | 50.04255 ±2.767318 (13) | 46.32 ±3.659454 (6) | | | | | | |
| 3 cluster | 58.50981 ±1.626559 (41) | 61.21099 ±4.202634 (19) | | | | | | |
| 4 cluster | 69.31729 ±1.768302 (62) | 73.37585 ±2.864374 (31) | | | | | | |
| 5 cluster | 77.41753 ±1.644942 (51) | 79.00563 ±1.947779 (43) | | | | | | |
| 6 cluster | 85.90601 ±1.866263 (48) | 87.09742±1.86461 (42) | | | | | | |
| 7 cluster | 91.71197 ±2.401973 (25) | 96.54527 ±2.184172 (39) | | | | | | |
| 8 cluster | 105.5407 ±4.687239 (12) | 100.5765 ±1.613575 (32) | | | | | | |
| 9 cluster | 110.2033 ±3.11011 (13) | 107.1862 ±2.334373 (22) | | | | | | |
| 10 cluster | 122.5422 ±8.202804 (7) | 117.4547 ±3.490544 (13) | | | | | | |
| 11 cluster | 116.0642 ±5.732338 (3) | 127.1289 ±2.450859 (6) | | | | | | |
| 12 cluster | | 123.4093 ±5.022826 (5) | | | | | | |
| | **(n = number of NMJs)** | | | | | | | |
| **Change in cluster intensity** | 0.6369 ± 0.1701 (10) | 0.5853 ± 0.2812 (11) | 0.7045 | U = 49 | 0.8799 | t=0.1531 df=19 | 0.3957 | 0.001** |
| **Average distance of BRP clusters from active zone center (nm)** | **(n = single AZ from min. 11 NMJs from min. 3 animals)** | | | | | | | |
| 1 cluster | | | | | | | | |
| 2 cluster | 67.02933 ± 5.403277 (15) | 76.97274 ± 4.803899 (10) | | | | | | |
| 3 cluster | 70.36249 ± 2.927582 (27) | 73.56786 ± 3.659123 (18) | | | | | | |
| 4 cluster | 83.56571 ± 1.570682 (57) | 86.9743 ± 1.88956 (58) | | | | | | |
| 5 cluster | 87.71714 ± 1.308983 (62) | 90.52964 ± 1.937795 (65) | | | | | | |
| 6 cluster | 94.14333 ± 2.105772 (44) | 102.4955 ± 1.581542 (58) | | | | | | |
| 7 cluster | 111.2629 ± 2.039094 (50) | 109.8833 ± 1.998917 (77) | | | | | | |
| 8 cluster | 114.6796 ± 3.073702 (22) | 115.953 ± 2.172386 (51) | | | | | | |
| 9 cluster | 121.5185 ± 2.408235 (32) | 127.226 ± 2.28958 (45) | | | | | | |
| 10 cluster | 129.2212 ± 2.610988 (22) | 126.4502 ± 2.809251 (38) | | | | | | |
| 11 cluster | 135.5288 ± 4.742505 (14) | 133.8797 ± 2.498992 (30) | | | | | | |
| 12 cluster | 139.8351 ± 4.198617 (9) | 140.0481 ± 3.58417 (22) | | | | | | |
| | **(n = number of NMJs)** | | | | | | | |
| **Change in cluster intensity** | 0.8936 ± 0.2662 (11) | 0.9372 ± 0.1900 (11) | 0.847 | U = 57 | 0.8954 | t=0.1331 df=20 | 0.9568 | 0.0265* |
| **Average distance of RBP clusters from active zone center (nm)** | **(n = single AZ from min. 10 NMJs from min. 3 animals)** | | | | | | | |
| 1 cluster | | | | | | | | |
| 2 cluster | 51.78764 ± 30 (30) | 45.17186 ± 4.213212 (4) | | | | | | |
| 3 cluster | 55.75776 ± 49 (49) | 56.60377 ± 2.942738 (31) | | | | | | |
| 4 cluster | 64.35592 ± 55 (55) | 65.97104 ± 2.195476 (45) | | | | | | |
| 5 cluster | 74.95163 ± 47 (47) | 76.63934 ± 2.145343 (51) | | | | | | |
| 6 cluster | 81.19657 ± 32 (32) | 89.99515 ± 2.534665 (44) | | | | | | |
| 7 cluster | 86.89632 ± 20 (20) | 91.16457 ± 3.030604 (37) | | | | | | |
| 8 cluster | 92.94559 ± 14 (14) | 98.99152 ± 3.997981 (15) | | | | | | |
| 9 cluster | 95.14497 ± 10 (10) | 102.0089 ± 3.760186 (16) | | | | | | |
| 10 cluster | 100.3762 ± 3 (3) | 118.1631 ± 9.725718 (8) | | | | | | |
| 11 cluster | 101.8272 ± 2 (2) | 109.5169 ± 6.034848 (8) | | | | | | |
| 12 cluster | | | | | | | | |
| | **(n = number of NMJs)** | | | | | | | |
| **Change in cluster intensity** | 0.4977 ± 0.1293 (10) | 0.5306 ± 0.2281 (10) | 0.9118 | U = 48 | 0.9017 | t=0.1253 df=18 | 0.7009 | 0.4251 |

| Parameter<br>(Figure) | Wild-type (n)<br>(n = single AZ from min. 11 NMJs from min. 3 animals) | *gluRIIA* $^{Null}$ (n) | Mann-Whitney U-test (P values)<br>two-tailed | test statistic | Unpaired t test (P values)<br>two-tailed | test statistic | D'Agostino & Pearson omnibus normality test<br>Wild-type | <br>*gluRIIA* $^{Null}$ |
|---|---|---|---|---|---|---|---|---|
| **Average distance of Unc13A clusters from active zone center (nm)** | | | | | | | | |
| **(Supplementary Figure 2c)** | **mean ± SEM** | **mean ± SEM** | | | | | | |
| 1 cluster | | | | | | | | |
| 2 cluster | 59.41527 ± 6.890971 (13) | 69.50085 ± 9.069723 (3) | | | | | | |
| 3 cluster | 74.00562 ± 3.24745 (29) | 66.74512 ± 5.064179 (6) | | | | | | |
| 4 cluster | 74.58403 ± 1.613661 (42) | 76.61343 ± 3.554462 (12) | | | | | | |
| 5 cluster | 82.90544 ± 1.676531 (61) | 84.72668 ± 1.846411 (30) | | | | | | |
| 6 cluster | 88.87216 ± 1.705312 (64) | 88.88644 ± 1.729488 (48) | | | | | | |
| 7 cluster | 96.2793 ± 1.458856 (76) | 98.94801 ± 1.657797 (60) | | | | | | |
| 8 cluster | 105.0945 ± 2.077271 (49) | 98.16327 ± 1.472583 (58) | | | | | | |
| 9 cluster | 105.6874 ± 1.392574 (52) | 113.2974 ± 2.716558 (45) | | | | | | |
| 10 cluster | 115.9825 ± 1.653134 (44) | 114.7475 ± 1.951101 (34) | | | | | | |
| 11 cluster | 120.8966 ± 2.33358 (21) | 124.8844 ± 2.679401 (42) | | | | | | |
| 12 cluster | 128.6036 ± 2.660905 (26) | 125.8772 ± 3.599899 (26) | | | | | | |
| | **(n = number of NMJs)** | | | | | | | |
| **Change in cluster intensity** | 0.8001 ± 0.1978 (11) | 0.9306 ± 0.1587 (11) | 0.6063 | U = 52 | 0.6125 | t=0.5145 df=20 | 0.9965 | 0.4582 |
| **Average distance of BRP clusters from active zone center (nm)** | **(n = single AZ from min. 9 NMJs from min. 3 animals)** | | | | | | | |
| 1 cluster | | | | | | | | |
| 2 cluster | 70.38702 ± 3.527233 (14) | | | | | | | |
| 3 cluster | 73.18752 ± 2.329127 (21) | 81.82907 ± 3.846205 (9) | | | | | | |
| 4 cluster | 84.18296 ± 1.848546 (49) | 88.73561 ± 2.659988 (24) | | | | | | |
| 5 cluster | 92.53258 ± 1.703938 (72) | 97.16723 ± 1.76761 (43) | | | | | | |
| 6 cluster | 99.34858 ± 1.421175 (83) | 107.3792 ± 2.343863 (40) | | | | | | |
| 7 cluster | 108.8781 ± 1.872128 (69) | 107.6504 ± 1.676641 (51) | | | | | | |
| 8 cluster | 113.9288 ± 2.238248 (36) | 119.0085 ± 2.270118 (33) | | | | | | |
| 9 cluster | 120.3507 ± 2.153296 (30) | 120.3751 ± 2.483314 (27) | | | | | | |
| 10 cluster | 127.6512 ± 4.199829 (14) | 135.2184 ± 3.281767 (17) | | | | | | |
| 11 cluster | 131.1873 ± 3.370411 (20) | 139.5937 ± 3.861568 (22) | | | | | | |
| 12 cluster | 140.3929 ± 2.591664 (10) | 145.8077 ± 3.904849 (14) | | | | | | |
| | **(n = number of NMJs)** | | | | | | | |
| **Change in cluster intensity** | 1.404 ± 0.3680 (11) | 1.077 ± 0.4395 (9) | 0.8238 | U = 46 | 0.5718 | t=0.5758 df=18 | 0.0441* | 0.7773 |
| **Average distance of RBP clusters from active zone center (nm)** | **(n = single AZ from min. 10 NMJs from min. 3 animals)** | | | | | | | |
| 1 cluster | | | | | | | | |
| 2 cluster | 43.88898 ± 6.103397 (6) | 45.43383 ± 3.090357 (3) | | | | | | |
| 3 cluster | 57.36867 ± 3.283943 (19) | 58.36404 ± 9.065937 (4) | | | | | | |
| 4 cluster | 68.39753 ± 2.399475 (36) | 67.48476 ± 2.067253 (18) | | | | | | |
| 5 cluster | 73.29922 ± 1.556646 (47) | 78.63725 ± 3.764054 (27) | | | | | | |
| 6 cluster | 77.155 ± 1.817131 (57) | 80.18989 ± 2.763971 (26) | | | | | | |
| 7 cluster | 85.36455 ± 2.016044 (44) | 86.7642 ± 2.646105 (21) | | | | | | |
| 8 cluster | 90.6732 ± 1.519085 (52) | 96.27685 ± 2.458481 (34) | | | | | | |
| 9 cluster | 99.29681 ± 1.749993 (47) | 98.73542 ± 2.23987 (26) | | | | | | |
| 10 cluster | 101.5426 ± 2.611634 (31) | 103.0713 ± 2.448689 (28) | | | | | | |
| 11 cluster | 108.8944 ± 2.756421 (16) | 105.9697 ± 1.941058 (20) | | | | | | |
| 12 cluster | 111.8132 ± 3.512184 (10) | 110.1278 ± 3.19536 (16) | | | | | | |
| | **(n = number of NMJs)** | | | | | | | |
| **Change in cluster intensity** | 0.6404 ± 0.1627 (11) | 0.6577 ± 0.1104 (10) | 0.8633 | U = 52 | 0.932 | t=0.08642 df=19 | 0.8795 | 0.8188 |
